# Rationally Engineered Self-Assembling Protein Nanoparticles for Allergen-Specific Peanut Allergy Immunotherapy

**DOI:** 10.64898/2026.10.08.757764

**Authors:** Yi-Nan Zhang, Connor DesRoberts, Linling He, Jiang Zhu

**Affiliations:** Department of Integrative Structural and Computational Biology; Department of Immunology and Microbiology, The Scripps Research Institute, La Jolla, CA 92037, USA

## Abstract

Peanut allergy is a prevalent immunoglobulin E (IgE)-mediated food allergy that can cause life-threatening anaphylaxis. Current treatment options provide limited efficacy and variable durability of protection. While allergen-displaying virus-like particles (VLPs) have shown promise, their development is often constrained by limited design flexibility and manufacturing complexity. Here, we develop self-assembling protein nanoparticles (SApNPs) displaying the major allergens Ara h 2 and Ara h 6 as a modular platform for peanut allergy immunotherapy. Using the 60-mer TIP60 scaffold, we engineer nanoparticles by expanding particle size through α-helical insertion and replacing the human myosin-X segment with evolutionarily distant catfish homologs, followed by structure-guided and evolution-inspired optimization to enhance thermostability while minimizing human sequence content. Formulated with a Toll-like receptor 9 (TLR9) agonist, these allergen-displaying SApNPs confer robust protection against peanut-induced anaphylaxis in both prophylactic and therapeutic settings. Mechanistically, SApNPs exhibit prolonged lymph node retention, sustained presentation on follicular dendritic cells, and enhanced germinal center responses compared with soluble allergens, promoting allergen-specific IgG and functional T cell responses while limiting IgE escalation. Collectively, these findings establish allergen-displaying SApNPs as a rationally engineered, scalable, and modular platform for peanut allergy treatment with potential applicability to other allergic diseases.

**ONE-SENTENCE SUMMARY:** Rationally engineered allergen nanoparticles establish a modular platform for peanut allergy immunotherapy.

## INTRODUCTION

Peanut allergy is a common food allergy and a major public health concern due to its increasing prevalence and risk of life-threatening anaphylaxis (*1–4*). This immunoglobulin E (IgE)–mediated hypersensitivity typically develops in early childhood and, unlike many other pediatric food allergies, persists into adulthood in most affected individuals (*5–9*). Epidemiological studies indicate a substantial increase in prevalence over recent decades, particularly in Western countries, where peanut allergy affects approximately 1–3% of children and up to 1% of adults (*1, 10, 11*). Clinically, peanut allergy presents with a spectrum of symptoms, ranging from mild cutaneous manifestations, such as urticaria and angioedema, to severe systemic reactions, making it a leading cause of food-induced anaphylaxis and allergy-related mortality (*1, 4, 12–14*). At the molecular level, the disease is driven by a T helper 2 (Th2)–biased immune response and the production of allergen-specific IgE antibodies (*5, 6*). Among the identified peanut allergens, the 2S albumins Ara h 2 and Ara h 6 are considered the most potent because of their high IgE-binding capacity and strong association with severe clinical reactions (*15–19*). Similar to other type I hypersensitivity disorders, peanut allergy involves two distinct phases: sensitization, in which allergen-specific IgE is induced and subsequently binds to high-affinity receptors on mast cells and basophils, and elicitation, in which allergens cross-link these receptor-bound IgE molecules, triggering cellular degranulation and the release of histamine and other inflammatory mediators (*20–24*).

Given the severity and persistence of peanut allergy, substantial efforts have been directed toward developing therapies that modify the underlying immune response (*2, 25*). Allergen-specific immunotherapy (AIT) remains the only disease-modifying treatment, aiming to induce desensitization by increasing the threshold of peanut protein required to trigger allergic reactions and thereby reducing the risk of anaphylaxis following accidental exposure (*26–28*). Current AIT modalities include oral (OIT), epicutaneous (EPIT), and sublingual immunotherapy (SLIT) (*27, 29*). Among these, OIT achieves the highest rates of desensitization, as exemplified by the U.S. Food and Drug Administration (FDA) approval of Palforzia^®^ for pediatric patients aged 1–17 years. However, its reported discontinuation highlights challenges related to treatment burden, frequent adverse events, and limited real-world adoption (*30–34*). EPIT (e.g., Viaskin® Peanut) offers a favorable safety profile with high patient adherence and has shown promise in Phase 3 trials in young children, although its efficacy is lower than that of OIT (*35, 36*). While SLIT provides excellent safety and ease of administration, its efficacy is generally modest and may be insufficient to protect high-risk patients (*37, 38*). Recent studies suggest that low-dose OIT may represent a middle ground by offering clinically meaningful protection while minimizing gastrointestinal side effects (*39, 40*). To further improve outcomes, adjunctive biologic therapies are increasingly being incorporated into AIT regimens (*41, 42*). The anti-IgE monoclonal antibody omalizumab (Xolair^®^) can improve the safety and pace of AIT up-dosing by increasing baseline reaction thresholds, particularly in multi-allergic and high-risk patients (*43, 44*). Together, these advances reflect a maturing field; however, an effective therapy that combines high efficacy, durable protection, and minimal patient burden remains an unmet need.

Next-generation AIT must more precisely reprogram allergen-specific immune responses to enhance protective efficacy, reduce treatment burden, and achieve durable protection (*45, 46*). In this context, virus-like particles (VLPs) have emerged as an attractive platform for allergen presentation and delivery (*47–51*). VLPs are noninfectious, self-assembling nanostructures that mimic the size and repetitive surface geometry of viruses, enabling efficient B cell receptor (BCR) cross-linking and potent humoral responses without the risks associated with viral replication or pathogenicity (*52–56*). Specifically, VLPs derived from cucumber mosaic virus and incorporating a universal tetanus toxin–derived T cell epitope to enhance immunogenicity (CuMVtt) have been used to display the major peanut allergens Ara h 1 and Ara h 2 (*57*). When genetically fused or chemically coupled to CuMVtt, these allergen-displaying VLPs elicit high-titer, high-avidity IgG responses that outcompete IgE for allergen binding and prevent IgE-mediated activation of effector cells (*57–61*). Notably, these IgG–allergen immune complexes engage the inhibitory Fcγ receptor IIb (FcγRIIb), thereby suppressing mast cell and basophil degranulation and protecting against peanut extract challenge (*61*). Preclinical studies in peanut-sensitized mice further support the therapeutic potential of this platform (*59*): CuMVtt–Ara h 1, CuMVtt–Ara h 2, and mosaic “VLP Peanut” vaccines induce strong protective IgG responses without triggering allergic reactions, reduce gastrointestinal inflammation, and protect against systemic anaphylaxis upon challenge (*57, 59, 62*). In addition, these VLPs shift immune responses from Th2-dominated allergic profiles toward Th1-skewed responses, an effect that can be enhanced by encapsulated prokaryotic RNA acting as a Toll-like receptor 7 (TLR7) ligand (*63–65*). The VLP Peanut construct, which displays Ara h 2 on CuMVtt, has shown favorable safety and immunological profiles in recent Phase I/IIa clinical trials (*62*). Collectively, these findings position VLP-based approaches as a promising AIT strategy for peanut allergy. However, the structural and manufacturing constraints of viral capsid-based particles remain barriers to broader implementation of this strategy.

Self-assembling protein nanoparticles (SApNPs) have emerged as a versatile alternative to capsid-derived VLPs for antigen delivery (*66*). We previously established a rational vaccine design strategy that integrates antigen optimization with multivalent display on ferritin (FR) 24-mers and engineered bacterial E2p and I3-01 60-mers (*67–72*). These SApNPs recapitulate key geometric features of VLPs while offering greater flexibility in antigen presentation and manufacturability. They have been applied to vaccine development for HIV-1 (*67, 68, 71*), HCV (*69, 73*), filoviruses (*70, 74*), SARS-CoV-1/2 (*72, 75*), and influenza (*76–78*). Unlike soluble antigens, which are typically cleared within hours of administration, SApNPs exhibit prolonged lymph node retention, thereby promoting sustained immune cell engagement and robust germinal center (GC) reactions (*71, 74–77*). In this study, we extend this rational design strategy to peanut AIT. We displayed Ara h 2 and a tandem Ara h 6–Ara h 2 construct on the SApNP TIP60 (*79–81*), which is composed of 60 subunits, each containing a coiled-coil domain derived from human myosin X (*82*). These constructs yielded well-formed SApNPs in the ExpiCHO expression system. We further optimized the allergen–TIP60 fusion constructs by elongating the hMyoX coiled-coil to increase particle size and replacing the hMyoX segment with a distantly related catfish homolog to reduce human sequence content. Adjuvanted with a Toll-like receptor 9 (TLR9) agonist, these SApNPs provided enhanced prophylactic and therapeutic protection compared with monomers in mouse models of peanut-induced anaphylaxis, including a severe allergy model. Mechanistically, we evaluated lymph node trafficking and immune activation associated with different allergen formats at tissue, cellular, and subcellular resolution. Relative to monomers, SApNPs exhibited prolonged lymph node retention, sustained presentation on follicular dendritic cell (FDC) dendrites, and more robust and durable GC responses. These effects were accompanied by higher IgG titers and enhanced functional T cell responses. Using an evolution-inspired design strategy, we further increased the thermostability of the catfish-derived TIP60 scaffold through targeted mutations in its myosin-X segment. Together, these findings establish allergen-displaying SApNPs as a promising platform for peanut AIT, with potential applicability to other food and environmental allergies.

## RESULTS

### Rational design and in vitro characterization of TIP60 SApNPs presenting Ara h 2/6

Peanut allergy is primarily driven by the immunodominant 2S albumins Ara h 2 and Ara h 6 (*15–17, 19*). These seed storage proteins are highly resistant to heat and proteolysis and are major triggers of severe IgE-mediated reactions (*83–85*). During sensitization, exposure to Ara h 2/6 induces a Th2-biased response and allergen-specific IgE production, which arms mast cells and basophils (*20–24*). Upon re-exposure, multivalent IgE-binding epitopes on Ara h 2/6 cross-link surface-bound IgE, triggering rapid degranulation of effector cells and the release of inflammatory mediators such as histamine (*19, 86*) (**Fig. 1a**). Because these allergens are exceptionally potent, even microgram quantities can precipitate systemic responses. Recently, multivalent display of allergens on VLPs has emerged as a promising strategy for AIT, as demonstrated by Qβ presenting the major cat allergen Fel d 1 (*58*) and CuMVtt VLPs presenting peanut allergens (*59, 64*).

**Fig. 1.**
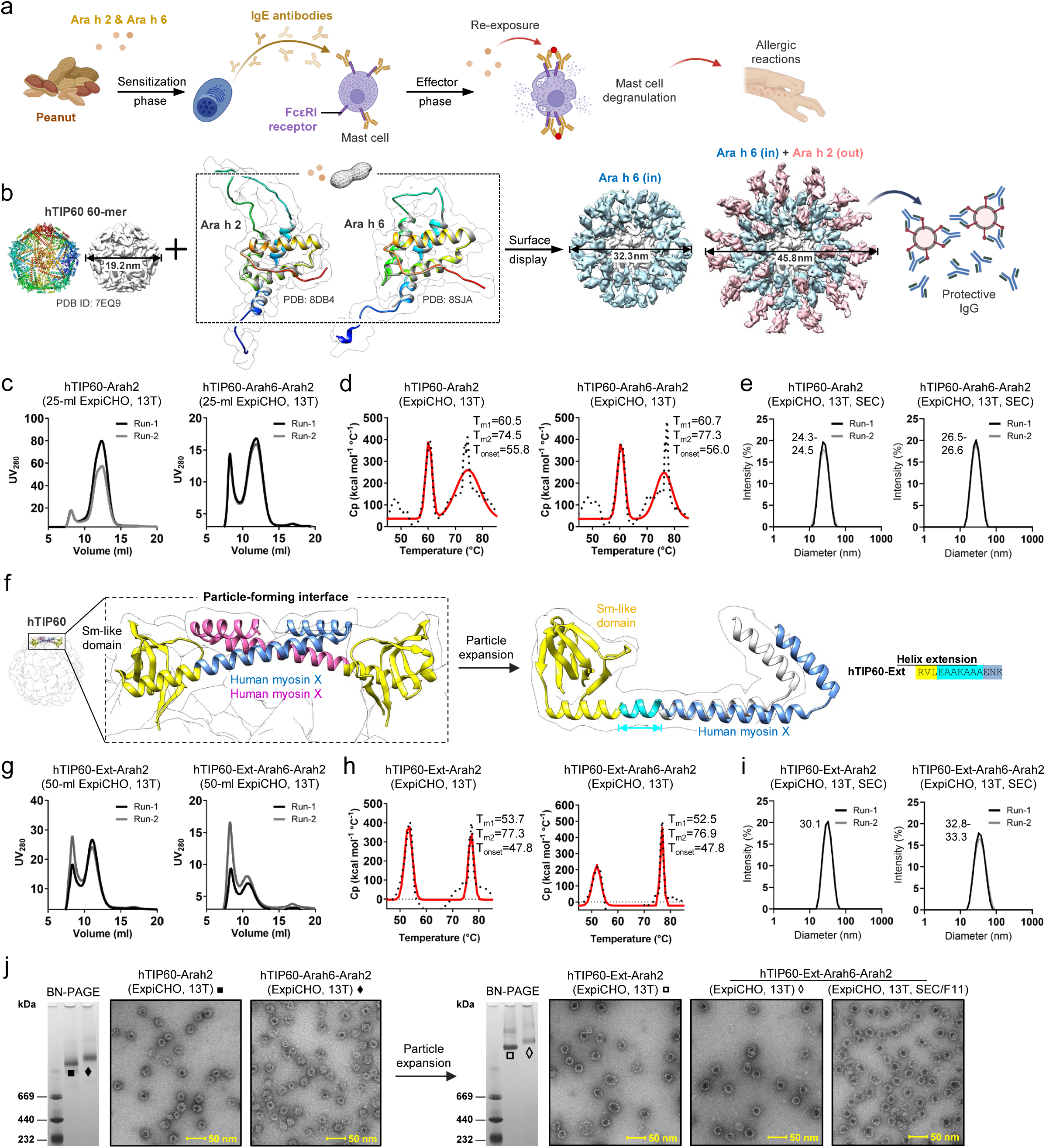
Rational design and in vitro characterization of hTIP60 nanoparticles displaying Ara h 2/6 allergens. (**a**) Schematic of the two-phase peanut allergy response. During sensitization, exposure to Ara h 2 and Ara h 6 induces allergen-specific IgE. Upon re-exposure, IgE bound to the high-affinity IgE receptor FcεRI on mast cells and basophils is cross-linked, thereby triggering degranulation and the release of inflammatory mediators that drive allergic reactions. (**b**) Structure-based design of allergen-displaying hTIP60 nanoparticles (SApNPs). Left: ribbon and surface models of hTIP60 (PDB: 7EQ9) and full-length Ara h 2 (PDB: 8DB4) and Ara h 6 (PDB: 8SJA) embedded within the molecular surface; right: surface models of hTIP60 nanoparticles displaying a tandem Arah6–Arah2 construct, with the inner layer formed by Ara h 6 (sky blue) and the outer layer by Ara h 2 (hot pink), to promote protective IgG responses. (**c**) Size-exclusion chromatography (SEC) profiles of ExpiCHO-expressed, 13T-purified hTIP60-Arah2 and hTIP60-Arah6-Arah2 from two independent productions runs. (**d**) Differential scanning calorimetry (DSC) thermograms of 13T-purified samples from one production run. Experimental data (black points) and Gaussian fits (red lines) are shown, with melting temperatures (T_m1_/T_m2_) and onset temperature (T_onset_) indicated. (**e**) Dynamic light scattering (DLS) profiles of ExpiCHO-expressed, 13T/SEC-purified hTIP60-Arah2 and hTIP60-Arah6-Arah2 from two independent production runs. Hydrodynamic diameters (Dh) are indicated. (**f**) Schematic of hTIP60 backbone expansion by insertion of an engineered 7-residue heptad repeat unit (EAAKAAA, cyan) between the Sm-like domain (yellow) and hMyoX segment (sky blue/magenta). (**g–i**) SEC **(g)**, DSC **(h)**, and DLS **(i)** characterization of expanded constructs hTIP60-Ext-Arah2 and hTIP60-Ext-Arah6-Arah2, shown as in (**c–e**). (**j**) BN-PAGE gels and negative-stain EM micrographs of hTIP60-Arah2/hTIP60-Arah6-Arah2 (left) and hTIP60-Ext-Arah2/hTIP60-Ext-Arah6-Arah2 (right). A representative nsEM micrograph of the first SEC peak of hTIP60-Ext-Arah6-Arah2 is also shown (far right). Scale bar, 50 nm.

We hypothesized that non-viral SApNPs provide a more versatile alternative to capsid-derived VLPs for peanut AIT due to their modular architecture and amenability to structure-based engineering. Previously, we utilized FR and engineered bacterial E2p and I3-01v9 scaffolds for the surface display of diverse viral glycoproteins for vaccine development (*67–78*). However, when fused to the major peanut allergens Ara h 2 and Ara h 6 (abbreviated here as Arah2 and Arah6 in construct names), all three SApNP platforms failed to express in the ExpiCHO system, suggesting that a different scaffold was required. To address this limitation, we evaluated hTIP60, a designed icosahedral nanoparticle assembled from 60 fusion proteins (*79–81*) (**Fig. 1b**, left). Each subunit comprises a pentameric Sm-like protein (*87*) and a dimeric coiled-coil domain derived from human myosin X (hMyoX) (*82*). The fully assembled hTIP60 shell has a diameter of 19.2 nm, measured at the C-terminal residue R136 (**Fig. 1b**, left). In this study, we engineered hTIP60 to display Ara h 2 and Ara h 6 through a layered fusion strategy. Ara h 6 (PDB ID: 8SJA (*88*)) was fused to the C terminus of the hTIP60 subunit via a 5GS linker to form an inner layer, followed by fusion of Ara h 2 (PDB ID: 8DB4 (*89*)) via a second 5GS linker to form the outer layer. These designs yielded modeled particle diameters of 32.3 and 45.8 nm, respectively (**Fig. 1b**, middle), suggesting that hTIP60-allergen SApNPs represent a viable alternative to allergen-displaying VLPs, such as CuMVtt-based particles (*57, 59, 62*), for peanut AIT (**Fig. 1b**, right).

Two fusion constructs, hTIP60-Arah2 and hTIP60-Arah6-Arah2, were generated to test this design concept, with a focus on Ara h 2, the dominant and more clinically important allergen (*15, 16*) (**Fig. S1a**). As demonstrated in our previous vaccine studies (*67–78*), antigen-specific antibodies can be used for tag-free purification of antigen-displaying SApNPs via immunoaffinity chromatography (IAC). In this study, monoclonal antibody 13T, isolated from a patient who achieved clinical tolerance following OIT and targeting the “bin 1” epitope of Ara h 2 (*89*), was used as the IAC ligand. Both hTIP60 constructs were transiently expressed in 25-mL ExpiCHO cultures and purified by IAC using a 13T column. Samples from two independent production runs were analyzed to confirm consistency. Size-exclusion chromatography (SEC) revealed well-defined particle peaks at ∼12.3 mL for hTIP60-Arah2 and ∼11.8 mL for hTIP60-Arah6-Arah2, with the former yielding approximately 2–3-fold more protein than the latter (**Fig. 1c**). Differential scanning calorimetry (DSC) showed distinct thermostability profiles for the two SApNPs (**Fig. 1d**). While previous indirect estimates placed the melting temperature (T_m_) of unmodified hTIP60 at ∼65–85 °C (*80*), direct DSC analysis identified an initial unfolding transition at 60.5–60.7 °C in both constructs (**Fig. 1d** and **Fig. S1b**), likely corresponding to unfolding of the hTIP60 scaffold. To assign the second thermal transition, centered at 74.5–77.4 °C, we analyzed a soluble Ara h 2 construct with a C-terminal His_6_ tag. Purified by nickel affinity chromatography, this Arah2-His_6_ construct exhibited a single unfolding event with a T_m_ of 71.1 °C (**Fig. S1c**), indicating that the higher-temperature transition in the hTIP60 constructs is attributable to the fused allergen. Finally, dynamic light scattering (DLS) analysis showed hydrodynamic diameters (Dh) of 24.3–24.5 nm for hTIP60-Arah2 and 26.5–26.6 nm for hTIP60-Arah6-Arah2 (**Fig. 1e**). The ∼2.2–2.3 nm increase in Dh is consistent with incorporation of the additional Ara h 6 layer.

A unique feature of the hTIP60 nanocage is that its edges are formed by a single helical coiled-coil spanning the C terminus of the Sm-like domain and the hMyoX segment (**Fig. 1f**, left). We reasoned that this architecture would enable precise size tuning, such that extension of the coiled-coil could proportionally increase particle dimensions without disrupting cage assembly. Because a 7-residue heptad repeat is the basic unit of an α-helical coiled-coil, insertion of such a sequence should, in principle, extend the edge by one full unit while preserving nanocage packing geometry (**Fig. 1f**, middle). To test this possibility, we inserted a single engineered heptad repeat, EAAKAAA, into the coiled-coil edge of hTIP60 (**Fig. 1f**, right). Two expanded constructs, hTIP60-Ext-Arah2 and hTIP60-Ext-Arah6-Arah2, were expressed in 50-mL ExpiCHO cultures and purified by IAC. SEC analysis showed leftward shifts of the particle peaks to ∼11.1 mL and ∼10.5 mL, respectively, although these constructs exhibited greater variability across production runs (**Fig. 1g**). As observed for the original hTIP60 constructs, addition of the Ara h 6 layer reduced SApNP yield by approximately 50%. Particle expansion significantly affected thermal stability (**Fig. 1h** and **Fig. S1d**). While the transition corresponding to allergen unfolding remained stable at 76.9–77.3 °C, the transition attributed to the expanded hTIP60 scaffold shifted downward by ∼6–8 °C to 52.5–53.7 °C. DLS measurements further supported particle expansion, showing increases in average Dh of up to ∼6 nm for hTIP60-Ext-Arah2 and ∼7 nm for hTIP60-Ext-Arah6-Arah2 (**Fig. 1i**). Finally, blue native polyacrylamide gel electrophoresis (BN-PAGE) and negative-stain electron microscopy (nsEM) confirmed the purity and structural integrity of the allergen-displaying particles following IAC purification (**Fig. 1j** and **Fig. S1e**). All four hTIP60 and hTIP60-Ext constructs showed discrete high-molecular-weight bands with no detectable species below 669 kDa. Incorporation of the Ara h 6 layer and particle expansion resulted in clear upward band shifts. nsEM images indicated that the early SEC peak (8–9 mL) corresponded to nanoparticle clusters rather than nonspecific aggregates. The hTIP60-Ext constructs exhibited increased size heterogeneity, with larger, less spherical particles enriched in early-elution fractions.

Our results establish hTIP60 as a robust SApNP platform for multivalent display of major peanut allergens. The hTIP60 scaffold enabled efficient expression, purification, and assembly of allergen-displaying nanoparticles in ExpiCHO cells while preserving favorable biophysical and structural properties. Moreover, the coiled-coil edge of hTIP60 permitted rational size expansion through the insertion of a single heptad repeat, though this modification introduced some structural heterogeneity. Collectively, these findings support hTIP60 and its derivatives as a foundation for the development of next-generation SApNP-based peanut AIT.

### Prophylactic hTIP60 SApNP vaccination protects against peanut-induced anaphylaxis

To establish a benchmark for subsequent construct optimization, we first assessed hTIP60-Arah6– Arah2 in a prophylactic mouse model of peanut-induced anaphylaxis. BALB/c mice are widely used in food allergy studies because they develop robust Th2-polarized immune responses with high IgE titers upon sensitization (*90*). Following allergen challenge, these mice consistently develop systemic anaphylaxis, characterized by reduced core body temperature and elevated clinical scores, providing quantitative readouts of AIT efficacy (*91*). This established sensitization and challenge protocol, previously used to evaluate allergen-displaying CuMVtt VLPs in peanut allergy models (*57, 59*), was adapted here to assess hTIP60-based allergen-displaying SApNPs, enabling direct comparison between the two platforms.

We first evaluated a mouse model of peanut–induced anaphylaxis. Briefly, naïve BALB/c mice were sensitized with two intraperitoneal (i.p.) injections of crude peanut extract (CPE) formulated with aluminum hydroxide (alum) at weeks 0 and 1. Sensitization doses of 50 µg or 200 µg CPE were tested. At week 3, mice were challenged i.p. with 200–2000 µg CPE, and rectal body temperature was monitored every 10 min for 60 min and at later time points up to 180 min post-challenge (**Fig. 2a,b**). Both the magnitude of temperature loss and the rate of recovery were dose–dependent. The strongest anaphylactic response was observed in mice sensitized with 200 µg CPE and challenged with 2000 µg CPE, resulting in a maximal temperature drop of 5.2 ± 0.5 °C at 40 min and recovery to baseline by 180 min. Two independent experiments produced highly consistent results, establishing this sensitization-challenge regimen as a robust benchmark protocol for subsequent evaluation of AIT efficacy.

**Fig. 2.**
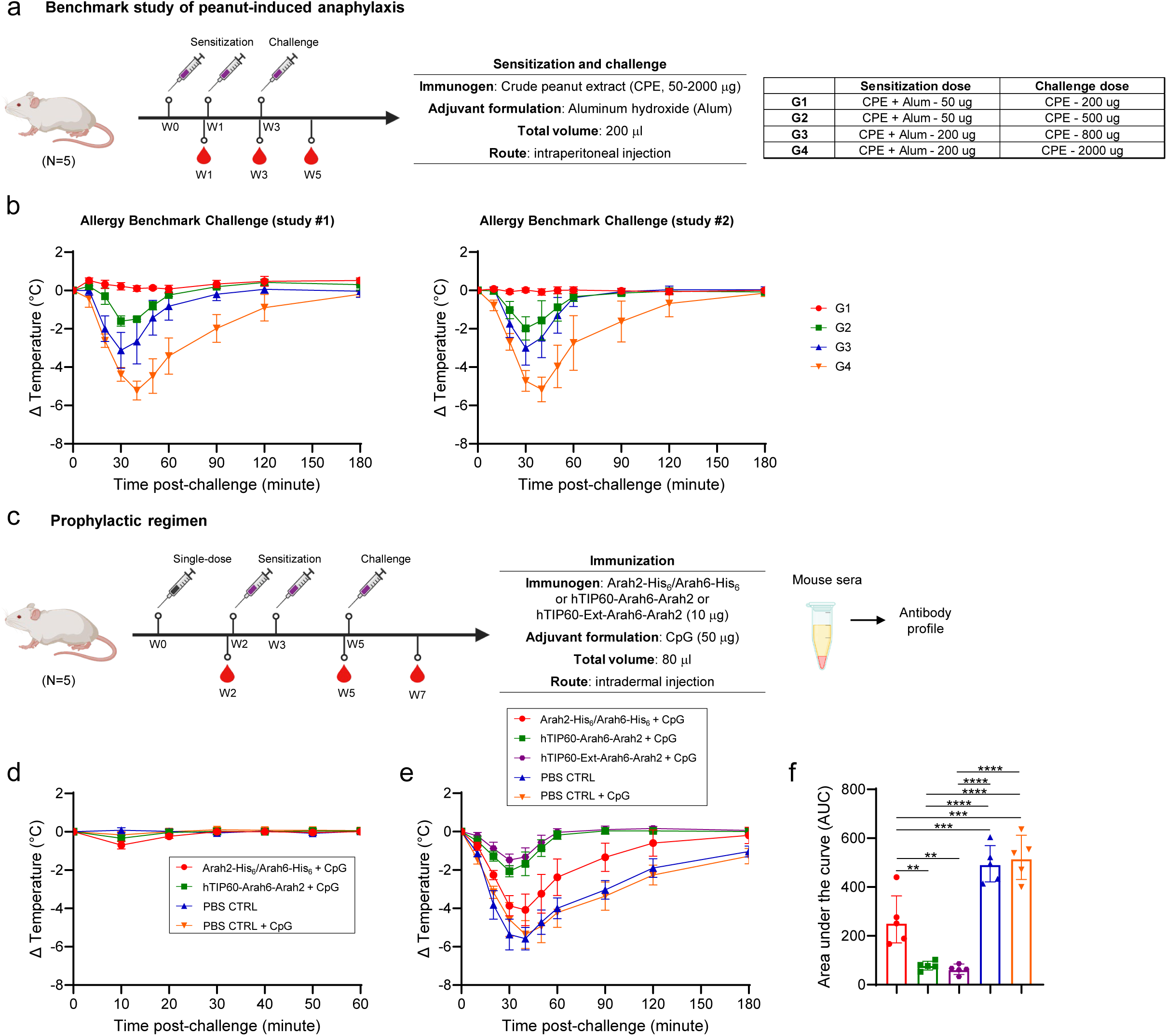
Nanoparticle vaccination protects against peanut-induced anaphylaxis. (**a**) Schematic of the mouse sensitization and challenge regimen (*n* = 5 mice per group). BALB/c mice were sensitized with crude peanut extract (CPE; 50 µg or 200 µg) formulated with aluminum hydroxide (alum) via intraperitoneal (i.p.) injection at weeks 0 and 1 and challenged at week 3 with 200– 2000 µg CPE by i.p. injection. (**b**) Rectal temperature was recorded every 10 min for 60 min and up to 180 min post-challenge. Two independent experiments established this sensitization– challenge regimen as a benchmark. (**c**) Schematic of the mouse immunization, sensitization, and challenge regimen (*n* = 5 mice per group). Mice were vaccinated intradermally with Ara h 2/6 monomers (Arah2-His_6_/Arah6-His_6_) or Ara h 2/6-displaying hTIP60 nanoparticles (SApNPs) (hTIP60-Arah6-Arah2) or hTIP60-Ext-Arah6-Arah2 SApNPs (10 µg total) formulated with CpG ODN 1826 (50 µg); PBS and CpG served as controls. (**d**) Rectal temperature was monitored for 60 min after vaccination to assess acute responses. Two weeks after vaccination, mice were sensitized and challenged with CPE (200 µg/2000 µg), and rectal temperature was monitored for 180 min. (**e**) Time-dependent curves and (**f**) area under the curve (AUC) of temperature loss over 180 min.

Using this model, we next assessed whether prophylactic vaccination could protect against peanut-induced anaphylaxis. Mice received a single intradermal dose of mixed Arah2-His_6_ and Arah6-His_6_ monomers or hTIP60-Arah6-Arah2 SApNPs or hTIP60-Ext-Arah6-Arah2 SApNPs (2.5 µg per footpad; 10 µg total), each formulated with the TLR9 agonist CpG ODN 1826 (50 µg), with PBS and CpG-alone groups as controls (**Fig. 2c**). No acute allergic reactions were observed following vaccination, as body temperature remained stable across all groups (**Fig. 2d**). Two weeks later, mice were sensitized and challenged using the established CPE protocol (200 µg sensitization; 2000 µg challenge), and body temperature was monitored for 180 min (**Fig. 2e**). Among the four groups, PBS and CpG controls exhibited severe anaphylaxis, with a temperature drop of 5.4–5.7 °C at 40 min post-challenge, followed by gradual recovery by 180 min. Vaccination with adjuvanted Arah2-His_6_/Arah6-His_6_ monomers provided partial protection, reducing temperature loss to 4.2 ± 0.7 °C, although recovery still required 120–180 min. In contrast, mice vaccinated with hTIP60-Arah6-Arah2 SApNPs showed greater protection, exhibiting a markedly smaller temperature drop of 2.1 ± 0.4 °C at 30 min and full recovery within 90 min. Notably, mice immunized with the larger SApNP construct hTIP60-Ext-Arah6-Arah2 demonstrated the strongest protection, with a temperature drop of only 1.5 ± 0.4 °C at 30 min and complete recovery within 90 min. Area-under-the-curve (AUC) analysis indicated significantly reduced temperature loss in the SApNP-vaccinated group compared with the other groups (**Fig. 2f**). These results demonstrate that prophylactic SApNP vaccination confers the most effective protection against peanut-induced anaphylaxis in this mouse model.

### Therapeutic hTIP60 SApNP treatment reduces peanut-induced anaphylaxis

Given that peanut allergy is typically diagnosed after sensitization has occurred (*5–9*), it is essential to determine whether allergen-directed treatment can provide therapeutic benefit in an established disease setting. Using the BALB/c model, mice were first sensitized with two i.p. injections of 200 µg CPE. Two weeks later, sensitized mice received a single intradermal dose of adjuvanted Arah2-His_6_/Arah6-His_6_ monomers or hTIP60-Arah6-Arah2 SApNPs or hTIP60-Ext-Arah6-Arah2 SApNPs, with PBS as the negative control (**Fig. 3a**). Body temperature remained stable after treatment in all groups, indicating no acute reactions despite prior sensitization (**Fig. 3b**). Two weeks later, all mice were challenged with 2000 µg CPE, and body temperature was monitored for 180 min (**Fig. 3c**). Mice in the PBS control group exhibited severe anaphylaxis, with a temperature drop of 4.7 ± 0.6 °C at 40 min post-challenge, followed by gradual recovery by 180 min. Mice treated with adjuvanted allergen monomers showed modest improvement, with a reduced temperature loss of 3.5 ± 0.5 °C at 40 min, although recovery still required the full 180 min. In contrast, mice treated with hTIP60- and hTIP60-Ext-allergen SApNPs showed the strongest therapeutic effect, with a markedly smaller temperature drop of 2.2 ± 0.6 °C at 30 min post-challenge and recovery to baseline within 90 min. The SApNP-treated group also exhibited a significantly lower AUC for overall temperature loss than the other groups, mirroring the robust protection observed in the prophylactic setting (**Fig. 3c,d**). Together, these findings demonstrate that hTIP60-allergen SApNP treatment confers significantly greater protection against peanut-induced anaphylaxis than soluble monomers, reducing both the severity of temperature loss and the duration of recovery in the therapeutic setting.

**Fig. 3.**
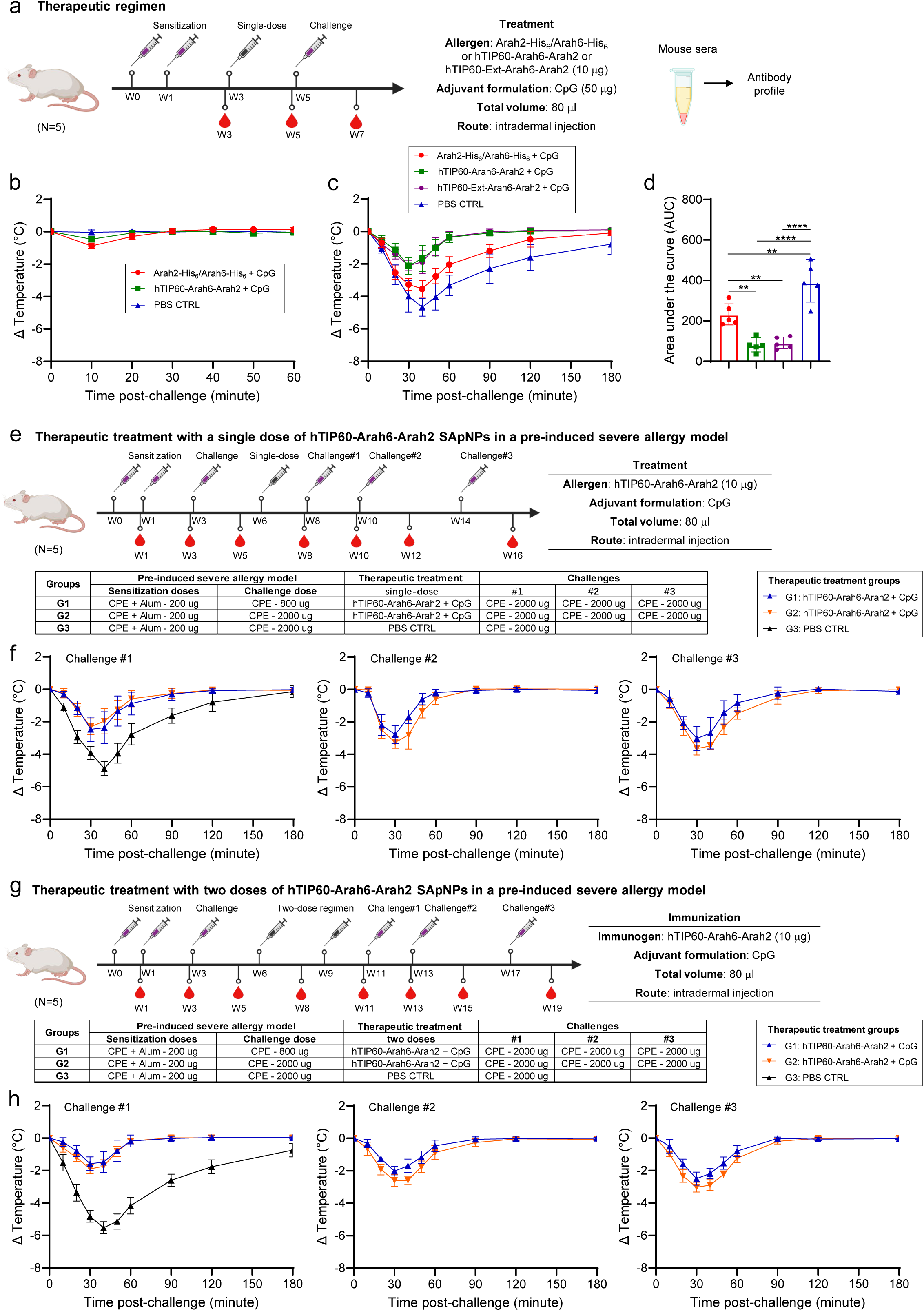
Nanoparticle treatment reduces peanut-induced anaphylaxis in established and severe models. (**a**) Schematic of the mouse sensitization, treatment, and challenge regimen (n = 5 mice per group). Therapeutic treatment following sensitization. BALB/c mice were sensitized with CPE (200 µg) via intraperitoneal (i.p.) injection at weeks 0 and 1 and injected intradermally (i.d.) two weeks later with adjuvanted Ara h 2/6 monomers (Arah2-His_6_/Arah6-His_6_) or Ara h 2/6-displaying hTIP60 nanoparticles (SApNPs) (hTIP60-Arah6-Arah2) or hTIP60-Ext-Arah6-Arah2 SApNPs (10 µg total); PBS served as a control. (**b**) Rectal temperature was recorded for 60 min after treatment to assess acute responses. Two weeks after treatment, mice were challenged with CPE (2000 µg) by i.p. injection, and rectal temperature was monitored for 180 min. (**c**) Time-dependent curves and (**d**) area under the curve (AUC) of temperature loss over 180 min. (**e**) Schematic of therapeutic protection in a pre-induced severe allergy model. Mice were sensitized with alum-adjuvanted CPE and challenged at week 3 to induce a heightened allergic state, then treated with one intradermal injection of Ara h 2/6-displaying hTIP60 nanoparticles (hTIP60-Arah6-Arah2) or PBS as a control. (**f**) Following treatment, mice were challenged three times with CPE (2000 µg) by i.p. injection, and rectal temperature was monitored for 180 min. (**g**) Schematic of a two-dose therapeutic regimen in a pre-induced severe allergy model. Mice with established severe allergy received two intradermal doses of Ara h 2/6-displaying hTIP60 nanoparticles (hTIP60-Arah6-Arah2) two weeks apart. (**h**) Two weeks after the second dose, mice were challenged three times with CPE (2000 µg) by i.p. injection, and rectal temperature was monitored for 180 min.

### Therapeutic hTIP60 SApNP treatment protects in a severe peanut allergy model

Because a subset of patients with peanut allergy exhibits severe and persistent disease, often with high sensitivity to trace allergen exposure (*1, 4, 12–14*), it is essential to evaluate therapeutic efficacy under stringent conditions that better reflect advanced clinical scenarios. In addition, such severe models have not been systematically examined in prior preclinical studies of peanut AIT. To model this heightened sensitivity, mice were sensitized with two i.p. injections of 200 µg CPE formulated with alum at weeks 0 and 1, followed by an i.p. challenge with 800 or 2000 µg CPE at week 3 to induce a heightened allergic state. Following this intensified sensitization protocol, mice were treated with hTIP60-allergen SApNPs using different regimens.

We first evaluated the therapeutic effect of a single-dose treatment. Three weeks after the disease-enhancing CPE challenge, mice received a single intradermal administration of adjuvanted hTIP60-Arah6-Arah2 SApNPs, while PBS-treated mice served as negative controls (**Fig. 3e**). Two weeks after treatment, mice underwent CPE challenge #1 (2000 µg). PBS controls exhibited severe anaphylaxis, with a temperature drop of 4.9 ± 0.4 °C at 40 min post-challenge and recovery by 180 min. In contrast, SApNP-treated mice showed markedly reduced temperature loss of 2.3–2.6 °C at 30 min and recovered within 90 min (**Fig. 3f**). CPE challenge #2, performed two weeks after CPE challenge #1, or four weeks after the treatment, resulted in a slightly greater temperature drop (2.8–3.3 °C at 30 min), but recovery again occurred within 90 min. During CPE challenge #3, conducted four weeks after challenge #2, temperature loss increased modestly (3.1–3.7 °C at 30 min), and recovery required up to 120 min. Across all three challenges, SApNP-treated mice consistently exhibited reduced hypothermia and rapid recovery despite the pre-induced severe allergic state. These findings demonstrate that even a single SApNP treatment can overcome established severe allergy and provide sustained protection against repeated peanut challenges.

We next examined whether a two-dose regimen could further enhance protection. Mice with established severe allergy received two intradermal injections of adjuvanted hTIP60-Arah6-Arah2 SApNPs, followed by three CPE challenges (**Fig. 3g**). At CPE challenge #1 (2000 µg), performed two weeks later, the two-dose group showed a smaller temperature drop (1.6–2.0 °C at 30 min) than the single-dose regimen (2.3–2.6 °C loss) and recovered to baseline within 90 min (**Fig. 3h**). PBS controls again developed severe anaphylaxis, with the greatest temperature loss at 40 min and delayed recovery by 180 min. At CPE challenge #2, performed four weeks after the two-dose treatment, temperature loss increased modestly to 2.1–2.8 °C at 30 min, but recovery remained complete within 90 min. At CPE challenge #3, conducted four weeks later, temperature loss increased slightly further to 2.6–3.1 °C at 30 min, yet the two-dose group still returned to baseline within 90 min. Across all three challenges, the two-dose regimen reduced hypothermia and accelerated recovery more effectively than the single-dose treatment, providing stronger and more durable protection in this severe disease model.

Taken together, these preclinical studies―from establishing the sensitization–challenge model to evaluating prophylactic, therapeutic, and severe disease treatment settings―demonstrate that engineered SApNPs provide an effective alternative to capsid-derived VLPs for peanut AIT and that hTIP60 represents a robust platform for further assessment and optimization.

### Rational design and in vitro characterization of catfish-derived TIP60-allergen nanoparticles

Although hTIP60 enables multivalent display of individual and tandemly linked peanut allergens and has demonstrated prophylactic and therapeutic efficacy in a murine peanut AIT model, the human-derived hMyoX segment may complicate clinical translation. An optimal SApNP scaffold for peanut AIT should minimize human-derived sequences to reduce the potential for off-target immune responses or self-recognition-associated immune interference. Reducing these “self” elements may help focus the host immune response on the displayed allergens, an important consideration for the repeated dosing regimens typical of long-term AIT. We therefore sought to replace hMyoX with distantly related non-human homologs while preserving the favorable self-assembly, structural integrity, and thermal stability of the parental TIP60 platform.

Guided by this rationale, we performed a homology search against the nonredundant (nr) database using Position-Specific Iterated BLAST (PSI-BLAST) (*92*), with the 51-residue hMyoX segment as the query. Among hits with 100% query coverage, a cluster with the lowest sequence identity (∼45–51%) was identified within the fish taxonomic group (NCBI taxid: 7898). These myosin X-like sequences were distributed across multiple branches in a phylogenetic tree rooted by hMyoX (**Fig. 4a**, left). Seven sequences representing diverse fish families were selected for initial screening of TIP60-Arah2 constructs. Only two candidates, both from the catfish lineage and designated cf1 and cf2, showed detectable expression. Sequence alignment revealed multiple mismatches relative to hMyoX, with sequence identities of 45.1% for cf1 and 47.1% for cf2 (**Fig. 4a**, right). Notably, both cf1 and cf2 contained unpaired cysteines that required mutation to avoid nonspecific disulfide formation. We next generated two pairs of TIP60 constructs displaying either Arah2 or tandem Arah6-Arah2 for in vitro characterization (**Fig. S2a**). SEC analysis revealed clear differences in assembly and yield between the two catfish scaffolds (**Fig. 4b**). A 25-mL ExpiCHO culture yielded 0.15 mg of cf1TIP60-Arah2 after IAC, representing ∼10% of the hTIP60 yield under identical conditions. In addition to a left-shifted particle peak at 11.4 mL, cf1TIP60-Arah2 lacked the prominent nanoparticle cluster peak and instead showed a visible low-molecular-weight peak at 17.1 mL, consistent with unassembled subunits (**Fig. 4b**, top). For cf1TIP60-Arah6-Arah2, yield was measurable only after scale-up to 100 mL of culture. In contrast, cf2TIP60 constructs showed substantial yields comparable to those of the parental hTIP60 platform, with well-defined particle peaks at 11.9 mL when displaying Arah2 and 11.4 mL when displaying tandem Arah6-Arah2 (**Fig. 4b**, bottom). Consistent with SEC, DSC revealed clear differences in thermal stability between the cf1 and cf2 scaffolds (**Fig. 4c** and **Fig. S2b**). cf1TIP60 exhibited a low T_m_ of 31.5–34.2 °C, below physiological temperature (37 °C), whereas cf2TIP60 displayed a higher T_m_ of 45.9–46.5 °C, although still ∼14 °C lower than hTIP60. The low yield and thermal stability of cf1TIP60 limit the suitability of this scaffold for further development. DLS showed that IAC/SEC-purified catfish-derived nanoparticles were slightly larger than their hTIP60 counterparts (**Fig. 4d**), with average diameters of ∼26 nm for Arah2-displaying constructs and 27.1–28.7 nm for tandem Arah6-Arah2-displaying constructs. This modest increase suggests that the catfish-derived TIP60 backbone adopts a slightly expanded configuration. Finally, two cf2TIP60-Ext constructs were generated for SEC, DSC, and DLS characterization (**Fig. S2a**). Particle expansion produced effects similar to those observed for hTIP60, including reduced expression yield (**Fig. S2c**), reduced thermostability (**Fig. S2d**), and increased particle size (**Fig. S2e**). Notably, cf2TIP60-Ext-Arah2 exhibited a T_m_ of 40.7 °C and a T_onset_ of 37.4 °C, indicating borderline stability at physiological temperature, which may limit the translational potential of the cf2TIP60-Ext scaffold.

**Fig. 4.**
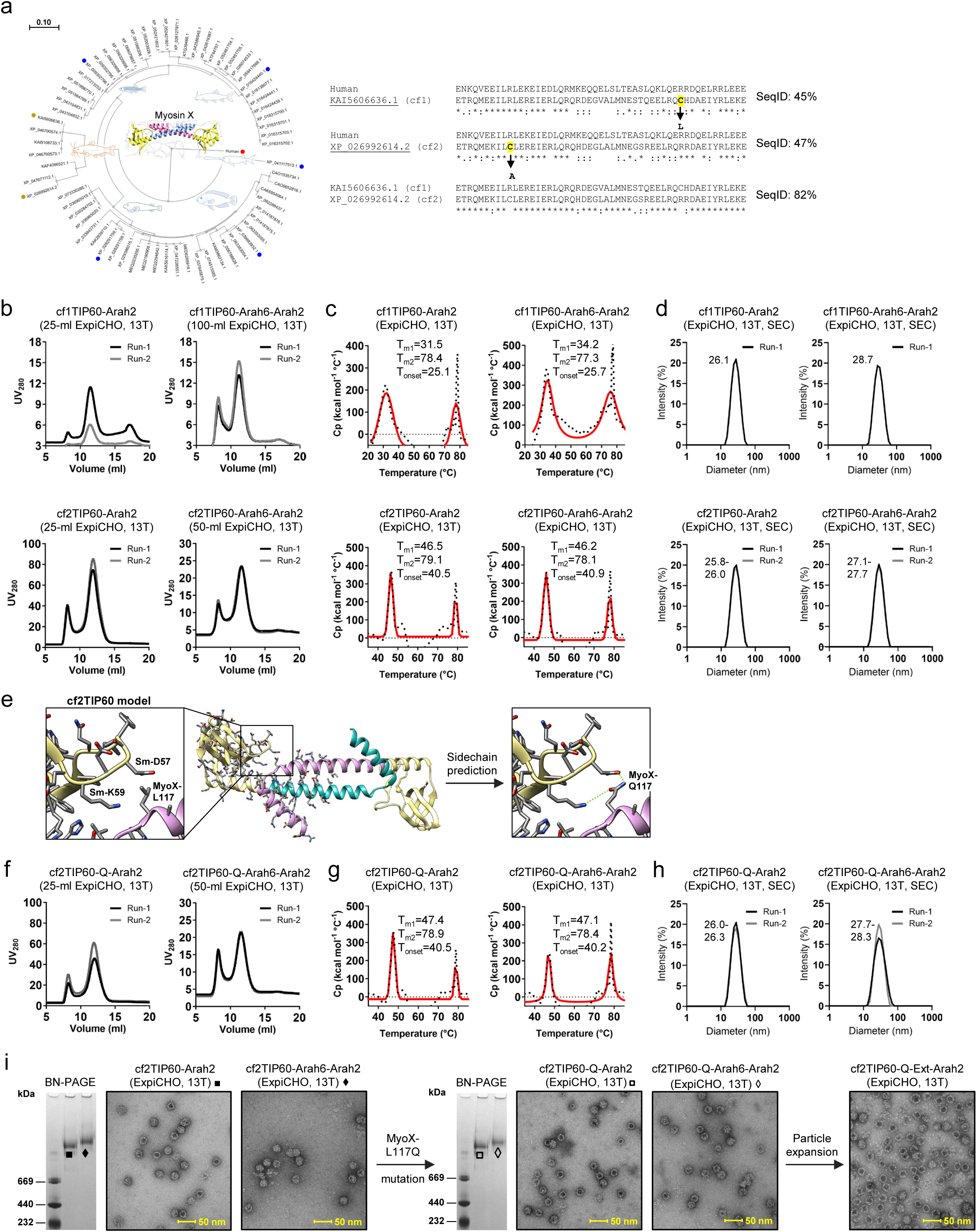
Rational design and in vitro characterization of catfish-derived TIP60 nanoparticles displaying Ara h 2/6 allergens. (**a**) Bioinformatic identification of non-human homologs of hMyoX for TIP60 design. Left: phylogenetic tree of 67 fish-derived homologs rooted by hMyoX, with each branch labeled by a schematic drawing of a fish species from the subfamily. Seven sequences representing diverse branches selected for experimental validation are indicated by colored dots on the radial phylogram (catfish: orange; other fish species: blue). Right: sequence alignments of hMyoX with cf1 (GenBank: KAI5606636.1) and cf2 (GenBank: XP_026992614.2), and between cf1 and cf2, with percent identity indicated for each sequence pair. (**b**) Size-exclusion chromatography (SEC) profiles of ExpiCHO-expressed, 13T-purified cf1/2TIP60-Arah2 and cf1/2TIP60-Arah6-Arah2 from two independent production runs. (**c**) Differential scanning calorimetry (DSC) thermograms of 13T-purified samples from one production run. Experimental data (black points) and Gaussian fits (red lines) are shown, with melting temperatures (T_m1_/T_m2_) and onset temperature (T_onset_) indicated. (**d**) Dynamic light scattering (DLS) profiles of ExpiCHO-expressed, 13T/SEC-purified cf1/2TIP60-Arah2 and cf1/2TIP60-Arah6-Arah2 from two independent production runs. Hydrodynamic diameters (Dh) are indicated. (**e**) Schematic of interactions at the interface between the Sm-like domain (D57 and K59) and cf2-MyoX residue L117, and the L117Q mutation. An enlarged view of this contact is shown as an inset (left), along with a cf2TIP60 dimer model highlighting surrounding side chains (middle) and an enlarged view of the L117Q mutation (right). (**f–h**) SEC **(f)**, DSC **(g)**, and DLS **(h)** characterization of constructs cf2TIP60-Q-Arah2 and cf2TIP60-Q-Arah6-Arah2, shown as in (**b–d**). (**i**) BN-PAGE gels and negative-stain EM micrographs of cf2TIP60-Arah2/cf2TIP60-Arah6-Arah2 (left) and cf2TIP60-Q-Arah2/cf2TIP60-Q-Arah6-Arah2 (right). A representative nsEM micrograph of cf2TIP60-Q-Ext-Arah6-Arah2 is also shown (far right). Scale bar, 50 nm.

Given its robust expression and self-assembly, cf2TIP60 provided a promising scaffold for further optimization. Homology modeling of the cf2TIP60 dimer identified a site for structural refinement: the side chain of L117 is positioned between D57 and K59 in the Sm-like domain (**Fig. 4e**, left). This configuration suggests a localized source of instability, likely driven by steric clashes and electrostatic repulsion with D57 (C_β_–C_β_ = 4.3 Å) and a weaker contact with K59 (C_β_–C_β_ = 8.5 Å). We therefore hypothesized that an L117Q mutation would not only eliminate these unfavorable interactions but also introduce stabilizing hydrogen bonds (**Fig. 4e**, right). To test this possibility, we generated two cf2TIP60 variants harboring the L117Q mutation and characterized them in vitro. SEC profiles demonstrated that the yield and composition of these variants were largely comparable to those of the parental cf2TIP60 constructs (**Fig. 4f**). DSC measurements revealed a modest but consistent increase in T_m_ (∼1 °C) relative to the parental constructs (**Fig. 4g** and **Fig. S2f**), while DLS analysis showed that particle size remained largely unaffected by the mutation (**Fig. 4h**). Finally, nsEM was used to visualize purified cf1TIP60 and several cf2TIP60 variants. Micrographs of cf1TIP60 revealed substantial impurity and structural heterogeneity, reflecting particle dissociation under ambient conditions (**Fig. S2g**). In contrast, cf2TIP60 constructs with or without the L117Q mutation yielded well-defined, homogeneous nanoparticles (**Fig. 4i** and **Fig. S2g**). However, the L117Q mutation proved insufficient to rescue the suboptimal yield and purity associated with scaffold expansion, as observed for cf2TIP60-Q-Ext-Arah2.

Our findings suggest that catfish-derived homologs may provide viable alternatives to hMyoX for the TIP60 platform. While cf1 proved unsuitable due to suboptimal yield and thermal stability, the cf2 variant, cf2TIP60, maintained an in vitro profile largely comparable to that of hTIP60. Structure-based analysis identified a hotspot at the dimer interface in cf2TIP60, where the L117Q mutation conferred a modest stabilizing effect. Together, the cf2TIP60-Q scaffold and its derivative constructs provide a robust foundation for further preclinical assessment.

### Catfish-derived TIP60 SApNPs confer protection and modulate IgG/IgE responses

Using the established peanut allergy models (**Figs. 2**, **3**), we assessed the efficacy of two catfish-derived SApNPs, cf2TIP60-Q displaying Arah2 or tandem Arah6-Arah2, against peanut-induced anaphylaxis (**Fig. 5**). In the prophylactic setting, mice received a single intradermal dose of adjuvanted SApNP (2.5 µg per footpad; 10 µg total). Two weeks later, mice were sensitized and challenged with CPE (200 µg sensitization; 2000 µg challenge), with body temperature monitored for 180 min (**Fig. 5a**). Both cf2TIP60-Q SApNPs conferred strong protection, with temperature drops of 2.1–2.6 °C at 30–40 min and full recovery within 90 min, accompanied by low AUC values and closely matching the protective profile seen with hTIP60–Arah6–Arah2 (**Fig. 2c-f**). In the therapeutic setting, sensitized mice received a single intradermal dose of adjuvanted SApNP. Two weeks later, mice were challenged with 2000 µg CPE, and body temperature was monitored for 180 min (**Fig. 5b**). Both cf2TIP60-Q SApNPs conferred therapeutic protection, with 2.6–2.7 °C temperature drops at 40 min and recovery to baseline within 120 min. This protection was slightly weaker than that observed with hTIP60 SApNPs (**Fig. 3**), suggesting that the current cf2TIP60-Q constructs may require optimization to achieve maximal therapeutic efficacy.

**Fig. 5.**
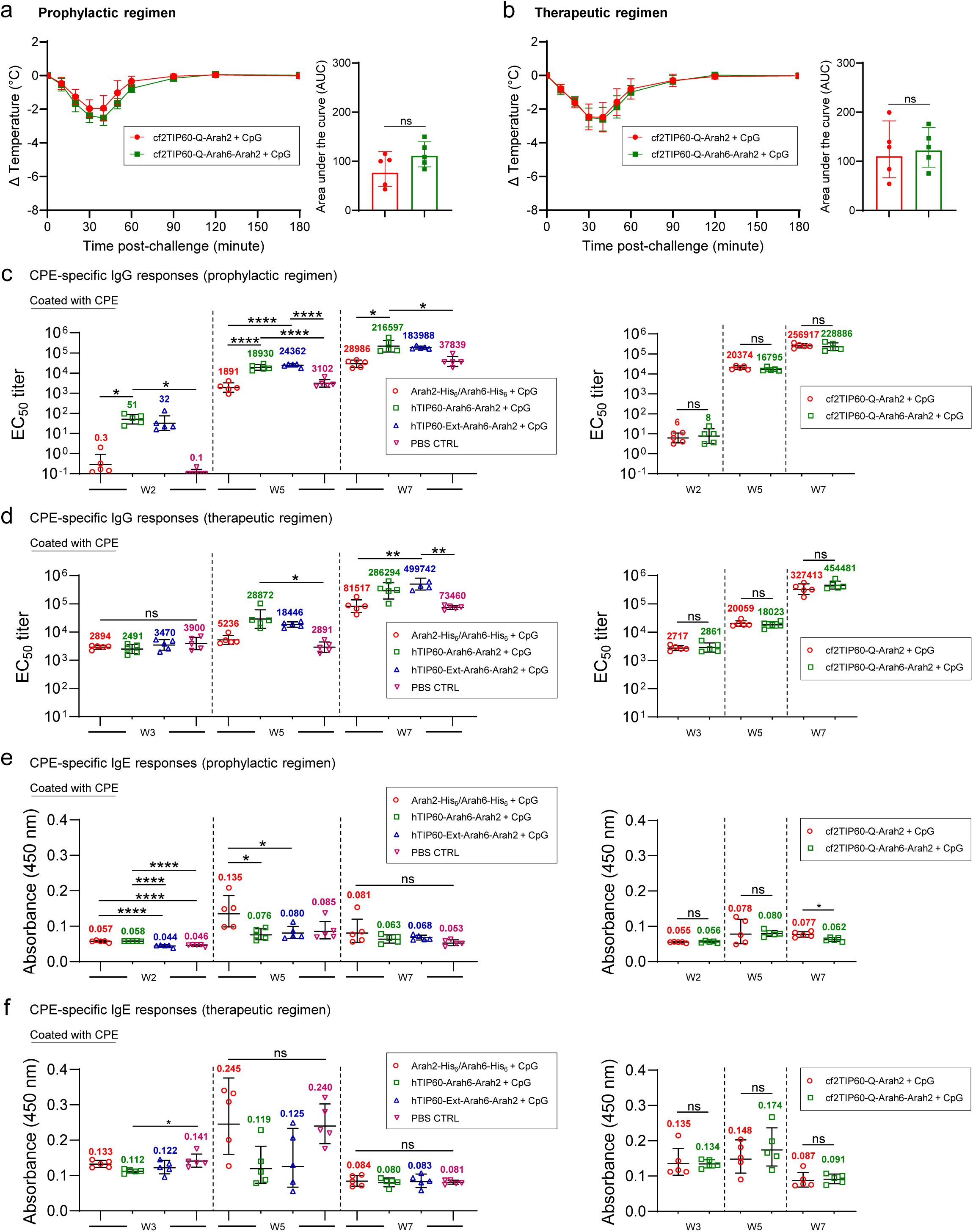
Catfish-derived TIP60 Nanoparticles protect against peanut-induced anaphylaxis in prophylactic and therapeutic peanut allergy models. **(a)** In a prophylactic peanut allergy model, mice were vaccinated intradermally with catfish–derived SApNPs, cf2TIP60–Q–Arah2 or cf2TIP60–Q–Arah6–Arah2 (10 µg total), formulated with CpG ODN 1826 (50 µg) (n = 5 per group). Two weeks later, mice were sensitized and challenged with CPE (200 µg/2000 µg), and rectal temperature was monitored for 180 min. Shown are time–dependent temperature curves and AUC values for temperature loss over 180 min. (b) In a therapeutic peanut allergy model, mice were sensitized with CPE (200 µg, i.p.) at weeks 0 and 1 and treated intradermally two weeks later with adjuvanted cf2TIP60–Q–Arah2 or cf2TIP60–Q–Arah6–Arah2 (10 µg total) (n = 5 per group). Two weeks after treatment, mice were challenged with CPE (2000 µg, i.p.), and rectal temperature was monitored for 180 min. Time–dependent temperature curves and AUC values for temperature loss over 180 min are shown. (**c-f**) Antibody responses: CPE–specific IgG titers in the (**c**) prophylactic and (**d**) therapeutic settings, and CPE–specific IgE responses in the (**e**) prophylactic and (**f**) therapeutic settings. EC_50_ titers were derived from serum ELISA against CPE. For samples with EC_50_ values below 0.1, values were set to 0.1 to maintain consistent plotting and comparison. A_450_ values are reported for IgE responses. Geometric means are indicated on the plots, and data are presented as geometric mean ± geometric SD. Statistical analysis was performed using one–way ANOVA with Tukey’s post hoc test for multiple comparisons at each time point. Two–tailed unpaired t–tests with Welch’s correction were used for comparisons between two independent groups. Statistical significance is indicated as: ns (not significant), *p < 0.05, **p < 0.01, and ****p < 0.0001.

To determine the allergen-specific antibody profile during AIT using human- and catfish-derived TIP60 SApNPs, we analyzed sera collected at weeks 2/3, 5, and 7 from the SApNP, Arah2-His_6_/Arah6-His_6_ monomer mix, and PBS control groups by enzyme-linked immunosorbent assay (ELISA) with crude peanut extract (CPE)-coated plates (**Fig. 5c-f** and **Fig. S3**). In the prophylactic setting, both adjuvanted hTIP60 SApNPs elicited the highest IgG responses at week 2 after a single dose, with EC_50_ titers 107–170–fold above the monomer group (**Fig. 5c** and **Fig. S3a,b**). At week 5, after two doses of CPE sensitization, the hTIP60 and hTIP60–Ext SApNP groups reached EC_50_ titers of 18,930 and 24,362, respectively, 10–13–fold above monomers. By week 7, two weeks after CPE challenge, both hTIP60 SApNP groups maintained the highest responses, with EC_50_ titers 6–7–fold above the monomer group. The two adjuvanted cf2TIP60 SApNPs elicited lower IgG responses at week 2 than the hTIP60 constructs, but their EC_50_ titers remained 20–27–fold above monomers. After CPE sensitization and challenge, the two cf2TIP60–Q groups exhibited IgG responses comparable to those of hTIP60 SApNPs, with EC_50_ titers 9–11–fold above monomers at week 5 and 8–9–fold at week 7. In the therapeutic setting, the two hTIP60 SApNP groups and the two controls showed high IgG responses at week 3 following two doses of CPE sensitization, with EC_50_ titers of 2,491–3,900 (**Fig. 5d** and **Fig. S3c,d**). At week 5, two weeks after treatment, the monomer group showed only a modest increase, ∼2–fold above its week-3 value. In contrast, both hTIP60 SApNP groups exhibited a substantial boost, with EC_50_ titers 4–6–fold higher than the monomer group and 6–10–fold higher than the PBS control. By week 7, two weeks after CPE challenge, these four groups showed a further increase in IgG responses, with EC_50_ titers exceeding those observed in the prophylactic setting. Both hTIP60 SApNPs maintained the highest IgG levels among the four groups, with EC_50_ titers 4–6–fold above monomers. The two cf2TIP60 SApNP groups showed a similar pattern, with EC_50_ titers comparable to those of hTIP60 SApNPs across all time points. At week 5, for example, the cf2TIP60–Q–Arah2 and cf2TIP60–Q–Arah6–Arah2 groups reached EC_50_ titers of 20,059 and 18,023, 4– and 3–fold above monomers, respectively. Together, these data show that multivalent display of single or tandem allergens on either hTIP60 or cf2TIP60-Q SApNPs can substantially enhance allergen–specific IgG responses, consistent with the protective effects observed in the prophylactic and therapeutic AIT models.

We next evaluated IgE responses during peanut AIT. In the prophylactic setting, both hTIP60 SApNP groups and the two control groups exhibited relatively low IgE responses at week 2 after a single prophylactic dose, with absorbance (A_450_) values of 0.044–0.057 (**Fig. 5e** and **Fig. S3e**). At week 5, after two doses of CPE sensitization, all four groups showed increased IgE levels, with the monomer group displaying the largest and most significant increase, reaching an A_450_ value of 0.135, 2.4–fold above its week 2 value. By week 7, two weeks after CPE challenge, IgE responses declined across all groups, with A_450_ values of 0.053–0.081, lower than those observed at week 5. Both cf2TIP60 SApNP groups similarly maintained low IgE levels across all time points, comparable to those of the two hTIP60 SApNP groups, indicating that multivalent TIP60 SApNP vaccination does not amplify IgE responses. In the therapeutic setting, both hTIP60 SApNP groups and the two control groups showed high IgE levels at week 3 following two doses of CPE sensitization, with A_450_ values of 0.112–0.141 (**Fig. 5f** and **Fig. S3f**). At week 5, two weeks after treatment, IgE levels further increased in the monomer and PBS control groups, reaching 0.240–0.245, corresponding to 1.8–fold and 1.7–fold increases, respectively, over their week 3 values. In contrast, both hTIP60 SApNP groups displayed negligible changes in their IgE levels, remaining ∼2.0-fold below the monomer group. At week 7, two weeks after CPE challenge, IgE responses declined across all groups, with A_450_ values returning to 0.080–0.084, lower than those observed at both weeks 3 and 5. The cf2TIP60 SApNP groups showed a similar pattern, although their IgE levels were slightly elevated at week 5, with A_450_ values of 0.148–0.174. Nonetheless, these values remained lower than those of the monomer and PBS control groups, suggesting that treatment with TIP60 SApNPs can limit IgE escalation.

These mouse studies confirm the prophylactic and therapeutic efficacy of catfish-derived cf2TIP60-Q for peanut AIT, with performance largely comparable to human-derived hTIP60. Longitudinal serum analysis demonstrated that allergen format influences the antibody profile generated during AIT. Whereas the monomer and PBS control groups developed heightened IgE responses during sensitization and challenge, multivalent display on hTIP60 and cf2TIP60-Q SApNPs elicited high peanut-specific IgG titers while limiting IgE escalation. This enhanced IgG response likely contributes to protection by competing with IgE for allergen binding, reducing effector-cell activation and supporting a favorable humoral environment for AIT.

### Distribution, trafficking, and retention of TIP60 SApNPs in lymph nodes

In previous vaccine studies, we investigated the in vivo behavior of soluble antigens and antigen-presenting SApNPs, including the E2p and I3-01v9 60-mers (*71, 74–77*). Using a similar approach, we characterized allergen-displaying TIP60 SApNPs to better understand their lymph node interactions and ability to promote favorable, non-allergic immune responses. To mount an effective humoral response, these SApNPs must be transported through the lymphatic system and accumulate in lymph node follicles. However, immunostaining with 13T and 15T antibodies (*89*) produced no signals in lymph node tissues, precluding direct analysis of TIP60 SApNP trafficking (**Fig. S4a**). To overcome this, we designed two surrogate constructs by displaying tandemly linked human, avian, and swine influenza M2e sequences (M2ex3) (*76*) on the hTIP60 and cf2TIP60-Q scaffolds (**Fig. S4b**). Although M2ex3 is smaller than Arah2 and Arah6-Arah2, it provides a well-defined, non-glycosylated surrogate for assessing TIP60 scaffold trafficking.

The M2ex3-displaying TIP60 constructs were first characterized in vitro. Structural modeling indicated that M2ex3-hTIP60 and M2ex3-cf2TIP60-Q, if successfully assembled, would form particles with diameters of ∼30.5 nm, measured at P10 (**Fig. 6a**, left). The two M2ex3-TIP60 fusion constructs were transiently expressed in 25-mL ExpiCHO cultures and purified by IAC using an mAb148 antibody column (*76*). Samples from two independent production runs were analyzed to confirm reproducibility. SEC revealed broad, shouldered peaks eluting between 10 and 13 mL for M2ex3-hTIP60 and at ∼11.8 mL for M2ex3-cf2TIP60-Q (**Fig. 6a**, column 1). DSC showed thermostability profiles largely consistent with their Arah2-displaying counterparts, with T_m1_ values of 59.7 and 46.0 °C, respectively, closely matching those of hTIP60 and cf2TIP60-Q displaying Ara h 2. Interestingly, M2ex3 exhibited a thermal unfolding transition similar to that of Ara h 2, with T_m2_ values of 74.8–75.5 °C (**Fig. 6a**, column 2; **Fig. S4c**). DLS analysis showed Dh values of 30.1-30.5 nm for both constructs (**Fig. 6a**, column 3), consistent with the structural models (**Fig. 6a, left**). nsEM further confirmed the structural integrity of mAb148-purified M2ex3-hTIP60 and M2ex3-cf2TIP60-Q, revealing well-formed nanoparticles. Consistent with the SEC profile, nsEM analysis revealed a mixture of smaller and normal-sized M2ex3-hTIP60 particles (**Fig. 6a**, column 1; **Fig. S4d**), a pattern not readily observed in allergen-displaying SApNPs.

**Fig. 6.**
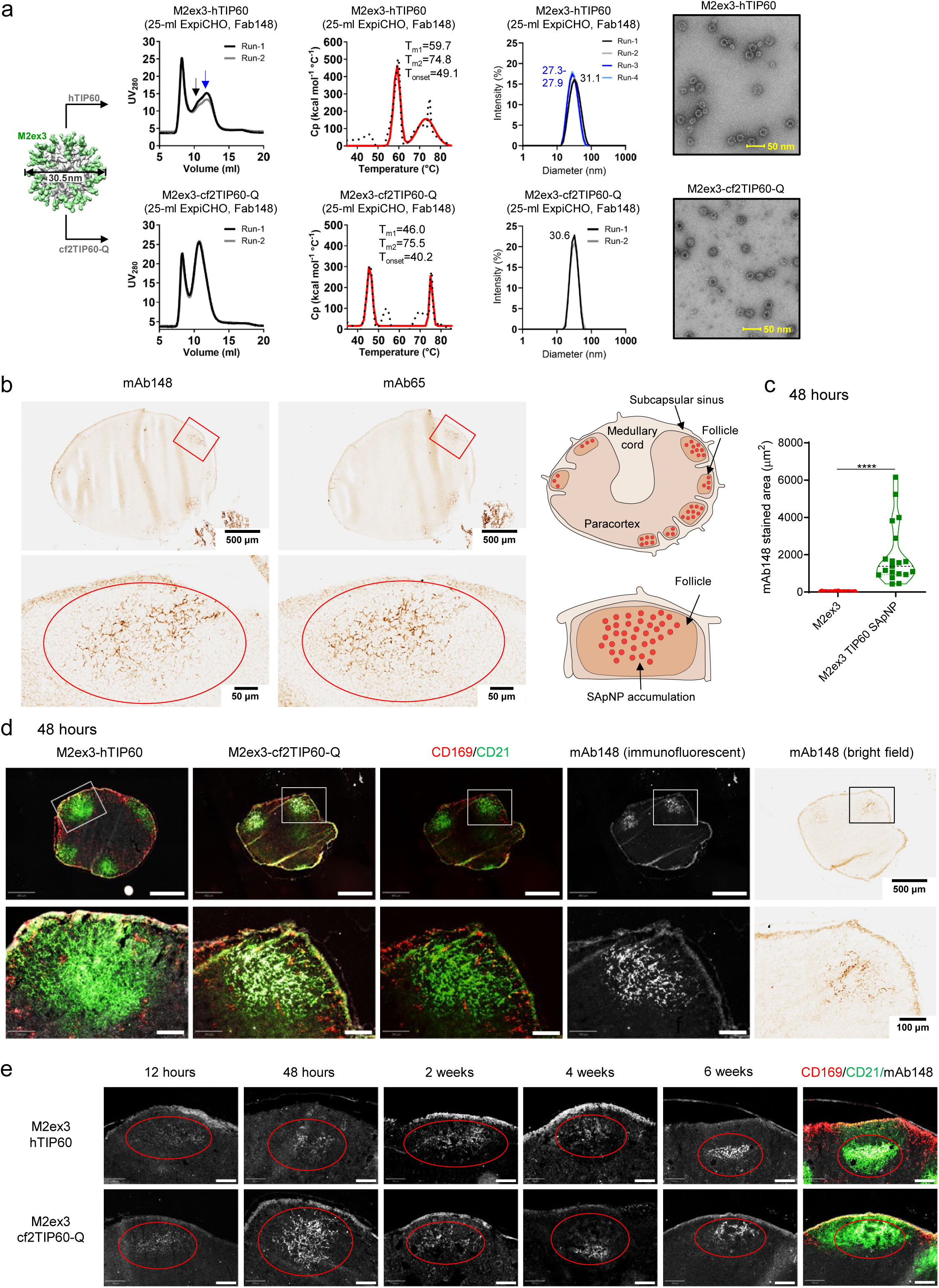
Distribution, trafficking, and retention of TIP60 SApNPs in lymph nodes. (**a**) characterization of M2ex3-hTIP60 and M2ex3-cf2TIP60-Q. (**b**) Distribution of M2ex3-displaying TIP60 constructs in a lymph node at 48 h after a single-dose injection (10 μg/injection, 40 μg/mouse, left; n = 3-5 mice/group). Each mouse received 80 μl of antigen/CpG mixture containing 40 μg SApNP immunogen and 40 μl CpG adjuvant. Sentinel lymph node sections were stained with mAb148 and mAb65. A schematic illustrating M2ex3–TIP60 SApNP accumulation in lymph node follicles is shown at right. (**c**) Quantification of M2ex3 immunogen accumulation in lymph node follicles at 48 h post–injection. (**d**) Association of M2ex3–displaying TIP60 constructs with FDC networks at 48 h. Left panels correspond to M2ex3–hTIP60 and M2ex3–cf2TIP60–Q, respectively. Both TIP60 SApNPs colocalized with FDC networks. Immunofluorescent images are pseudo–colored (CD21⁺, green; CD169⁺, red; mAb148, white). Scale bars: 500 μm (whole lymph node) and 100 μm (enlarged follicle). (**e**) Trafficking and retention of TIP60 SApNPs in lymph node follicles from 12 h to 6 weeks after a single–dose injection. Scale bar: 100 μm. Data in panel (c) are shown as mean ± SD. Statistical analysis was performed using two–tailed unpaired t-tests with Welch’s correction. ****p < 0.0001.

Having completed the in vitro characterization, we next studied the lymph node transport and distribution of the two M2ex3-displaying TIP60 SApNPs. Mice were immunized by intradermal injection into all four footpads (10 μg per footpad), and sentinel brachial and popliteal lymph nodes were collected from both sides of the body 48 h later. Immunostaining of lymph node sections was performed using mAb148 (*93*) and mAb65 (*94*) to detect M2ex3 displayed on TIP60 SApNPs. Images obtained with both antibodies showed similar localization of TIP60 SApNPs in the centers of lymph node follicles (**Fig. 6b**, images and schematics), consistent with distribution patterns observed in our previous vaccine studies (*71, 74–77*). Because mAb148 provided a better signal-to-noise ratio (*76*), it was selected for subsequent analyses of follicular accumulation, cellular interactions, and retention. Compared with the soluble M2ex3 trimer (*76*), quantification of mAb148-stained follicular areas showed that the M2ex3-cf2TIP60-Q group exhibited ∼155-fold greater follicular accumulation at 48 h (**Fig. 6c**), highlighting the markedly enhanced follicular delivery and retention conferred by the TIP60 SApNP platform.

We then examined the interactions between M2ex3-displaying TIP60 SApNPs and cellular populations within lymph node follicles. Our previous studies established that FDC networks are the primary follicular structures responsible for retaining large antigen-presenting nanoparticles, including SARS-CoV-2 spike, HIV-1 Env, SUDV GPΔmuc, and influenza M2ex3 and hemagglutinin SApNPs (*71, 74–77*). To determine whether M2ex3-hTIP60 and M2ex3-cf2TIP60-Q follow the same pattern, we collected sentinel lymph nodes at 48 h and stained the sections with mAb148 (white) for M2ex3, anti-CD21 antibodies (green) for FDCs, and anti-CD169 antibodies (red) for subcapsular sinus macrophages (**Fig. 6d**). Immunofluorescence imaging showed clear colocalization of TIP60 SApNPs with FDC networks, supporting the notion that FDCs mediate follicular accumulation and retention of M2ex3-displaying TIP60 SApNPs.

Lastly, we assessed long-term retention of both TIP60 SApNP formulations in lymph node follicles following a single intradermal dose (4 footpads, 10 μg/footpad) (**Fig. 6e**). Histological images revealed that both M2ex3-hTIP60 and M2ex3-cf2TIP60-Q SApNPs were detected in follicles within 12 h and remained detectable throughout the 6-week period. These findings are consistent with our previous observations (*71, 74–77*) that small antigens (<15 nm) are rapidly cleared from follicles within 48 h, whereas large antigen-displaying SApNPs (30–100 nm) remain in follicles for up to 8 weeks, underscoring the advantage of TIP60 SApNPs in sustaining follicular antigen availability and promoting effective humoral immune responses during AIT.

### Interaction of allergen-displaying SApNPs with various immune cell groups in lymph nodes

FDC networks form a reservoir within lymph node follicles that captures and preserves native-like antigens, immune complexes, particulate immunogens, viruses, and bacteria on their surfaces and dendrites through complement receptor-mediated mechanisms (*95–98*). Our previous transmission electron microscopy (TEM) analyses revealed that FDC networks retain ovalbumin-conjugated gold nanoparticles (*99, 100*) and antigen-presenting SApNPs (*71, 74, 75, 77*) along their surfaces and dendrites. To assess how FDC networks interact with cf2TIP60 SApNPs, we visualized the interface between FDC dendrites and B cells after intradermal injection of CpG–adjuvanted cf2TIP60–Q–Arah2 or cf2TIP60–Q–Arah6–Arah2 SApNPs into the hind footpads (2 footpads, 50 μg/footpad). Fresh popliteal sentinel lymph nodes were isolated 12 and 48 h after injection and processed for TEM. At both time points, TEM images revealed the characteristic morphology of long FDC dendrites engaging B cells within follicles (**Figs. 7a-c**). Intact SApNPs (round granules, yellow arrows) were aligned along FDC dendrites at both time points (**Figs. 7a-c** and **Fig. S5a-b**). At 48 h, cf2TIP60 SApNPs were also observed within endolysosomes of B cells (**Fig. 7d** and **Fig. S5c**). These findings demonstrate that FDC networks collect and present allergen-displaying SApNPs, with their long dendrites maximizing allergen-B cell interactions.

**Fig. 7.**
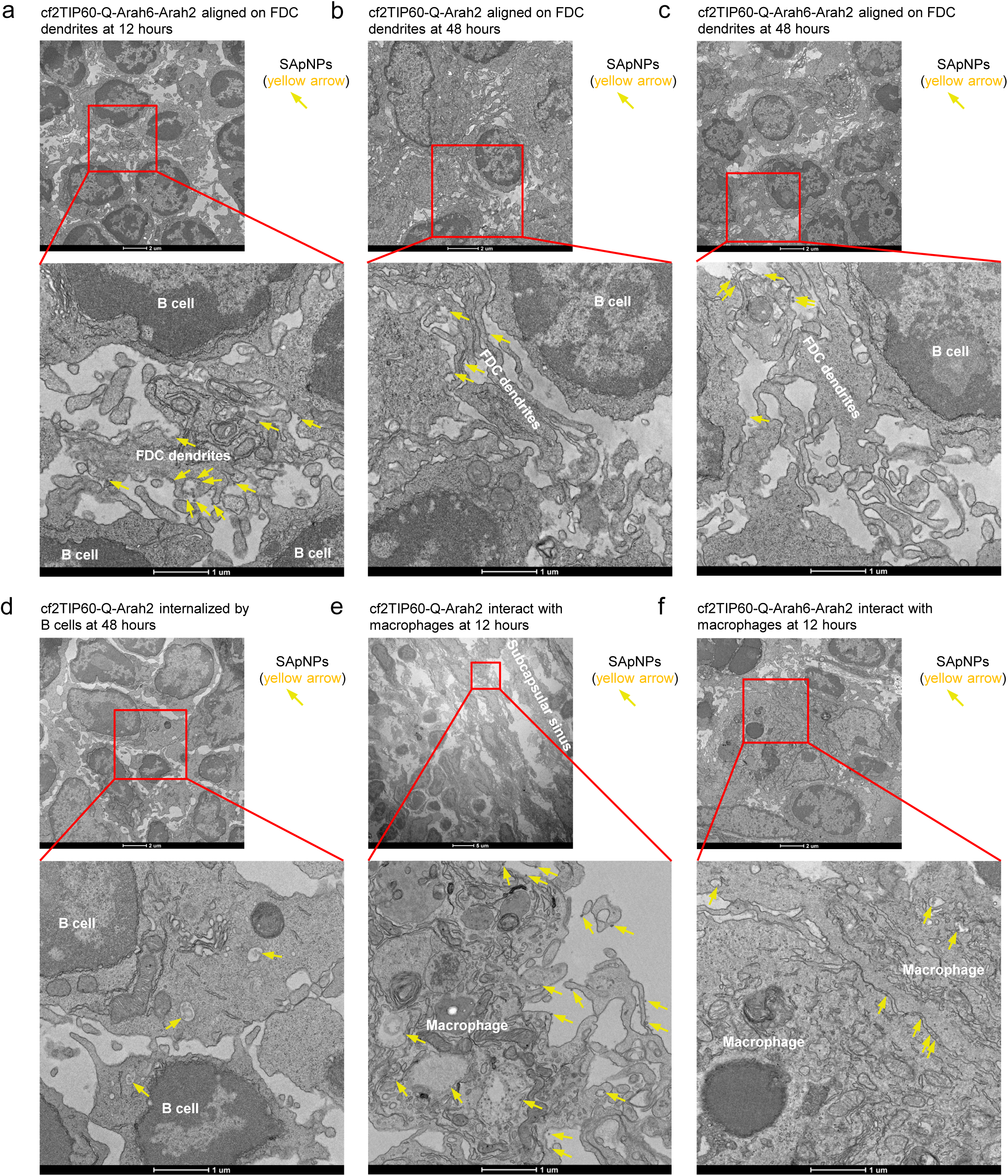
Interaction of allergen-displaying SApNPs with FDCs, B cells, and macrophages in lymph nodes. (**a**) TEM images showing cf2TIP60-Q-Arah6-Arah2 retained along FDC dendrites at 12 h after injection. TEM images of (**b**) cf2TIP60-Q-Arah2 and (**c**) cf2TIP60-Q-Arah6-Arah2 aligned along FDC dendrites at 48 h post-injection. Mice received 200 μl of antigen/CpG mixture containing 100 μg TIP60 SApNP and 100 μl CpG adjuvant, delivered into two hind footpads (50 μg/footpad). TIP60 SApNPs are indicated by yellow arrows. (**d**) TEM images showing cf2TIP60–Q–Arah2 (yellow arrows) within B–cell endolysosomes at 48 h. TEM images of (**e**) cf2TIP60–Q–Arah2 and (**f**) cf2TIP60–Q–Arah6–Arah2 (yellow arrows) located on the surface or within endolysosomes of macrophages at 12 h post–injection. TEM images were performed on two popliteal lymph nodes for each SApNP construct and time point.

Phagocytic cell populations, including macrophages and dendritic cells, not only internalize and process antigens to facilitate adaptive immunity but also transport intact antigens to migrating B cells and deposit them onto FDC networks in their native conformation (*96, 97, 101–104*). Here, we assessed interactions between phagocytic cells and cf2TIP60 SApNPs. At 12 h, both cf2TIP60-Q-Arah2 and cf2TIP60-Q-Arah6-Arah2 SApNPs were found on the surface and within endolysosomes of phagocytic cells (**Fig. 7e, f** and **Fig. S5d**). Consistent with our previous findings for SApNPs presenting diverse viral antigens (*71, 74, 75, 77*), these results indicate that cf2TIP60 SApNPs distribute across both intercellular and intracellular compartments within lymph nodes. The two cf2TIP60-Q SApNPs, despite displaying different allergens, exhibited similar associations with FDC networks, B cells, and macrophages, reflecting a shared trafficking and processing pathway. FDC networks retained SApNPs along their dendrites and presented native allergens to B cells within follicles, while a substantial fraction of SApNPs was internalized within endolysosomes of phagocytic cells. Together, these observations highlight the coordination among phagocytic cells, B cells, and FDC networks in distributing, retaining, and presenting SApNP-displayed antigens to support effective humoral immune activation.

### Characterization of germinal center responses induced by allergen-displaying SApNPs

In lymph node follicles, GCs are the sites where B cell somatic hypermutation, selection, affinity maturation, and class switch recombination occur, ultimately generating immune memory and high–affinity antibody responses following vaccination (*97, 105–107*). FDC networks serve as critical stromal components supporting GC formation and maintenance by retaining and presenting antigens within follicles (*95, 108*). Based on the prolonged follicular retention of cf2TIP60 SApNPs, we hypothesized that these SApNPs would elicit more robust and long-lived GC reactions than soluble allergens. To test this, we first characterized GC responses induced by CpG–adjuvanted cf2TIP60–Q–Arah2 following a single–dose injection (4 footpads, 10 μg/footpad). We used immunohistological staining to identify GC B cells (GL7⁺) in lymph node sections (*71, 74–77*). At 2 weeks post–injection, cf2TIP60–Q–Arah2 induced well–organized GCs that were closely associated with CD21⁺ FDC networks (green) and exhibited distinct dark zone (DZ) and light zone (LZ) compartments within B220⁺ follicles (blue) (**Fig. 8a**). We next compared GC responses induced by soluble monomers and SApNPs 6 weeks after a single–dose injection (**Figs. 8b, c**). GC size, defined as the follicular area occupied by GL7⁺ cells, was used as an image–based metric of GC magnitude (*71, 74–77*). CpG–adjuvanted allergen monomers and SApNPs generated measurable GC reactions, with cf2TIP60–Q–Arah6–Arah2 producing the largest GCs at week 6. Relative to soluble monomers, cf2TIP60 SApNPs induced 1.8–2.2-fold larger GC responses, demonstrating the enhanced capacity of the TIP60 SApNP platform to sustain GC responses.

**Fig. 8.**
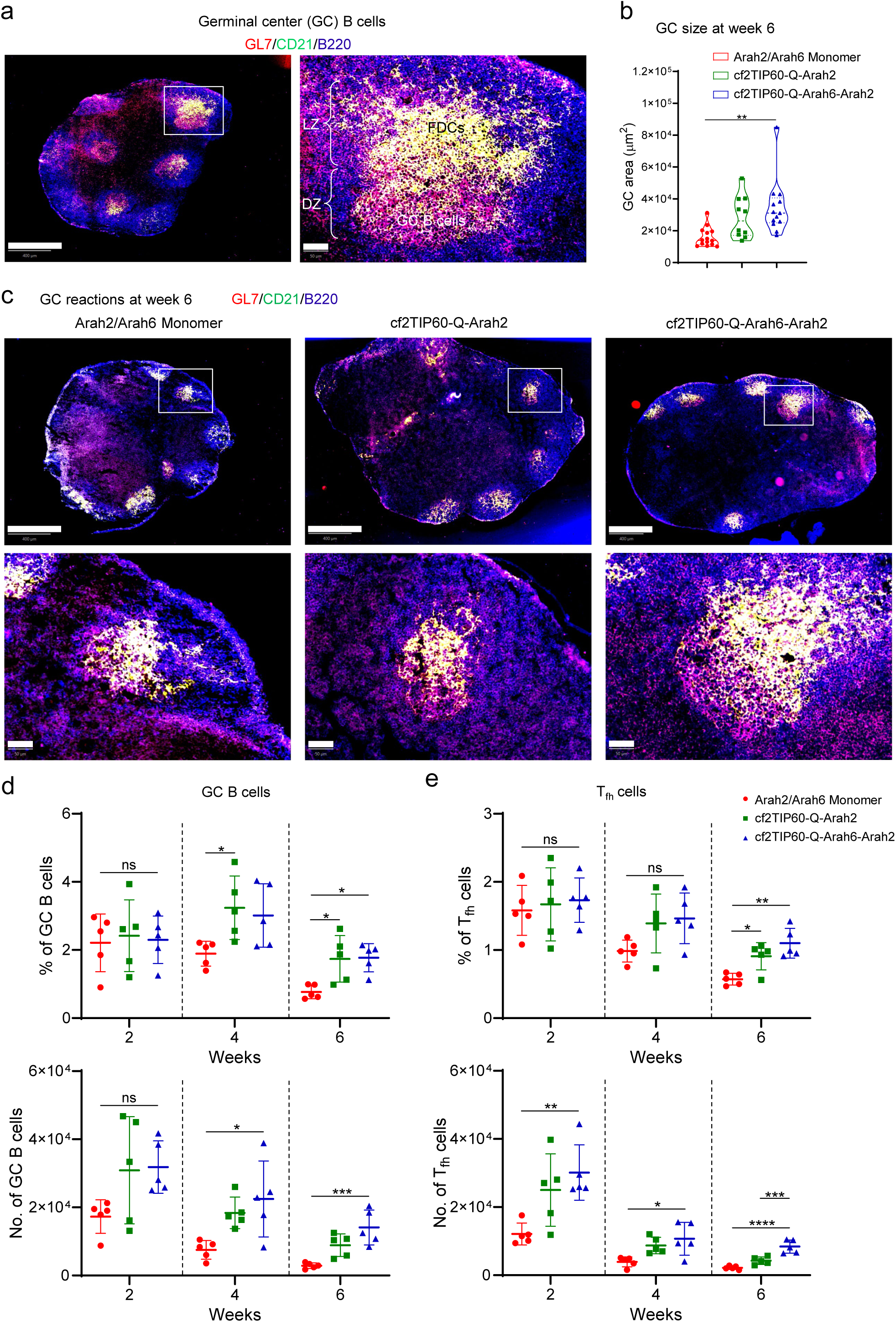
Induction of robust and long-lived germinal centers by allergen-presenting SApNPs in lymph nodes. (**a**) Left: Representative immunofluorescent images showing germinal centers (GCs) induced by CpG–adjuvanted cf2TIP60–Q–Arah2 at 2 weeks after a single–dose injection (10 μg/injection; 40 μg/mouse). Each mouse received 80 μl of antigen/CpG mixture containing 40 μg SApNP and 40 μl CpG adjuvant. Right: FDC networks (CD21⁺, green) associated with well–organized light zone (LZ) and dark zone (DZ) compartments within B–cell follicles (B220⁺, blue). GC B cells (GL7⁺, red) are primarily localized in the DZ. Scale bars: 500 μm (whole lymph node) and 50 μm (enlarged follicle). (**b**) Quantification of GC size from immunofluorescent images at 6 weeks after a single–dose injection (n = 5 mice/group). (**c**) Representative GC immunofluorescent images from allergen monomer and allergen–displaying SApNPs at 6 weeks post–immunization. Top: Scale bars = 500 μm (whole lymph node). Bottom: Scale bars = 50 μm (enlarged follicle). Quantification of (**d**) GC B cells and (**e**) T_fh_ cells by flow cytometry following a single-dose injection (n = 5 mice/group). Data are presented as mean ± SD. Statistical analysis was performed using one-way ANOVA followed by Tukey’s multiple comparison post hoc test for each time point. Significance: ns (not significant), *p < 0.05, **p < 0.01, ***p < 0.001, ****p < 0.0001.

To complement our semi–quantitative assessment of GC area by immunohistology, we further evaluated GC responses by flow cytometry using previously established protocols (*71, 74–77*). Mice were intradermally injected with the three allergen formulations, and sentinel lymph nodes were collected at 2, 4, and 6 weeks after a single–dose injection (four footpads, 10 μg per footpad). Fresh lymph node tissues were disaggregated into single–cell suspensions and stained with a defined antibody cocktail to quantify GC reactions based on the frequencies and absolute numbers of GC B cells (B220⁺GL7⁺CD95⁺) and T_fh_ cells (CD3⁺CD4⁺CXCR5⁺PD–1⁺) (**Fig. S6**). Flow cytometry revealed that all groups generated comparable frequencies of GC B cells and T_fh_ cells; however, cf2TIP60-Q-Arah6-Arah2 SApNPs elicited the highest absolute numbers of both cell populations at week 2 (**Figs. 8d, e**). Although GC B and T_fh_ cell populations declined over time, both cf2TIP60 SApNPs consistently outperformed soluble monomers at every time point. At week 6, cf2TIP60 SApNPs induced 3.1–4.9-fold more GC B cells and 2.0–3.8-fold more T_fh_ cells than monomers, mirroring the trends observed by immunohistology. Together, these results demonstrate that cf2TIP60 SApNPs elicit more robust and persistent GC responses than soluble monomers, supporting their potential to drive durable humoral immunity.

### Functional T cell responses induced by allergen-displaying SApNPs

T cells in splenic tissue comprise multiple functional subsets, including CD4⁺ and CD8⁺ T cells, which play central roles in regulating allergic inflammation. AIT reshapes these responses by modulating Th1/Th2/Th17 balance and CD4⁺/CD8⁺ cytokine pathways (*7, 27, 109*). Here, we characterized functional T–cell responses induced by soluble allergens and SApNPs using our established activation–induced marker (AIM) and intracellular cytokine staining (ICS) protocols (*76*). We implemented a 13–color spectral flow cytometry panel to quantify AIM expression and intracellular cytokine production (**Fig. S7**), enabling detailed analysis of antigen–specific T cell activation. Mice were intradermally immunized with CpG–adjuvanted allergen formulations, and spleens were collected 2 weeks after a single–dose injection (four footpads, 10 μg per footpad). Spleens were mechanically disaggregated, red blood cells removed, and splenocytes prepared for downstream stimulation. To evaluate Arah2–specific responses, splenocytes from mice immunized with mixed Arah2-His_6_/Arah6-His_6_ monomers, cf2TIP60–Q–Arah2, or cf2TIP60–Q–Arah6–Arah2 SApNPs were stimulated with Arah2–His_6_ prior to analysis. All three formulations elicited Th1–skewed responses, characterized by robust IFN–γ production, modest Th17 activity, and relatively low Th2 cytokines (**Fig. 9a**), consistent with effective allergen–specific T–cell activation.

**Fig. 9.**
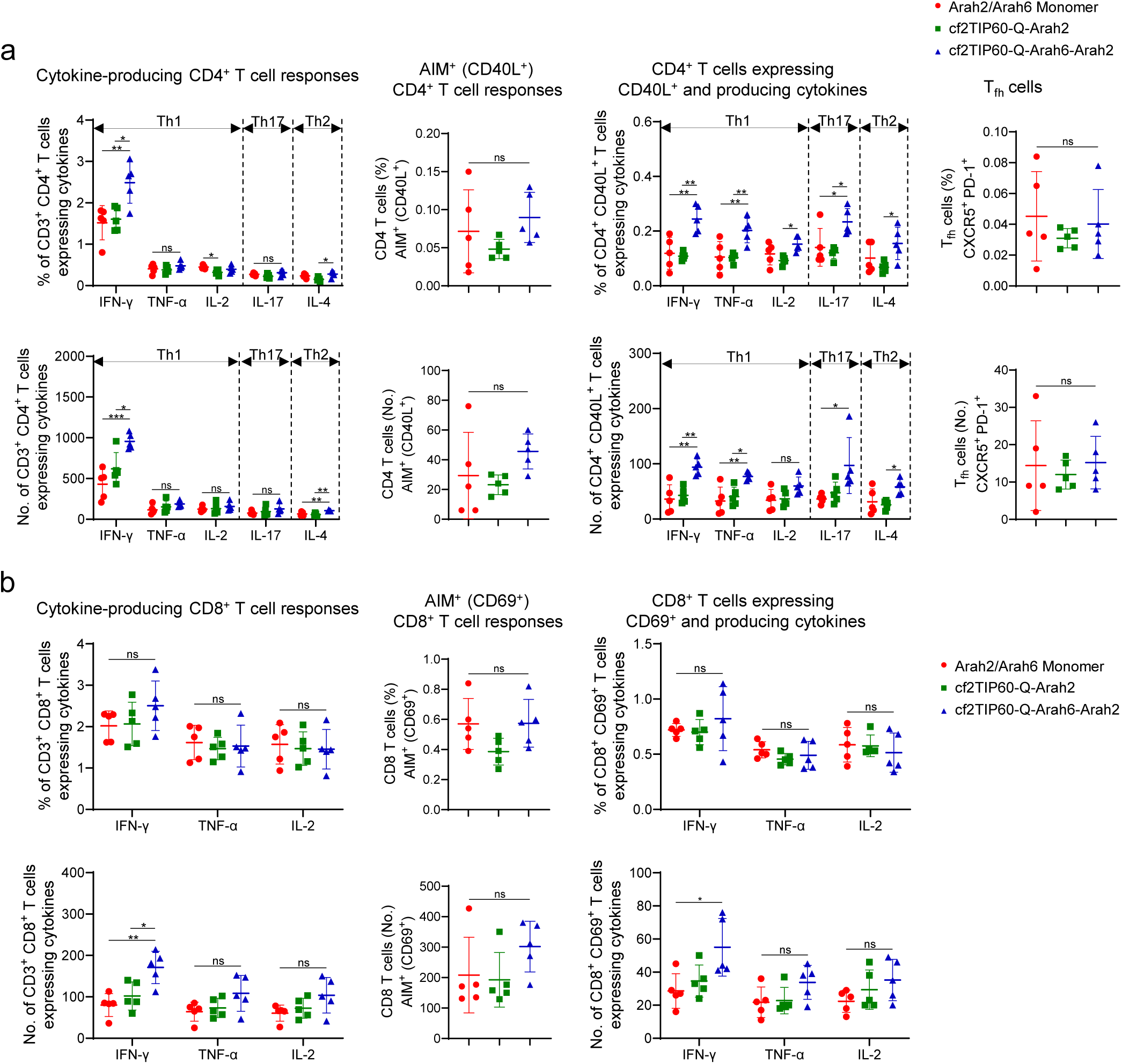
T cell responses induced by allergen and allergen-presenting SApNPs in the spleen. Quantification of the percentage and number of allergen–induced functional (a) CD4⁺ and (b) CD8⁺ T cell responses assessed by flow cytometry (n = 5 mice/group). Each mouse received 80 μl of an antigen–adjuvant mixture containing 40 μg allergen (10 μg per footpad) formulated with CpG adjuvant, delivered intradermally into four footpads. Spleens were isolated 2 weeks after a single–dose injection. Splenocytes were stimulated with Arah2–His_6_, and functional Arah2–specific T–cell responses were evaluated using activation–induced marker (AIM) and intracellular cytokine staining (ICS) assays. Data are presented as mean ± SD. Statistical analysis was performed using one–way ANOVA followed by Tukey’s multiple comparison post hoc test for each cytokine. Significance: ns (not significant), *p < 0.05, **p < 0.01, ***p < 0.001.

Among the tested constructs, cf2TIP60–Q–Arah6–Arah2 SApNP induced the strongest CD4⁺ T cell responses, generating the highest frequencies and absolute numbers of IFN–γ⁺, TNF–α⁺, IL–2⁺, IL–17A⁺, and IL–4⁺ CD4⁺ T cells, as well as CD40L⁺ CD4⁺ T cells and CD40L⁺ cytokine–producing CD4⁺ T cells. T_fh_ cell frequencies did not differ significantly across groups. Importantly, cf2TIP60–Q–Arah6–Arah2 SApNP induced 2.2–fold more IFN–γ⁺ CD4⁺ T cells and 2.6–fold more IFN–γ⁺CD40L⁺ CD4⁺ T cells than the Arah2-His_6_/Arah6-His_6_ monomer group, demonstrating a markedly enhanced Th1–biased effector profile. Similarly, CD8⁺ T cell responses were strongest in the cf2TIP60–Q–Arah6–Arah2 SApNP group, which elicited higher frequencies of IFN–γ⁺, TNF–α⁺, and IL–2⁺ CD8⁺ T cells, as well as CD69⁺ CD8⁺ T cells and IFN–γ⁺CD69⁺ CD8⁺ T cells, than both the monomer and cf2TIP60–Q–Arah2 groups (**Fig. 9b**). These findings highlight the ability of multivalent SApNPs to enhance cytokine–producing effector T cell subsets. Overall, cf2TIP60–Q–Arah6–Arah2 SApNP induced a strong Th1–skewed response and greater functional CD4⁺ and CD8⁺ T cell activity than allergen monomers, consistent with the superior prophylactic and therapeutic protection observed in peanut–induced anaphylaxis models. These results support cf2TIP60–Q as a promising platform for peanut AIT.

### Evolution-inspired optimization and evaluation of more thermostable TIP60 SApNPs

Having characterized the in vitro and in vivo profiles of hTIP60 and cf2TIP60 SApNPs, including their trafficking, lymphatic retention, and immune activation when displaying Arah2 and Arah6-Arah2 allergens, we established the TIP60 platform as a robust delivery modality for peanut AIT. However, the ∼14 °C lower T_m_ of cf2TIP60 relative to its human counterpart hTIP60 remained a key limitation, motivating an evolution-inspired effort to recover thermostability while preserving the favorable properties of the catfish-derived scaffold.

Our engineering of the TIP60-based peanut AIT platform followed a stepwise strategy (**Fig. 10a**): we first replaced the human myosin-X segment with a catfish homolog to reduce human sequence content, thereby generating cf2TIP60, and subsequently improved its stability through structure-guided design to yield cf2TIP60-Q. We revisited the evolutionary divergence between human and catfish myosin-X to identify sequence features underlying the superior thermostability of hTIP60 (**Fig. 10a**, left top). Sequence comparison revealed two glycine residues unique to the catfish homolog (**Fig. 10a**, left bottom). Structurally, these two glycines are located at the turn of the myosin-X “hook” (**Fig. 10a**, right), which is essential for myosin movement (*110*). Because glycine imparts high backbone flexibility, we hypothesized that these residues contribute to the reduced thermostability of cf2TIP60 relative to hTIP60, potentially reflecting adaptation of catfish myosin-X to the physiological temperatures of ectothermic fish. To rigidify the catfish myosin-X segment within cf2TIP60, we introduced G114N and E120P/G121E substitutions, generating the cf2TIP60-Q-NPE variant (**Fig. 10a**, left bottom and right).

**Fig. 10.**
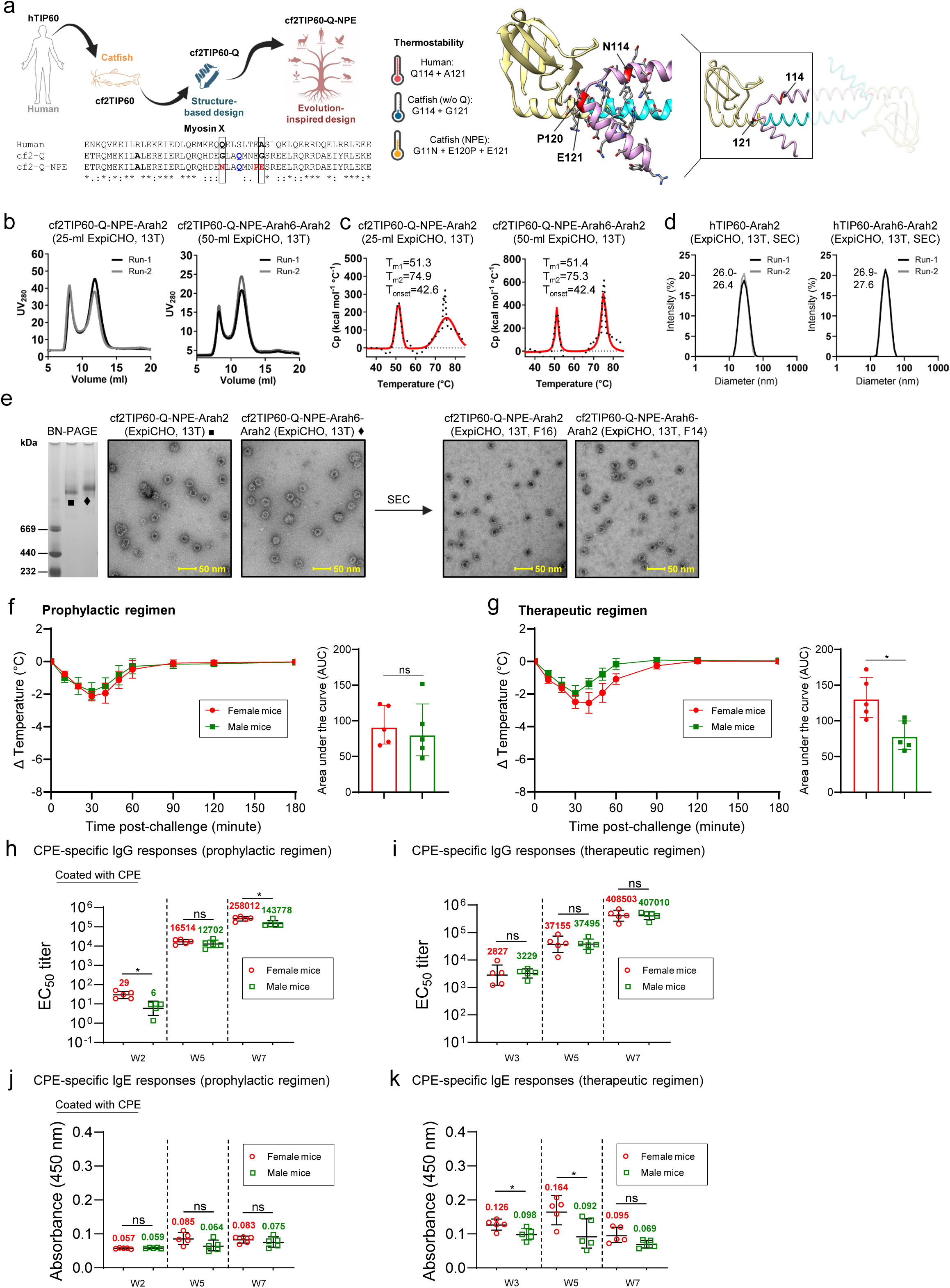
Evolution-inspired optimization and evaluation of thermostable TIP60 SApNPs against peanut-induced anaphylaxis in prophylactic and therapeutic peanut allergy models. (**a**) Overall strategy for rational design of thermostable TIP60 SApNPs. Left top: design progression from human-derived hTIP60 to catfish-derived cf2TIP60, structurally optimized cf2TIP60-Q, and evolution-inspired cf2TIP60-Q-NPE. Left bottom: alignment of the human myosin X sequence with cf2-Q and cf2-Q-NPE sequences. Right: glycine residues in cf2 myosin X are shown in red in a tube representation of the cf2TIP60 backbone, with an atomic view of the NPE mutations shown on a ribbon model. (**b**) Size-exclusion chromatography (SEC) profiles of ExpiCHO-expressed, 13T-purified cf2TIP60-Q-NPE-Arah2 and cf2TIP60-Q-NPE-Arah6-Arah2 from two independent production runs. (**c**) Differential scanning calorimetry (DSC) thermograms of 13T-purified samples from one production run. Experimental data (black points) and Gaussian fits (red lines) are shown, with melting temperatures (Tm1/Tm2) and onset temperature (Tonset) indicated. (**d**) Dynamic light scattering (DLS) profiles of ExpiCHO-expressed, 13T/SEC-purified cf2TIP60-Q-NPE-Arah2 and cf2TIP60-Q-NPE-Arah6-Arah2 from two independent production runs. Hydrodynamic diameters (Dh) are indicated. (e) BN-PAGE gels and negative-stain EM micrographs of cf2TIP60-Q-NPE-Arah2 and cf2TIP60-Q-NPE-Arah6-Arah2 (left panels), along with negative-stain EM micrographs after SEC purification (right panels). Scale bar, 50 nm. (**f**) In a prophylactic peanut allergy model, female and male mice were treated intradermally with cf2TIP60–Q–NPE SApNPs (10 µg total) formulated with CpG ODN 1826 (50 µg) (n = 5 per group). Two weeks later, mice were sensitized and challenged with CPE (200 µg/2000 µg), and rectal temperature was monitored for 180 min. Shown are time–dependent temperature curves and AUC values for temperature loss over 180 min. (**g**) In a therapeutic peanut allergy model, female and male mice were sensitized with CPE (200 µg, i.p.) at weeks 0 and 1 and treated intradermally two weeks later with adjuvanted cf2TIP60–Q–NPE SApNPs (10 µg total) (n = 5 per group). Two weeks after treatment, mice were challenged with CPE (2000 µg, i.p.), and rectal temperature was monitored for 180 min. Time–dependent temperature curves and AUC values for temperature loss over 180 min are shown. (**h-k**) Antibody responses: CPE–specific IgG titers in the (**h**) prophylactic and (**i**) therapeutic settings, and CPE–specific IgE responses in the (**j**) prophylactic and (**k**) therapeutic settings. EC_50_ titers were derived from serum ELISA against CPE. A_450_ values are reported for IgE responses. Geometric means are indicated on the plots, and data are presented as geometric mean ± geometric SD. Statistical analysis was performed using two–tailed unpaired t–tests with Welch’s correction for comparisons between two independent groups. Statistical significance is indicated as: ns (not significant) and *p < 0.05.

Following the established experimental workflow, we characterized the in vitro properties of cf2TIP60-Q-NPE SApNPs displaying Arah2 and tandem Arah6-Arah2 (**Fig. S8a**). SEC profiles demonstrated that the yield and composition of the NPE variants were highly similar to those of the parental cf2TIP60-Q constructs (**Fig. 10b**). Importantly, DSC analysis revealed a ∼5 °C increase in T_m_, with thermal transitions at 51.3 °C and 51.4 °C for the Arah2- and Arah6-Arah2-displaying SApNPs, respectively (**Fig. 10c** and **Fig. S8b**). These data support the hypothesis that residues within this hook region contribute substantially to the reduced thermostability of catfish myosin-X. DLS indicated that the Dh of the NPE variants remained comparable to that of the parental SApNPs (**Fig. 10d**). BN-PAGE analysis yielded well-defined bands for both cf2TIP60-Q-NPE constructs, with slower migration observed for the larger Arah6-Arah2-displaying SApNP (**Fig. 10e**, left). Finally, structural validation by nsEM confirmed homogeneous nanoparticle assembly; samples purified by 13T/SEC exhibited higher purity than those purified by 13T IAC alone and revealed a distinct electron-dense core (**Fig. 10e**, right; **Fig. S8c**).

Using the established prophylactic and therapeutic peanut allergy models (**Figs. 2, 3, 5**), we evaluated the more thermostable cf2TIP60–Q–NPE SApNPs for their potential in peanut AIT and assessed whether sex influences AIT efficacy (**Fig. 10f, g**). In the prophylactic setting, both female and male mice received a single intradermal dose of CpG–adjuvanted cf2TIP60–Q–NPE SApNPs (2.5 µg per footpad; 10 µg total). Two weeks later, mice were sensitized and challenged using the established CPE protocol (200 µg sensitization; 2000 µg challenge), and body temperature was monitored for 180 min (**Fig. 10f**). In both sexes, cf2TIP60–Q–NPE SApNPs conferred strong prophylactic protection, with temperature drops of 1.8–2.2 °C at 30 min and full recovery within 90 min, accompanied by low AUC values for temperature loss. These responses closely matched the protective profile of hTIP60–Arah6–Arah2 SApNPs, which produced a 2.1 °C temperature drop at 30 min and full recovery within 90 min (**Fig. 2e**), underscoring the robust prophylactic efficacy of the cf2TIP60–Q–NPE platform. In the therapeutic setting, sensitized female and male mice received a single intradermal dose of CpG-adjuvanted cf2TIP60–Q–NPE SApNPs (2.5 µg per footpad; 10 µg total). Two weeks later, mice were challenged with 2000 µg CPE, and body temperature was monitored for 180 min (**Fig. 10g**). In female mice, cf2TIP60–Q–NPE SApNPs produced therapeutic responses similar to those observed for earlier cf2TIP60 constructs, with temperature drops of 2.6–2.7 °C at 40 min and recovery to baseline within 120 min. In contrast, male mice exhibited greater therapeutic protection, with a smaller temperature drop of ∼2.0 °C at 30 min and full recovery within 90 min. These results are consistent with previous reports that female mice tend to develop more severe peanut–induced anaphylaxis than males (*111, 112*), highlighting sex–dependent differences in AIT efficacy. Overall, cf2TIP60–Q–NPE demonstrated strong prophylactic efficacy in both sexes and greater therapeutic efficacy in male mice, supporting its potential as a thermostable platform for peanut AIT.

We next assessed allergen–specific and sex–dependent antibody responses during AIT with cf2TIP60–Q–NPE SApNPs in both peanut allergy models. Sera collected at weeks 2/3, 5, and 7 from female and male mice were analyzed by CPE–coated ELISA (**Figs. 10h-k** and **Fig. S9**). In the prophylactic setting, cf2TIP60–Q–NPE SApNPs induced higher IgG titers in female mice at week 2 following a single–dose injection, with an EC_50_ value of 29, 5–fold higher than in males and reaching statistical significance (**Fig. 10h** and **Figs. S9a,b**). Notably, cf2TIP60–Q–NPE elicited higher week–2 IgG titers than either cf2TIP60–Q construct (**Fig. 5c**), supporting the optimized scaffold design. At weeks 5 and 7, after CPE sensitization and challenge, female mice continued to generate higher EC_50_ titers than males. In the therapeutic setting, female and male mice showed similar IgG responses across all three time points (**Fig. 10i** and **Figs. S9c,d**). These responses were comparable to those observed in female mice treated with hTIP60 and cf2TIP60 SApNPs (**Fig. 5d**), although the cf2TIP60-Q-NPE SApNP was associated with higher week-5 EC_50_ titers in both sexes. Together, these data demonstrate robust IgG immunogenicity and reveal sex-dependent differences in the prophylactic setting. IgE responses during peanut AIT were also evaluated. Overall, female mice showed IgE profiles similar to those observed in hTIP60 and cf2TIP60-Q SApNP treatment groups (**Figs. 5e, f**). In the prophylactic model, female mice generated slightly higher IgE levels than males at weeks 5 and 7, although these differences were not statistically significant (**Fig. 10j** and **Fig. S9e**). In the therapeutic model, sex-dependent differences were more pronounced. Female mice showed higher IgE responses after two doses of CPE sensitization at week 3 and again after cf2TIP60-Q-NPE treatment at week 5, reaching an A_450_ value of 0.164, which was 1.8–fold higher than that in males (**Fig. 10k** and **Fig. S9f**). These findings are consistent with previous reports that female mice tend to develop more severe peanut–induced anaphylaxis than males (*111–113*), highlighting the influence of sex on humoral responses and therapeutic outcomes in peanut AIT, **r**ather than an effect specific to the SApNP backbone.

## DISCUSSION

Most allergic diseases result from misdirected humoral immune responses to otherwise harmless environmental or food antigens. Peanut allergy is particularly challenging because it begins early in life, often persists into adulthood, and can cause severe anaphylaxis after minimal exposure (*1–9*). Although OIT, EPIT, SLIT, and adjunctive biologics have advanced peanut allergy treatment, their broader use remains constrained by treatment burden, adverse reactions, variable efficacy, and incomplete durability (*26–44*). Recently, CuMVtt-based VLPs displaying peanut allergens have shown promising prophylactic and therapeutic efficacy in mouse models, highlighting the potential of the VLP approach for peanut AIT (*51, 57, 59, 64*). An important goal for next-generation AIT is to redirect allergen-specific immunity more efficiently toward a protective state.

Here, we investigated a rationally engineerable nanoparticle platform for peanut AIT. By displaying Arah2 and tandem Arah6–Arah2 on human-derived 60-mer TIP60 (*79–81*) and catfish-derived variants, we established a modular SApNP platform that combines multivalent allergen presentation with tunable structural and biophysical properties. Across prophylactic, therapeutic, and severe peanut allergy models, TIP60 SApNPs consistently provided greater protection than soluble allergen monomers. Notably, this protection was achieved after only one or two SApNP administrations. A single prophylactic dose markedly reduced the temperature drop following peanut challenge, while a single treatment after sensitization provided substantial therapeutic protection. Efficacy was maintained in an even more stringent model in which mice underwent sensitization and anaphylactic challenge before treatment. Under these severe disease conditions, one SApNP treatment reduced responses to repeated challenges, whereas two treatments provided stronger and more durable protection. Although mouse models cannot be directly compared with clinical AIT, these results suggest the potential to reduce the treatment burden associated with conventional AIT. SApNP treatment also did not cause a detectable temperature drop in sensitized mice, supporting further evaluation of this approach for peanut AIT. The optimized cf2TIP60-Q-NPE scaffold retained strong prophylactic efficacy in both sexes, whereas therapeutic protection was greater in male mice, consistent with previous reports of greater peanut-induced anaphylaxis in female mice (*111–113*). Thus, sex may influence both disease severity and treatment outcome and should be considered in future preclinical studies.

Similar to VLPs (*57–61*), the type and balance of antibody responses induced during AIT may help explain SApNP-mediated protection. Soluble allergens are less effective at promoting sustained multivalent BCR engagement and, in a Th2-biased environment, may favor IgE-skewed responses. In contrast, SApNPs display allergens as repetitive arrays that promote BCR clustering (*114*), enhance lymph node retention, and sustain GC responses. Consistent with this model, both hTIP60- and cf2TIP60-based SApNPs induced substantially higher peanut-specific IgG responses than soluble monomers while limiting IgE escalation. This distinction was particularly evident in the therapeutic setting, where IgE levels increased markedly in the monomer and PBS groups after sensitization but remained substantially lower in SApNP-treated mice. Thus, particulate allergen presentation appears to redirect rather than simply suppress allergen-specific immunity, favoring a protective IgG/IgE balance. The resulting IgG antibodies may act through two complementary mechanisms (*115–117*). First, high-affinity IgG can sequester allergens and block IgE-accessible epitopes, thereby reducing cross-linking of FcεRI-bound IgE and effector-cell degranulation. Second, IgG–allergen immune complexes can engage FcγRIIb, suppressing mast cell and basophil activation through inhibitory co-signaling with FcεRI and potentially dampening allergen-specific B cell activation by co-engaging the BCR (*57–61*). These mechanisms are supported by previous AIT and VLP studies and may explain the protective antibody profile observed here, although IgG transfer or depletion studies remain important for direct validation. Notably, in line with their AIT outcomes, female mice also showed distinct IgG and IgE profiles, consistent with previous reports of sex-dependent differences in peanut allergy (*111–113*).

Our tissue and cellular analyses provide mechanistic insights into why particulate and soluble allergens produce different AIT outcomes. Overall, SApNP protection was associated with prolonged follicular retention, sustained GC responses, and enhanced functional T cell responses (*71, 74–77, 118*). After efficient trafficking to the lymph nodes, TIP60 SApNPs accumulated in follicles and remained detectable for 6 weeks after a single administration. Immunofluorescence demonstrated close association with FDC networks, while TEM directly visualized intact allergen-displaying particles along FDC dendrites near B cells and within B cell and phagocytic-cell endolysosomes. These findings suggest that SApNPs alter the geometry and duration of allergen exposure: repetitive allergen arrays enhance initial BCR engagement, whereas prolonged retention provides a persistent source of native-like antigen for repeated B cell encounters and GC selection (*52, 96, 97, 106, 118*). Consistent with this model, thermostable cf2TIP60-Q SApNPs generated larger and more persistent GCs and maintained greater numbers of GC B cells and T_fh_ cells than soluble monomers. The TIP60 SApNP platform also enhanced functional T cell responses during AIT. Although all CpG-adjuvanted formulations were predominantly Th1-skewed, the cf2TIP60-Q-Arah6-Arah2 SApNP induced the strongest CD4⁺ and CD8⁺ responses, including enhanced IFN-γ⁺ and cytokine-producing CD40L⁺ CD4⁺ T cell responses. Such Th1-biased activity may help counterbalance the Th2-dominated environment of peanut allergy, while CD40L-expressing CD4⁺ cells may further support GC and B cell responses (*119–121*). The CpG adjuvant undoubtedly contributed to this polarization, but the stronger responses to SApNPs than to monomers indicate an additional effect of particulate allergen presentation. Together, these findings suggest that SApNP efficacy arises from coordinated remodeling of both humoral and cellular immunity through sustained allergen presentation in a repetitive, multivalent format.

Another major finding of this work is that TIP60 can be systematically engineered rather than used as a fixed carrier. Scaffold compatibility was clearly antigen dependent: FR, E2p, and I3-01v9, despite their success with many viral antigens (*67–78*), failed to yield Ara h 2/6-displaying particles in the ExpiCHO system, whereas hTIP60 supported efficient expression and assembly. TIP60 geometry could also be altered by inserting a single α-helical heptad into the myosin-X coiled-coil. This increased particle size and produced strong prophylactic protection but reduced thermostability and structural homogeneity, showing a tradeoff between geometric tuning and biophysical robustness. Replacement of the human myosin-X segment with evolutionarily distant catfish homologs reduced human sequence content while preserving assembly, with cf2TIP60 emerging as the most favorable scaffold. Structure-guided introduction of L117Q provided modest stabilization in the cf2TIP60-Q construct, which was tested extensively in vivo, while subsequent evolution-inspired substitutions in the myosin-X hook generated cf2TIP60-Q-NPE and increased T_m_ by ∼5 °C without compromising particle size, assembly, yield, or protective efficacy. Notably, cf2TIP60-Q-NPE elicited stronger early IgG responses than the parental cf2TIP60-Q constructs, suggesting a link between improved scaffold stability and better immunological performance. Thus, TIP60 provides an engineerable platform in which scaffold sequence, particle dimensions, allergen composition, and stability can be systematically optimized for AIT.

Future studies may focus on several directions. First, further engineering of cf2TIP60-Q-NPE should aim to increase thermostability while preserving its favorable expression, assembly, and immunological properties. Although the ∼5 °C improvement is substantial, the catfish-derived scaffolds remain less thermostable than hTIP60. Additional optimization based on structural and evolutionary principles may further improve manufacturability and AIT performance. Second, the allergen repertoire displayed on TIP60 can be expanded. Ara h 2 and Ara h 6 are among the most clinically important peanut allergens (*15–19*), but patients vary in their sensitization to individual peanut allergens; incorporating additional allergens, particularly Ara h 1 and Ara h 3, may broaden coverage across diverse sensitization profiles (*122, 123*). Finally, the design principles established here should be applicable beyond peanut allergy. Major allergens from tree pollen, grass pollen, and house dust mite could be displayed similarly on optimized TIP60 scaffolds, extending the platform across environmental and food allergies (*124, 125*). Continued scaffold optimization and expanded allergen coverage may establish TIP60 SApNPs as a broadly applicable platform for next-generation AIT, redirecting allergic immunity toward durable protection.

## METHODS

### Structure-guided rational design of peanut allergen-displaying nanoparticles

Sequences of Ara h 2 (GenBank: QGA89290.1; amino acids A21–G156) and Ara h 6 (GenBank: AAD56337.1; amino acids A21–C145) were fused at the C-terminus to a His6 tag to generate monomeric allergens. Crystal structures of Ara h 2 (PDB: 8DB4 (*89*)) and Ara h 6 (PDB: 8SJA (*88*)) were used to build full-length atomic models using the homology modeling program NEST (*126*). Initial models of the hTIP60-Arah6 and hTIP60-Arah6-Arah2 subunits were generated by manually docking the allergen models onto a single hTIP60 subunit (PDB: 7EQ9 (*79*)) using UCSF Chimera (*127*). Flexible linkers (GGGGS) between the hTIP60 backbone and Ara h 6, and between Ara h 6 and Ara h 2, were built and optimized using the loop modeling program LOOPY (*128*). Full nanoparticle models were generated by applying symmetry parameters from the cryo-EM structure (PDB: 7EQ9 (*79*)). Modeling was performed iteratively to ensure that the linker length allowed both allergens to be displayed on the particle surface without steric clashes. To increase the display capacity, the hTIP60 backbone was expanded by inserting a 7-residue engineered heptad repeat (EAAKAAA) between the Sm-like domain and the hMyoX segment. A 9-residue α-helix (AEAAKAAAA), modeled from a transcription factor template (PDB: 6G6L, residues 953–961), was connected to the C-terminus of the Sm-like domain and the N-terminus of the hMyoX segment by superposing the backbone atoms of the flanking terminal residues. The expanded hTIP60-Ext backbone was compared to the original hTIP60 backbone to assess its structural suitability for assembly. To identify non-human hMyoX homologs, the 51-residue hMyoX segment was searched against the nonredundant protein database (nr) using Position-Specific Iterated BLAST (PSI-BLAST) (*92*) with the following parameters: 40–48% sequence identity, 100% query coverage, and 20,000 sequences. A total of 68 sequences meeting these criteria were identified, all from fish (NCBI taxid: 7898). After removing a low-quality sequence, the remaining 67 sequences were aligned using CLUSTALW (*129*) and used to construct a neighbor-joining phylogenetic tree (*130*). Seven sequences from diverse branches were selected to replace hMyoX in TIP60-Arah2 constructs for experimental screening. Two catfish-derived MyoX-like sequences, cf1 (GenBank: KAI5606636.1) and cf2 (GenBank: XP_026992614.2), were selected based on TIP60-Arah2 expression yield. The cf2TIP60 subunit was modeled with NEST (*126*) using hTIP60 as the template, and the L117Q mutation was introduced using SCAP (*131, 132*). Construct sequences are detailed in **Fig. S1a**, **Fig. S2a**, **Fig. S4a**, and **Fig. S8a**, and structural renderings of allergens/antigens and TIP60 nanoparticles in **Fig. 1**, **Fig. 4**, **Fig. 6**, and **Fig. 10** were generated using UCSF Chimera (*127*).

### Expression and purification of peanut allergen-displaying nanoparticles

Soluble allergens, Arah2-His_6_ and Arah6-His_6_, and allergen-displaying TIP60 nanoparticles were produced in ExpiCHO cells (Thermo Fisher). Briefly, ExpiCHO cells were thawed and incubated in ExpiCHO^TM^ Expression Medium (Thermo Fisher) in a shaker incubator at 37 °C, at 135 rpm, with 8% CO_2_. When the cells reached a density of 10 × 10^6^ ml^−1^, ExpiCHO^TM^ Expression Medium was added to adjust the cell density to 6 × 10^6^ ml^−1^ for transfection. ExpiFectamine^TM^ CHO-plasmid DNA complexes were prepared following the manufacturer’s instructions. For a 25-ml transfection volume, 25 μg of allergen plasmid and 80 μl of ExpiFectamine^TM^ CHO reagent were mixed in 2 ml of cold OptiPRO^TM^ medium (Thermo Fisher). After the first feed on day 1, ExpiCHO cells were cultured at 33 °C, at 115 rpm, with 8% CO_2_ according to the Max Titer protocol, with a second feed on day 5. Supernatants were harvested 13–14 days post-transfection, clarified by centrifugation at 3,724 × g for 25 min, and filtered through a 0.45 μm membrane (Thermo Fisher). Soluble allergens were purified using Ni Sepharose excel resin (Cytiva); bound proteins were eluted twice with 15 ml of 0.5 M imidazole and buffer-exchanged into Tris-buffered saline (TBS; pH 7.2). Two Ara h 2-specific antibodies, 13T and 15T (*89, 133*), were used to prepare affinity columns for IAC purification, as previously described for antigen-presenting nanoparticle vaccines (*67–78*). The 13T antibody column was used throughout this study to ensure consistency in IAC purification. TIP60-allergen proteins were eluted three times with 5 ml of 0.2 M glycine (pH 2.2), immediately neutralized with 0.5 ml of 2 M Tris base (pH 9.0), and buffer-exchanged into phosphate-buffered saline (PBS; pH 7.2). Nickel-purified soluble allergens and IAC-purified TIP60 nanoparticles were further polished by SEC using a Superdex 75 Increase 10/300 GL column and a Superose 6 Increase 10/300 GL column (Cytiva), respectively. Selected SEC fractions were flash-frozen in liquid nitrogen and stored at −80 °C until further use.

### Differential scanning calorimetry (DSC)

Thermal stability, characterized by melting temperature (T_m_) and onset of melting (T_onset_), was assessed for Ara h 2 (Arah2-His_6_) and TIP60-allergen nanoparticles when sufficient protein was available following affinity chromatography and, in some cases, SEC. DSC was performed on a MicroCal PEAQ-DSC instrument (Malvern). Purified samples in phosphate-buffered saline (PBS) were diluted to a final concentration of 0.1–5 μM. Thermal denaturation was monitored at a scan rate of 60 °C h⁻¹ over a temperature range of 20–100 °C. Data processing, including buffer background correction, concentration normalization, and baseline subtraction, was performed using MicroCal PEAQ-DSC software. Gaussian fitting was carried out using GraphPad Prism (version 10.3.1). DSC measurements were performed in duplicate for most hTIP60 and cf2TIP60 variants when expression yield permitted.

### Dynamic light scattering (DLS)

The hydrodynamic (Dh) size and polydispersity of allergen-displaying TIP60 nanoparticles were characterized using a Zetasizer Ultra instrument (Malvern). IAC/SEC-purified nanoparticle samples were diluted to approximately 0.2 mg ml⁻¹ in PBS, and 30 μl of each sample was loaded into a quartz batch cuvette (Malvern, cat. no. ZEN2112). Measurements were performed at 25 °C in backscatter mode (173°). Data acquisition and processing were performed using Zetasizer Ultra software to generate intensity-weighted particle size distributions. The resulting size distribution profiles were plotted using GraphPad Prism (version 10.3.1).

### Negative-stain electron microscopy (nsEM)

The nsEM analysis of allergen-displaying TIP60 nanoparticles, including both IAC-purified and IAC/SEC-purified variants, was performed by the Core Microscopy Facility at Scripps Research. Samples were diluted in PBS to a final concentration of 0.01 mg ml⁻¹. Carbon-coated copper grids (400 mesh) were glow-discharged for 60 s before 8 μl of each sample was adsorbed onto the grid surface for 2 min. Following adsorption, excess liquid was removed, and the grids were negatively stained with 2% (w/v) uranyl formate for 2 min. Excess stain was wicked away, and the grids were air-dried. Grids were imaged at an accelerating voltage of 120 kV using a Talos L120C transmission electron microscope (Thermo Fisher). Micrographs were acquired at a nominal magnification of 52,000× using a CETA 16M CMOS camera.

### Blue native polyacrylamide gel electrophoresis (BN-PAGE)

The allergen-displaying TIP60 nanoparticles were analyzed by BN-PAGE. Proteins were mixed with 4× native dye and loaded onto a 4–12% Bis-Tris NativePAGE^TM^ gel (Life Technologies). The gel was run for 2–2.5 h at 150 V with NativePAGE^TM^ running buffer (Life Technologies) according to the manufacturer’s instructions. Gels were stained with Coomassie Brilliant Blue R-250 (Bio-Rad) and destained using a solution of 6% ethanol and 3% glacial acetic acid. Gel images were acquired using a ChemiDoc^TM^ XRS+ imaging system and processed with Image Lab (version 6.1.0, build 7, Standard Edition; Bio-Rad).

### Enzyme-linked immunosorbent assay (ELISA)

Costar 96–well high–binding flat–bottom half–area plates (Corning) were coated with 50 µl PBS containing 10 µg CPE (Chondrex, Inc., cat. no. 3069) prepared in pH 8.0 buffer. Plates were incubated overnight at 4 °C, then washed five times with PBST (PBS + 0.05% Tween–20). Wells were blocked for 1 h at room temperature with 150 µl of 4% blotting–grade blocker (Bio–Rad) in PBS, followed by five PBST washes. Mouse sera were diluted 1:40 in blocking buffer and subjected to a 10–fold serial dilution. For each dilution, 50 µl was added to the wells and incubated for 1 h at room temperature, followed by five PBST washes. For secondary detection, plates received 50 µl of either HRP–conjugated goat anti–mouse IgG (1:3000; Jackson ImmunoResearch) or goat anti–mouse IgE (1:1000; Thermo Fisher) diluted in PBST. Plates were incubated for 1 h at room temperature and washed six times with PBST. Color development was performed using 50 µl TMB substrate (Life Sciences) for 3 min, and reactions were stopped with 50 µl of 2 N sulfuric acid. Absorbance was read immediately at 450 nm on a BioTek Synergy plate reader. EC_50_ values were calculated from full binding curves using GraphPad Prism. For samples with EC_50_ values below 0.1, values were set to 0.1 to maintain consistent plotting and comparison. Due to low IgE responses, OD_450_ values at 1:40 serum dilution are reported for IgE.

### Mouse immunization and peanut sensitization and challenge

All animal experiments were conducted in accordance with the guidelines of the Association for the Assessment and Accreditation of Laboratory Animal Care (AAALAC). All procedures were approved by the Institutional Animal Care and Use Committee (IACUC) at The Scripps Research Institute (TSRI). Six-to-eight-week-old female and male BALB/c mice were purchased from The Jackson Laboratory and housed in ventilated cages within environmentally controlled rooms in the Immunology Building at TSRI. To establish peanut-induced anaphylaxis, mice were sensitized with two intraperitoneal (i.p.) injections of CPE formulated 1:1 (v/v) with aluminum hydroxide (alum). Sensitization was performed at weeks 0 and 1 using either 50 µg or 200 µg CPE per dose. At week 3, mice were challenged i.p. with 200–2000 µg CPE. Rectal body temperature was measured using a digital thermometer every 10 min for the first 60 min and up to 180 min post-challenge. For prophylactic studies, mice received a single intradermal immunization at two footpads (2.5 µg per footpad; 10 µg total) with either Ara h 2/6 monomers or allergen-displaying SApNPs. All vaccine formulations contained CpG ODN 1826 (50 µg) as an adjuvant. PBS and CpG alone served as controls. Body temperature was monitored for 60 min after vaccination to assess acute responses. Two weeks later, mice were sensitized and challenged using the established 200 µg/2000 µg CPE protocol, and rectal temperature was recorded for 180 min post-challenge.

To evaluate therapeutic efficacy, mice were first sensitized with two i.p. injections of 200 µg CPE at weeks 0 and 1. Two weeks after sensitization, mice received a single intradermal injection with adjuvanted Ara h 2/6 monomers or allergen-displaying SApNPs (10 µg total). PBS served as the negative control. Body temperature was monitored for 60 min post-treatment. Two weeks later, mice were challenged with 2000 µg CPE, and rectal temperature was recorded for 180 min. To generate a heightened allergic state, mice were sensitized with two i.p. injections of 200 µg CPE formulated with alum at weeks 0 and 1, followed by an i.p. challenge with either 800 µg or 2000 µg CPE at week 3. Three weeks after this challenge, mice received a single intradermal injection of adjuvanted allergen-displaying SApNPs (10 µg total) or PBS as a control. Two weeks after treatment, mice underwent CPE challenge 1 (2000 µg), and rectal temperature was monitored for 180 min. Additional challenges (2 and 3) were performed subsequently, with temperature monitoring conducted identically. For enhanced therapeutic evaluation, mice with established severe allergy (generated as described above) received two intradermal doses of adjuvanted allergen-displaying SApNPs spaced two weeks apart. Two weeks after the second dose, mice underwent CPE challenge 1 (2000 µg), followed by challenge 2 and challenge 3. Rectal temperature was recorded for 180 min after each challenge.

For mechanistic studies of allergen (monomers versus SApNPs) trafficking, retention, presentation, and immunization-induced GC reactions in lymph nodes, adjuvanted allergen proteins were administered intradermally into the mouse footpads using a 29-gauge insulin needle under 3% isoflurane anesthesia with oxygen. Each mouse received 80 μL of an allergen/CpG adjuvant mixture, comprising 40 μg of allergen (10 μg per foot) and 40 μL of CpG adjuvant. Blood samples were collected from the maxillary/facial vein at designated time points. Serum was isolated by centrifugation at 13,200 × g for 10 min, heat-inactivated at 56 °C for 30 min, and clarified by an additional centrifugation at 4,300 × g for 10 min. Heat-inactivated serum was used for antigen-binding assays.

### Histology, immunostaining, and imaging for lymph node tissues

Allergen distribution, trafficking, retention, cellular interactions, and GC reactions in sentinel lymph nodes were evaluated in mice following intradermal immunization. Monomers and TIP60 SApNPs, each formulated with CpG adjuvant, were administered into four footpads. Mouse injection, tissue collection, and analytical procedures followed protocols established in our previous studies (*71, 74–77*). Each mouse received 80 μl of antigen–adjuvant mixture containing 40 μg antigen (10 μg per footpad). Sentinel lymph nodes were collected at time points ranging from 12 h to 6 weeks after a single–dose injection for immunohistological analysis. Lymph nodes were embedded in frozen section compound (VWR International, catalog no. 95057–838) in plastic cryomolds (Tissue–Tek, VWR catalog no. 4565), rapidly frozen in liquid nitrogen, and stored at −80 °C before transfer to the TSRI Core Facility for tissue processing, immunostaining, and imaging. Tissues were sectioned using a cryostat and mounted on charged slides. Sections were fixed in 10% neutral–buffered formalin and permeabilized in PBS containing 0.5% Triton X–100. Slides were blocked with Protein Block (Agilent) to minimize nonspecific antibody binding prior to immunostaining. Primary antibodies were applied and incubated overnight at 4 °C. After washing with TBST (TBS + 0.1% Tween–20), biotinylated or fluorophore–conjugated secondary antibodies were added and incubated for 1 h at 25 °C. Sections were stained with mAb148 (*93*) and mAb65 (*94*) at 1:100, followed by biotinylated goat anti–human IgG secondary antibody (Abcam, catalog no. ab7152, 1:300). Detection was performed using streptavidin–HRP (Vectastain Elite ABC–HRP Kit, Vector, catalog no. PK–6100) and diaminobenzidine (DAB) substrate (ImmPACT DAB, Vector, catalog no. SK–4105).

To examine interactions between allergens and cellular components in lymph nodes, specific cell populations were identified by immunostaining. FDCs were labeled using an anti–CD21 primary antibody (Abcam, cat. no. ab75985, 1:1800), followed by an anti–rabbit Alexa Fluor 555 secondary antibody (Thermo Fisher, cat. no. A21428, 1:200). Subcapsular sinus macrophages were detected with an anti–CD169 antibody (Abcam, cat. no. ab53443, 1:600), followed by an anti–rat Alexa Fluor 488 secondary antibody (Abcam, cat. no. ab150165, 1:200). B cells were labeled using an anti–B220 antibody (eBioscience, cat. no. 14–0452–82, 1:100), followed by an anti–rat Alexa Fluor 647 secondary antibody (Thermo Fisher, cat. no. A21247, 1:200). Allergen–induced GC reactions were assessed by staining GC B cells with a FITC–conjugated anti–GL7 antibody (BioLegend, cat. no. 144604, 1:250). Cell nuclei were counterstained with DAPI (Sigma–Aldrich, cat. no. D9542, 100 ng/ml). Stained lymph node sections were scanned using a Leica AT2 whole–slide scanner, and allergen trafficking and GC formation were quantified from bright–field and fluorescence images using ImageJ software (*134*).

### TEM analysis of allergen-presenting SApNPs in lymph node tissues

To assess interactions between allergens and cellular components in lymph nodes at both intracellular and intercellular levels, transmission electron microscopy (TEM) analysis was performed by the Core Microscopy Facility at TSRI, following procedures established in our previous studies (*71, 74, 75, 77*). To visualize TIP60 SApNPs in association with FDCs, B cells, and phagocytic cells, mice received 200 μl of an antigen–adjuvant mixture containing 100 μg allergen and 100 μl CpG, administered intradermally into the two hind footpads (50 μg per footpad). Fresh sentinel lymph nodes were collected at 12 h and 48 h after a single–dose injection. Lymph node tissues were bisected and fixed overnight at 4 °C in an oxygenated fixative containing 2.5% glutaraldehyde and 4% paraformaldehyde in 0.1 M sodium cacodylate buffer (pH 7.4). Samples were washed in the same buffer and post–fixed in 1% osmium tetroxide and 1.5% potassium ferrocyanide at 4 °C for 1 h. After rinsing with Corning Cell Culture Grade Water, tissues were stained with 0.5% uranyl acetate overnight at 4 °C. Samples were then rinsed with double–distilled water, dehydrated through a graded ethanol series followed by acetone, and infiltrated with LX–112 epoxy resin (Ladd), which was polymerized at 60 °C. Ultrathin sections (70 nm) were cut and mounted on copper grids for TEM imaging. Sections were imaged at 80 kV using a Talos L120C TEM (Thermo Fisher), and images were acquired with a CETA 16M CMOS camera for downstream analysis of nanoparticle–cell interactions.

### Lymph node disaggregation, cell staining, and flow cytometry

To assess allergen–induced humoral immunity, the frequencies and absolute numbers of GC B cells (B220⁺GL7⁺CD95⁺) and T_fh_ cells (CD3⁺CD4⁺CXCR5⁺PD–1⁺) were quantified by flow cytometry. Lymph node collection, processing, and analysis followed protocols established in our previous studies (*71, 74–77*). Each mouse received 80 μl of an antigen–adjuvant mixture containing 40 μg allergen (10 μg per footpad) formulated with CpG adjuvant, delivered intradermally into four footpads. Sentinel lymph nodes were isolated at 2, 4, and 6 weeks after a single–dose injection and mechanically disaggregated. Tissues were filtered through a 70 μm cell strainer to obtain single–cell suspensions, centrifuged at 400 × g for 10 min, and resuspended in HBSS (Thermo Fisher Scientific, cat. no. 14185052) supplemented with 0.5% bovine serum albumin and 2 mM EDTA. To block Fc receptor–mediated nonspecific binding, cells were incubated on ice for 30 min with anti–CD16/32 (BioLegend, cat. no. 101302, 1:50). Cells were transferred to 96–well V–bottom plates and stained on ice for 30 min with a premixed antibody cocktail containing: Zombie NIR live/dead stain (BioLegend, catalog no. 423106, 1:100), Brilliant Violet 510 anti-mouse/human CD45R/B220 antibody (BioLegend, catalog no. 103247, 1:300), FITC anti-mouse CD3 antibody (BioLegend, catalog no. 100204, 1:300), Alexa Fluor 700 anti-mouse CD4 antibody (BioLegend, catalog no. 100536, 1:300), PE anti-mouse/human GL7 antibody (BioLegend, catalog no. 144608, 1:500), Brilliant Violet 605 anti-mouse CD95 (Fas) antibody (BioLegend, catalog no. 152612, 1:500), Brilliant Violet 421 anti-mouse CD185 (CXCR5) antibody (BioLegend, catalog no. 145511, 1:500), and PE/Cyanine7 anti-mouse CD279 (PD-1) antibody (BioLegend, catalog no. 135216, 1:500). Following staining, cells were centrifuged at 400 × g for 10 min, washed with HBSS blocking solution, and fixed in 1.6% paraformaldehyde (Thermo Fisher Scientific, cat. no. 28906) for 30 min on ice. Fixed cells were washed, resuspended in HBSS blocking solution, and stored at 4 °C until acquisition. Flow cytometry was performed on a five–laser ZE5 cell analyzer (Yeti model; Bio–Rad) equipped with Everest software at the TSRI Core Facility. Data were analyzed using FlowJo v10, and results were plotted in GraphPad Prism, enabling quantification of GC B cell responses and Tfh cell dynamics across time points.

### Splenocytes isolation and T cell stimulation

Functional Arah2–specific T cell responses were assessed using activation–induced marker (AIM) and intracellular cytokine staining (ICS) flow cytometry assays (**Fig. S7**), following a protocol similar to our previous study (*76*). Each mouse received 80 μl of an antigen–adjuvant mixture containing 40 μg allergen (10 μg per footpad) formulated with CpG adjuvant, delivered intradermally into four footpads. Splenocytes were isolated 2 weeks after a single–dose injection. Spleens were mechanically disaggregated in PBS, passed through a 70 μm cell strainer to generate single–cell suspensions, and centrifuged at 400 × g for 5 min. Supernatants were discarded, and cell pellets were resuspended in ACK lysing buffer (Quality Biological, cat. no. 118–156–101) and incubated on ice for 5 min to remove red blood cells. Cells were washed with PBS, centrifuged at 400 × g for 5 min at 4 °C, and resuspended in 90% FBS and 10% EDTA. Splenocytes were cryopreserved in liquid nitrogen until downstream analysis. For stimulation assays, cryopreserved splenocytes were thawed rapidly and immediately diluted into 10 ml pre–warmed complete RPMI containing 10% heat–inactivated FBS and 1% penicillin–streptomycin to promote recovery and maintain viability. Cells were centrifuged at 400 × g for 10 min, resuspended in RPMI, counted, and adjusted to 10 million cells/ml. For each mouse, 1 million splenocytes were plated into 96–well U–bottom plates. Cells were stimulated with Arah2-His_6_ probe (1 μg/well) at 37 °C for 24 h. After 20 h, Brefeldin A (BioLegend, cat. no. 420601) was added to inhibit protein transport and enhance ICS signal. After an additional 4 h, cells were centrifuged (400 × g, 10 min) and resuspended in HBSS blocking buffer. Anti–CD16/32 (BioLegend, cat. no. 101302) was added for 30 min on ice to block Fc–mediated nonspecific binding. Cells were transferred to 96–well V–bottom plates and stained with a premixed surface–marker antibody cocktail, including: LIVE/DEAD Fixable Blue Dead Cell Stain Kit (Thermo Fisher Scientific, catalog no. L34962), FITC anti-mouse CD3 antibody (BioLegend, catalog no. 100204), Alexa Fluor 700 anti-mouse CD4 antibody (BioLegend, catalog no. 100536), BUV737 Anti-Mouse CD8a antibody (BD Bioscience, catalog no. 612759), APC anti-mouse CD154 antibody (BioLegend, catalog no. 106510), Brilliant Violet 421 anti-mouse CD69 Antibody (BioLegend, catalog no. 104527), APC/Fire 810 anti-mouse CD279 (PD-1) antibody (BioLegend, catalog no. 135251), and Brilliant Violet 605 anti-mouse CD185 (CXCR5) Antibody (BioLegend, catalog no. 145513). Cells were incubated with the cocktail on ice for 30 min, washed, and fixed in 1.6% paraformaldehyde (Thermo Fisher, cat. no. 28906) for 30 min on ice. Fixed cells were washed twice with intracellular staining permeabilization buffer (BioLegend, cat. no. 421002) and stained with the ICS antibody cocktail, including: PE anti-mouse IFN-γ antibody (BioLegend, catalog no. 505808), Brilliant Violet 785 anti-mouse TNF-α antibody (BioLegend, catalog no. 506341), PE/Cyanine5 anti-mouse IL-2 antibody (BioLegend, catalog no. 503824), PE/Cyanine7 anti-mouse IL-4 antibody (BioLegend, catalog no. 504118), and Brilliant Violet 711 anti-mouse IL-17A Antibody (BioLegend, catalog no. 506941). Cells were incubated on ice for 30 min, washed with permeabilization buffer, and stored in HBSS blocking buffer at 4 °C prior to acquisition. Sample events were collected on a Cytek Aurora spectral flow cytometer using SpectroFlo software at the TSRI Flow Cytometry Core. Data were analyzed in FlowJo v10, enabling quantification of AIM responses and cytokine–producing T cell subsets.

## Statistical analysis

Data were collected from 5 mice per group for immunization, sensitization, and challenge studies, and 3-5 mice per group for mechanistic analyses. All ELISA measurements were performed in duplicate. Comparisons among different allergen constructs and adjuvant–formulated allergens were analyzed using one–way ANOVA, followed by Tukey’s multiple–comparison post hoc test. For comparisons between two independent groups, two–tailed unpaired t-tests with Welch’s correction were applied. Statistical significance in figures is indicated as: ns (not significant), *p < 0.05, **p < 0.01, ***p < 0.001, and ****p < 0.0001. All graphs and statistical analyses were generated using GraphPad Prism 10.6.1 software.

## Data availability

All data needed to evaluate the conclusions of this research are available in the main text and Supplementary Information. The authors declare that the data supporting the findings of this study are available within the article and its Supplementary Information files. Source data are provided with this paper.

## Acknowledgements

We acknowledge K. Vanderpool, T. Fassel, and S. Henderson of the Microscopy Core Facility at TSRI for their expert assistance with negative-stain EM analysis. We thank M. Livneh, K. Spencer, and S. Henderson of the Histology and Microscopy Core Facility at TSRI for their expertise and technical support in immunohistology. We acknowledge A. Saluk, B. Seegers, and B. Monteverde of the Flow Cytometry Core Facility at TSRI for their technical support.

## Author contributions

Project design by Y.-N.Z, L.H., and J.Z.; hTIP60-allergen SApNP design and hTIP60 expansion by L.H. and J.Z.; bioinformatics analysis, cf1/2TIP60 design, and structural optimization by L.H. and J.Z.; evolution-inspired cf2TIP60-Q-NPE design by L.H. and J.Z.; expression and purification of antibodies, allergens, hTIP60 SApNP variants, and cf1/2TIP60 SApNP variants by L.H.; BN-PAGE by C.D.; SEC, DSC, and DLS by L.H.; negative-stain EM by L.H. and J.Z.; mouse immunization and peanut sensitization–challenge by Y.-N.Z.; ELISA by Y.-N.Z.; lymph node analysis by Y.-N.Z.; flow cytometry and T cell analysis by Y.-N.Z.; manuscript written by Y.-N.Z., L.H., and J.Z. All authors were asked to comment on the manuscript.

## References

1. J. A. Lieberman et al., The global burden of illness of peanut allergy: A comprehensive literature review. Allergy 76, 1367–1384 (2021).

2. S. H. Sicherer, H. A. Sampson, Peanut allergy: Emerging concepts and approaches for an apparent epidemic. J. Allergy Clin. Immunol. 120, 491–503 (2007).

3. Z. Husain, R. A. Schwartz, Peanut allergy: An increasingly common life-threatening disorder. J. Am. Acad. Dermatol. 66, 136–143 (2012).

4. F. D. Finkelman, Peanut allergy and anaphylaxis. Curr. Opin. Immunol. 22, 783–788 (2010).

5. R. Czolk et al., IgE-mediated peanut allergy: Current and novel predictive biomarkers for clinical phenotypes using multi-omics approaches. Front. Immunol. 11, 594350 (2020).

6. J. Wang et al., Pathogenesis of allergic diseases and implications for therapeutic interventions. Signal Transduct. Target. Ther. 8, 138 (2023).

7. I. Ogulur et al., Type 2 immunity in allergic diseases. Cell. Mol. Immunol. 22, 211–242 (2025).

8. G. Lack, D. Fox, K. Northstone, J. Golding, C. Avon Longitudinal Study Parents, Factors associated with the development of peanut allergy in childhood. New Engl. J. Med. 348, 977–985 (2003).

9. S. Barni et al., Immunoglobulin E (IgE)-mediated food allergy in children: Epidemiology, pathogenesis, diagnosis, prevention, and management. Medicina (Kaunas) 56, 111 (2020).

10. S. H. Sicherer, H. A. Sampson, Food allergy: Epidemiology, pathogenesis, diagnosis, and treatment. J. Allergy Clin. Immunol. 133, 291–307 (2014).

11. R. S. Gupta et al., The prevalence, severity, and distribution of childhood food allergy in the United States. Pediatrics 128, e9–17 (2011).

12. P. J. Turner et al., Fatal anaphylaxis: Mortality rate and risk factors. J. Allergy. Clin. Immunol. Pract. 5, 1169–1178 (2017).

13. S. A. Bock, A. Muñoz-Furlong, H. A. Sampson, Fatalities due to anaphylactic reactions to foods. J. Allergy Clin. Immunol. 107, 191–193 (2001).

14. H. A. Sampson, Peanut Allergy. New Engl. J. Med. 346, 1294–1299 (2002).

15. R. J. Klemans et al., Ara h 2 is the best predictor for peanut allergy in adults. J. Allergy. Clin. Immunol. Pract. 1, 632–638.e1 (2013).

16. O. Hemmings, G. Du Toit, S. Radulovic, G. Lack, A. F. Santos, Ara h 2 is the dominant peanut allergen despite similarities with Ara h 6. J. Allergy Clin. Immunol. 146, 621–630.e5 (2020).

17. A. E. Koid et al., Ara h 6 complements Ara h 2 as an important marker for IgE reactivity to peanut. J. Agric. Food Chem. 62, 206–213 (2014).

18. H. J. M. Warmenhoven et al., IgE cross-inhibition between Ara h 1 and Ara h 2 is explained by complex formation of both major peanut allergens. J. Allergy Clin. Immunol. 152, 436–444.e6 (2023).

19. X. N. Chen, Q. Wang, R. El-Mezayen, Y. H. Zhuang, S. C. Dreskin, Ara h 2 and Ara h 6 have similar allergenic activity and are substantially redundant. Int. Arch. Allergy Immunol. 160, 251–258 (2013).

20. A. Long, The nuts and bolts of peanut allergy. *New Engl*. J. Med. 346, 1320–1322 (2002).

21. M. R. Datema et al., Estimating the risk of severe peanut allergy using clinical background and IgE sensitization profiles. *Front*. Allergy 2, 670789 (2021).

22. S. J. Galli, M. Tsai, IgE and mast cells in allergic disease. Nat. Med. 18, 693–704 (2012).

23. C. Kanagaratham, Y. S. El Ansari, O. L. Lewis, H. C. Oettgen, IgE and IgG antibodies as regulators of mast cell and basophil functions in food allergy. Front. Immunol. 11, 603050 (2020).

24. S. J. Galli, M. Tsai, A. M. Piliponsky, The development of allergic inflammation. Nature 454, 445–454 (2008).

25. B. P. Vickery, M. Ebisawa, W. G. Shreffler, R. A. Wood, Current and future treatment of peanut allergy. J. Allergy. Clin. Immunol. Pract. 7, 357–365 (2019).

26. J. Z. X. Lee et al., Next-generation allergen-specific immunotherapy for food allergy. Clin. Rev. Allergy Immunol. 68, 93 (2025).

27. S. R. Durham, M. H. Shamji, Allergen immunotherapy: Past, present and future. Nat. Rev. Immunol. 23, 317–328 (2023).

28. A. M. Schoos et al., Immunological outcomes of allergen-specific immunotherapy in food allergy. Front. Immunol. 11, 568598 (2020).

29. Y. H. Jeon, E. H. Kim, Recent advances in food allergen immunotherapy. Clin. Exp. Pediatr. 67, 386–394 (2024).

30. B. P. Vickery et al., Sustained unresponsiveness to peanut in subjects who have completed peanut oral immunotherapy. J. Allergy Clin. Immunol. 133, 468–475 (2014).

31. A. T. Clark et al., Successful oral tolerance induction in severe peanut allergy. Allergy 64, 1218–1220 (2009).

32. A. M. Scurlock, S. M. Jones, An update on immunotherapy for food allergy. Curr. Opin. Allergy Clin. Immunol. 10, 587–593 (2010).

33. E. H. Kim, A. W. Burks, Food allergy immunotherapy: Oral immunotherapy and epicutaneous immunotherapy. Allergy 75, 1337–1346 (2020).

34. S. M. Jones et al., Clinical efficacy and immune regulation with peanut oral immunotherapy. J. Allergy Clin. Immunol. 124, 292–300 (2009).

35. S. M. Jones et al., Epicutaneous immunotherapy for the treatment of peanut allergy in children and young adults. J. Allergy Clin. Immunol. 139, 1242–1252.e9 (2017).

36. M. Ravindran et al., Epicutaneous immunotherapy for the treatment of peanut allergy. Allergy 80, 63–76 (2025).

37. E. H. Kim et al., Sublingual immunotherapy for peanut allergy: Clinical and immunologic evidence of desensitization. J. Allergy Clin. Immunol. 127, 640–646.e1 (2011).

38. E. H. Kim et al., Long-term sublingual immunotherapy for peanut allergy in children: Clinical and immunologic evidence of desensitization. J. Allergy Clin. Immunol. 144, 1320–1326.e1 (2019).

39. J. E. M. Upton et al., Safety and efficacy of very low-dose multi-nut oral immunotherapy in children. Clin. Transl. Allergy 15, e70125 (2025).

40. K. Blumchen et al., Efficacy, safety, and quality of life in a multicenter, randomized, placebo-controlled trial of low-dose peanut oral immunotherapy in children with peanut allergy. J. Allergy. Clin. Immunol. Pract. 7, 479–491.e10 (2019).

41. P. Carlucci, F. Spataro, M. F. Daddato, G. Paoletti, D. Di Bona, Biologic drugs and allergen immunotherapy: Potential allies. Explor. Asthma Allergy 1, 126–141 (2023).

42. B. Olivieri, F. E. Gürounaydin, J. Corren, G. Senna, S. R. Durham, The combination of allergen immunotherapy and biologics for inhalant allergies: Exploring the synergy. *Ann. Allergy*, Asthma Immunol. 134, 385–395 (2025).

43. R. A. Wood et al., Omalizumab for the treatment of multiple food allergies. *New Engl*. J. Med. 390, 889–899 (2024).

44. T. Zuberbier et al., Omalizumab in IgE-mediated food allergy: A systematic review and meta-analysis. J. Allergy. Clin. Immunol. Pract. 11, 1134–1146 (2023).

45. U. M. Sahiner et al., Mechanisms of allergen immunotherapy and potential biomarkers for clinical evaluation. J. Pers. Med. 13, 845 (2023).

46. D. de Silva et al., Allergen immunotherapy and/or biologicals for IgE-mediated food allergy: A systematic review and meta-analysis. Allergy 77, 1852–1862 (2022).

47. M. Lao-Araya, Novel approaches to allergen immunotherapy for respiratory allergies. Pharmaceuticals 17, 1510 (2024).

48. M. C. Berin, Future therapies for IgE-mediated food allergy. Curr. Pediatr. Rep. 2, 119–126 (2014).

49. Y. Zheng, J. L. Subiza, Vaccines and adjuvants in allergen specific immunotherapy: Current status and future directions. Allergy Med. 6, 100065 (2025).

50. J. M. Smeekens et al., Immunization with virus-like particles displaying Ara h 2-derived peptides to mitigate peanut allergy. ACS Nano Med. 1, 1987–1996 (2026).

51. A. Jacquet, A. Fettelschoss-Gabriel, P. Johansen, New generation of allergen immunotherapy using virus-like particles: A comprehensive review. Clin. Exp. Allergy 56, 597–612 (2026).

52. M. F. Bachmann, G. T. Jennings, Vaccine delivery: A matter of size, geometry, kinetics and molecular patterns. Nat. Rev. Immunol. 10, 787–796 (2010).

53. S. Nooraei et al., Virus-like particles: Preparation, immunogenicity and their roles as nanovaccines and drug nanocarriers. J. Nanobiotechnol. 19, 59 (2021).

54. M. O. Mohsen, M. F. Bachmann, Virus-like particle vaccinology, from bench to bedside. Cell. Mol. Immunol. 19, 993–1011 (2022).

55. E. M. Plummer, M. Manchester, Viral nanoparticles and virus-like particles: Platforms for contemporary vaccine design. Wiley Interdiscip. Rev. Nanomed. Nanobiotechnol. 3, 174–196 (2010).

56. H. Berreiros-Hortala, G. Vilchez-Pinto, A. Diaz-Perales, M. Garrido-Arandia, J. Tome-Amat, Virus-like particles as vaccines for allergen-specific therapy: An overview of current developments. Int. J. Mol. Sci. 25, 7429 (2024).

57. F. Storni et al., Vaccine against peanut allergy based on engineered virus-like particles displaying single major peanut allergens. J. Allergy Clin. Immunol. 145, 1240–1253.e3 (2020).

58. P. Engeroff et al., Allergens displayed on virus-like particles are highly immunogenic but fail to activate human mast cells. Allergy 73, 341–349 (2018).

59. J. M. Sobczak et al., The next generation virus-like particle platform for the treatment of peanut allergy. Allergy 78, 1980–1996 (2023).

60. Z. Gharailoo et al., Generation of a virus-like particles based vaccine against IgE. Allergy 79, 2207–2221 (2024).

61. M. F. Bachmann et al., On the role of antibody affinity and avidity in the IgE-mediated allergic response. Allergy 80, 37–46 (2025).

62. J. A. Layhadi et al., Ara h 2-expressing cucumber mosaic virus-like particle (VLP Peanut) induces in vitro tolerogenic cellular responses in peanut-allergic individuals. J. Allergy Clin. Immunol. 155, 153–165 (2025).

63. G. T. Jennings, M. F. Bachmann, Immunodrugs: Therapeutic VLP-based vaccines for chronic diseases. Annu. Rev. Pharmacol. Toxicol. 49, 303–326 (2009).

64. P. S. Krenger et al., Influence of antigen density and TLR ligands on preclinical efficacy of a VLP-based vaccine against peanut allergy. Allergy 79, 184–199 (2024).

65. F. Heil et al., Species-specific recognition of single-stranded RNA via toll-like receptor 7 and 8. Science 303, 1526–1529 (2004).

66. B. Nguyen, N. H. Tolia, Protein-based antigen presentation platforms for nanoparticle vaccines. npj Vaccines 6, 70 (2021).

67. L. He et al., HIV-1 vaccine design through minimizing envelope metastability. Sci. Adv. 4, eaau6769 (2018).

68. L. He et al., Presenting native-like trimeric HIV-1 antigens with self-assembling nanoparticles. Nat. Commun. 7, 12041 (2016).

69. L. He et al., Proof of concept for rational design of hepatitis C virus E2 core nanoparticle vaccines. Sci. Adv. 6, eaaz6225 (2020).

70. L. He et al., Single-component multilayered self-assembling nanoparticles presenting rationally designed glycoprotein trimers as Ebola virus vaccines. Nat. Commun. 12, 2633 (2021).

71. Y.-N. Zhang et al., Single-component multilayered self-assembling protein nanoparticles presenting glycan-trimmed uncleaved prefusion optimized envelope trimmers as HIV-1 vaccine candidates. Nat. Commun. 14, 1985 (2023).

72. L. He et al., Single-component, self-assembling, protien nanoparticles presenting the receptor binding domain and stabilized spike as SARS-CoV-2 vaccine candidates. Sci. Adv. 7, eabf1591 (2021).

73. L. He et al., Native-like soluble E1E2 glycoprotein heterodimers on self-assembling protein nanoparticles for hepatitis C virus vaccine design. Nat. Commun. 17, 2633 (2026).

74. Y.-Z. Lee et al., Rational design of next-generation filovirus vaccines combining glycoprotein stabilization and nanoparticle display with glycan modification. Nat. Commun. 16, 11368 (2025).

75. Y.-N. Zhang et al., Mechanism of a COVID-19 nanoparticle vaccine candidate that elicits a broadly neutralizing antibody response to SARS-CoV-2 variants. Sci. Adv. 7, eabj3107 (2021).

76. K. B. Gomes et al., Single-component multilayered self-assembling protein nanoparticles displaying extracellular domains of matrix protein 2 as a pan-influenza A vaccine. ACS Nano 17, 23545–23567 (2023).

77. Y.-N. Zhang et al., Single-component self-assembling protein nanoparticles displaying stabilized prefusion-closed hemagglutinin trimers for influenza vaccine development. Nat. Commun. 17, 9717 (2026).

78. Y.-N. Zhang et al., A single-component multilayered self-assembling protein nanoparticle vaccine based on extracellular domains of matrix protein 2 against both influenza A and B. Vaccines 12, 975 (2024).

79. J. Obata et al., Icosahedral 60-meric porous structure of designed supramolecular protein nanoparticle TIP60. Chem. Commun. 57, 10226–10229 (2021).

80. M. Yamashita et al., misteINK: A protein nanocage-based ink with reversible, stimuli-responsive color shifts. Biomater. Sci. 13, 6652–6661 (2025).

81. N. Ohara et al., Reversible assembly of an artificial protein nanocage using alkaline earth metal ions. J. Am. Chem. Soc. 145, 216–223 (2023).

82. Q. Lu, F. Ye, Z. Wei, Z. Wen, M. Zhang, Antiparallel coiled-coil-mediated dimerization of myosin X. Proc. Natl. Acad. Sci. U. S. A. 109, 17388–17393 (2012).

83. X. Chen et al., Conformational IgE epitopes of peanut allergens Ara h 2 and Ara h 6. Clin. Exp. Allergy 46, 1120–1128 (2016).

84. K. Otsu, R. Guo, S. C. Dreskin, Epitope analysis of Ara h 2 and Ara h 6: Characteristic patterns of IgE-binding fingerprints among individuals with similar clinical histories. Clin. Exp. Allergy 45, 471–484 (2015).

85. K. Lehmann et al., Structure and stability of 2S albumin-type peanut allergens: Implications for the severity of peanut allergic reactions. Biochem. J. 395, 463–472 (2006).

86. H. S. Porterfield et al., Effector activity of peanut allergens: A critical role for Ara h 2, Ara h 6, and their variants. Clin. Exp. Allergy 39, 1099–1108 (2009).

87. D. Das et al., Crystal structure of a novel Sm-like protein of putative cyanophage origin at 2.60 Å resolution. Proteins 75, 296–307 (2009).

88. S. A. Smith et al., Structural determinants of peanut-induced anaphylaxis. J. Allergy Clin. Immunol. 155, 1547–1556.e3 (2025).

89. N. A. LaHood et al., Immunotherapy-induced neutralizing antibodies disrupt allergen binding and sustain allergen tolerance in peanut allergy. J. Clin. Invest. 133, e164501 (2023).

90. J. M. Smeekens, M. D. Kulis, Mouse models of food allergy in the pursuit of novel treatment modalities. *Front*. Allergy 2, 810067 (2021).

91. L. Wagenaar et al., Mouse strain differences in response to oral immunotherapy for peanut allergy. Immun. Inflamm. Dis. 7, 41–51 (2019).

92. S. F. Altschul et al., Gapped BLAST and PSI-BLAST: A new generation of protein database search programs. Nucleic Acids Res. 25, 3389–3402 (1997).

93. K. J. Cho et al., Structure of the Extracellular Domain of Matrix Protein 2 of Influenza A Virus in Complex with a Protective Monoclonal Antibody. J. Virol. 89, 3700–3711 (2015).

94. K. J. Cho et al., Crystal Structure of the Conserved Amino Terminus of the Extracellular Domain of Matrix Protein 2 of Influenza A Virus Gripped by an Antibody. J. Virol. 90, 611–615 (2016).

95. R. Rappuoli, Glycoconjugate vaccines: Principles and mechanisms. Sci. Transl. Med. 10, eaat4615 (2018).

96. J. G. Cyster, B cell follicles and antigen encounters of the third kind. Nat. Immunol. 11, 989–996 (2010).

97. B. A. Heesters, R. C. Myers, M. C. Carroll, Follicular dendritic cells: Dynamic antigen libraries. Nat. Rev. Immunol. 14, 495–504 (2014).

98. M. E. El Shikh, R. M. El Sayed, S. Sukumar, A. K. Szakal, J. G. Tew, Activation of B cells by antigens on follicular dendritic cells. Trends Immunol. 31, 205–211 (2010).

99. Y.-N. Zhang et al., Nanoparticle size influences antigen retention and presentation in lymph node follicles for humoral immunity. Nano Lett. 19, 7226–7235 (2019).

100. Y.-N. Zhang, W. Poon, E. Sefton, W. C. W. Chan, Suppressing subcapsular sinus macrophages enhances transport of nanovaccines to lymph node follicles for robust humoral immunity. ACS Nano 14, 9478–9490 (2020).

101. Y. R. Carrasco, F. D. Batista, B cells acquire particulate antigen in a macrophage-rich area at the boundary between the follicle and the subcapsular sinus of the lymph node. Immunity 27, 160–171 (2007).

102. T. Junt et al., Subcapsular sinus macrophages in lymph nodes clear lymph-borne viruses and present them to antiviral B cells. Nature 450, 110–114 (2007).

103. T. G. Phan, J. A. Green, E. E. Gray, Y. Xu, J. G. Cyster, Immune complex relay by subcapsular sinus macrophages and noncognate B cells drives antibody affinity maturation. Nat. Immunol. 10, 786–793 (2009).

104. S. F. Gonzalez et al., Capture of influenza by medullary dendritic cells via SIGN-R1 is essential for humoral immunity in draining lymph nodes. Nat. Immunol. 11, 427–434 (2010).

105. J. G. Cyster, P. C. Wilson, Antibody modulation of B cell responses-Incorporating positive and negative feedback. Immunity 57, 1466–1481 (2024).

106. G. D. Victora, M. C. Nussenzweig, Germinal centers. Annu. Rev. Immunol. 30, 429–457 (2012).

107. C. D. Allen, T. Okada, J. G. Cyster, Germinal-center organization and cellular dynamics. Immunity 27, 190–202 (2007).

108. S. Crotty, T follicular helper cell differentiation, function, and roles in disease. Immunity 41, 529–542 (2014).

109. Y. Tang et al., Antigen-specific effector CD8 T cells regulate allergic responses via IFN-γ and dendritic cell function. J. Allergy Clin. Immunol. 129, 1611–1620.e4 (2012).

110. V. Ropars et al., The myosin X motor is optimized for movement on actin bundles. Nat. Commun. 7, 12456 (2016).

111. S. K. Duraisamy, I. K. Sundar, Sex-specific leukocyte responses to peanut allergen: Uncovering potential links to the molecular circadian clock. J. Leukocyte Biol. 117, qiaf097 (2025).

112. S. Rajput, M. S. Vininski, L. A. Lehmann, N. J. Hobbs, J. J. Dolence, Androgen receptor signaling protects male mice from the development of immune response to peanut. Am. J. Clin. Exp. Immunol. 12, 60–71 (2023).

113. S. E. Dunn, W. A. Perry, S. L. Klein, Mechanisms and consequences of sex differences in immune responses. Nat. Rev. Nephrol. 20, 37–55 (2024).

114. P. Pechsrichuang, S. Namwongnao, A. Jacquet, Bioengineering of virus-like particles for the prevention or treatment of allergic diseases. Allergy Asthma Immunol. Res. 13, 23–41 (2021).

115. C. T. Cady et al., IgG antibodies produced during subcutaneous allergen immunotherapy mediate inhibition of basophil activation via a mechanism involving both FcγRIIA and FcγRIIB. Immunol. Lett. 130, 57–65 (2010).

116. R. T. Strait, S. C. Morris, F. D. Finkelman, IgG-blocking antibodies inhibit IgE-mediated anaphylaxis in vivo through both antigen interception and FcγRIIb cross-linking. J. Clin. Invest. 116, 833–841 (2006).

117. M. R. Strobl et al., The role of IgG1 and IgG4 as dominant IgE-blocking antibodies shifts during allergen immunotherapy. J. Allergy Clin. Immunol. 151, 1371–1378.e5 (2023).

118. J. López-Sagaseta, E. Malito, R. Rappuoli, M. J. Bottomley, Self-assembling protein nanoparticles in the design of vaccines. Comput. Struct. Biotechnol. J. 14, 58–68 (2016).

119. B. Pulendran, R. Ahmed, Immunological mechanisms of vaccination. Nat. Immunol. 12, 509–517 (2011).

120. B. Pulendran, R. Ahmed, Translating innate immunity into immunological memory: Implications for vaccine development. Cell 124, 849–863 (2006).

121. K. A. Fitzgerald, J. C. Kagan, Toll-like Receptors and the Control of Immunity. Cell 180, 1044–1066 (2020).

122. T. Petek et al., Clinical and immunological markers of peanut allergy severity and tolerance in children. *Front*. Allergy 7, 1826635 (2026).

123. N. Singh et al., Peanut component-specific IgE and IgG4 are modulated by sublingual immunotherapy and are associated with remission in 1-4-year-old children. Allergy 81, 2089–2102 (2026).

124. I. Bartha, N. Almulhem, A. F. Santos, Feast for thought: A comprehensive review of food allergy 2021-2023. J. Allergy Clin. Immunol. 153, 576–594 (2024).

125. C. A. Akdis et al., Allergy: Mechanistic insights into new methods of prevention and therapy. Sci. Transl. Med. 15, eadd2563 (2023).

126. D. Petrey et al., Using multiple structure alignments, fast model building, and energetic analysis in fold recognition and homology modeling. Proteins 53, 430–435 (2003).

127. E. F. Pettersen et al., UCSF chimera - A visualization system for exploratory research and analysis. J. Comput. Chem. 25, 1605–1612 (2004).

128. Z. X. Xiang, C. S. Soto, B. Honig, Evaluating conformational free energies: The colony energy and its application to the problem of loop prediction. Proc. Natl. Acad. Sci. U. S. A. 99, 7432–7437 (2002).

129. M. A. Larkin et al., Clustal W and clustal X version 2.0. Bioinformatics 23, 2947–2948 (2007).

130. N. Saitou, M. Nei, The neighbor-joining method: A new method for reconstructing phylogenetic trees. Mol. Biol. Evol. 4, 406–425 (1987).

131. M. P. Jacobson, R. A. Friesner, Z. X. Xiang, B. Honig, On the role of the crystal environment in determining protein side-chain conformations. J. Mol. Biol. 320, 597–608 (2002).

132. Z. X. Xiang, B. Honig, Extending the accuracy limits of prediction for side-chain conformations. J. Mol. Biol. 311, 421–430 (2001).

133. J. K. Min et al., Design of an Ara h 2 hypoallergen from conformational epitopes. Clin. Exp. Allergy 54, 46–55 (2024).

134. C. A. Schneider, W. S. Rasband, K. W. Eliceiri, NIH Image to ImageJ: 25 years of image analysis. Nat. Methods 9, 671–675 (2012).

