## Supplementary Information for "Rationally Engineered Self-Assembling Protein Nanoparticles for Allergen-Specific Peanut Allergy Immunotherapy"

**Fig. S1**

a

**>Arah2-His$_6$**

MGILPSPGMPALLSLVSLLSVLLMGCVAEARQQWELQGDRRCQSQLERANLRPCEQHLMQKIQRDEDSYERDPYSPSQDPYSPSPYDRRGAGSSQHQERC
CNELNEFENNQRCMCEALQQIMENQSDRLQGRQQEQQFKRELRNLPQQCGLRAPQRCDLDVESGGASGS**HHHHHH**

**>Arah6-His$_6$**

MGILPSPGMPALLSLVSLLSVLLMGCVAEAMRRERGRQGDSSSCERQVDGVNLKPCEQHIMQRIMGEQEQYDSYNFGSTRSSDQQQRCCDELNEMENTQR
CMCEALQQIMENQCDGLQDRQMVQHFKRELMNLPQQCNFGAPQRCDLDVSGGRCASGS**HHHHHH**

**>hTIP60-Arah2**

MGILPSPGMPALLSLVSLLSVLLMGCVAESQDPKNIKIMRLVTGEDIIGNISESQGLITIKKAFVIIPMQATPGKPVQLVLSPWQPYTDDKEIVIDDSKV
ITITSPKDDIIKSYESHTRVLENKQVEEILRLEKEIEDLQRMKEQQELSLTEASLQKLQERRDQELRRLEEEASGGGGSARQQWELQGDRRCQSQLERAN
LRPCEQHLMQKIQRDEDSYERDPYSPSQDPYSPSPYDRRGAGSSQHQERCCNELNEFENNQRCMCEALQQIMENQSDRLQGRQQEQQFKRELRNLPQQCG
LRAPQRCDLDVESGG

**>hTIP60-Arah6-Arah2**

MGILPSPGMPALLSLVSLLSVLLMGCVAESQDPKNIKIMRLVTGEDIIGNISESQGLITIKKAFVIIPMQATPGKPVQLVLSPWQPYTDDKEIVIDDSKV
ITITSPKDDIIKSYESHTRVLENKQVEEILRLEKEIEDLQRMKEQQELSLTEASLQKLQERRDQELRRLEEEASGGGGSAMRRERGRQGDSSSCERQVDG
VNLKPCEQHIMQRIMGEQEQYDSYNFGSTRSSDQQQRCCDELNEMENTQRCMCEALQQIMENQCDGLQDRQMVQHFKRELMNLPQQCNFGAPQRCDLDVS
GGRCGGGGSARQQWELQGDRRCQSQLERANLRPCEQHLMQKIQRDEDSYERDPYSPSQDPYSPSPYDRRGAGSSQHQERCCNELNEFENNQRCMCEALQQ
IMENQSDRLQGRQQEQQFKRELRNLPQQCGLRAPQRCDLDVESGG

**>hTIP60-Ext-Arah2**

MGILPSPGMPALLSLVSLLSVLLMGCVAESQDPKNIKIMRLVTGEDIIGNISESQGLITIKKAFVIIPMQATPGKPVQLVLSPWQPYTDDKEIVIDDSKV
ITITSPKDDIIKSYESHTRVLEAAKAAAENKQVEEILRLEKEIEDLQRMKEQQELSLTEASLQKLQERRDQELRRLEEEASGGGGSARQQWELQGDRRCQ
SQLERANLRPCEQHLMQKIQRDEDSYERDPYSPSQDPYSPSPYDRRGAGSSQHQERCCNELNEFENNQRCMCEALQQIMENQSDRLQGRQQEQQFKRELR
NLPQQCGLRAPQRCDLDVESGG

**>hTIP60-Ext-Arah6-Arah2**

MGILPSPGMPALLSLVSLLSVLLMGCVAESQDPKNIKIMRLVTGEDIIGNISESQGLITIKKAFVIIPMQATPGKPVQLVLSPWQPYTDDKEIVIDDSKV
ITITSPKDDIIKSYESHTRVLEAAKAAAENKQVEEILRLEKEIEDLQRMKEQQELSLTEASLQKLQERRDQELRRLEEEASGGGGSAMRRERGRQGDSSS
CERQVDGVNLKPCEQHIMQRIMGEQEQYDSYNFGSTRSSDQQQRCCDELNEMENTQRCMCEALQQIMENQCDGLQDRQMVQHFKRELMNLPQQCNFGAPQ
RCDLDVSGGRCGGGGSARQQWELQGDRRCQSQLERANLRPCEQHLMQKIQRDEDSYERDPYSPSQDPYSPSPYDRRGAGSSQHQERCCNELNEFENNQRC
MCEALQQIMENQSDRLQGRQQEQQFKRELRNLPQQCGLRAPQRCDLDVESGG

: Signal peptide
: Arah2; : Arah6;
Sm-hMyoX : hTIP60; Sm Ext hMyoX : hTIP60-Ext
AS: Enzymatic site; : Linker
**HHHHHH**: His$_6$ tag

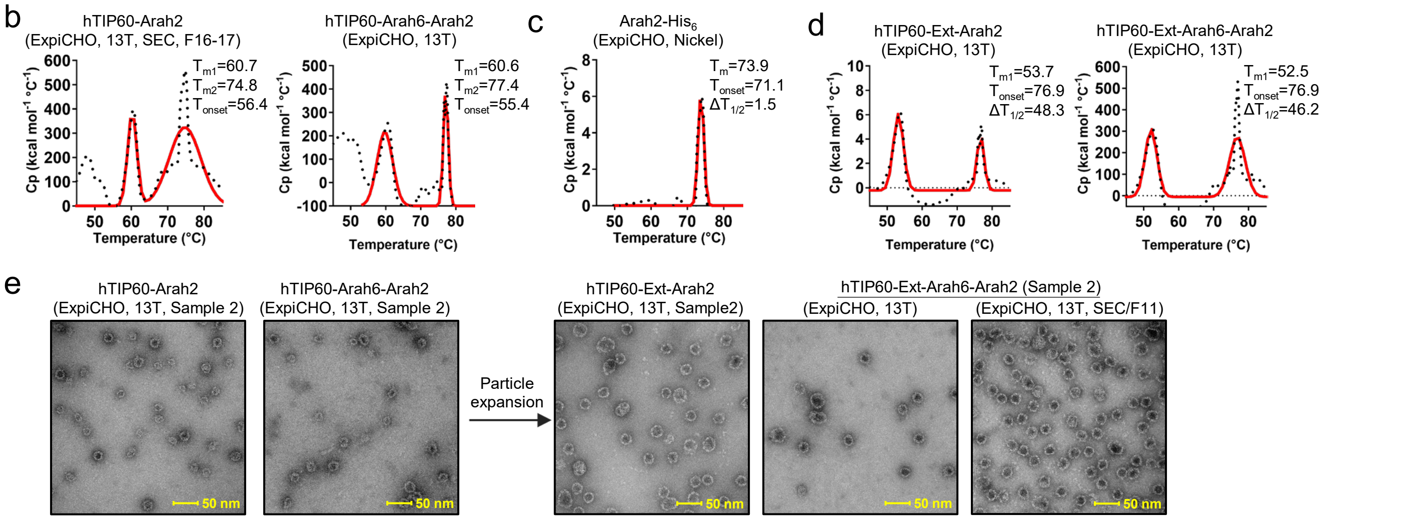


**Fig. S1. In vitro characterization of human TIP60 (hTIP60) constructs displaying peanut allergens**. (**a**) Amino acid sequences of peanut allergen constructs, including Arah2-His$_6$, Arah6-His$_6$, hTIP60 nanoparticles displaying Arah2 and tandem Arah6-Arah2, and expanded hTIP60 (hTIP60-Ext) displaying Arah2 and tandem Arah6-Arah2. Signal peptide, Arah2, Arah6, hTIP60, 7-residue expansion segment (Ext), enzymatic site, and linkers are highlighted in yellow, brown, green, dark moss green, cyan, magenta, and gray, respectively. (**b–d**) DSC thermograms of ExpiCHO-expressed constructs: (**b**) hTIP60-Arah2 and hTIP60-Arah6-Arah2 purified by immunoaffinity chromatography (IAC) using a 13T antibody column; (**c**) soluble Arah2-His$_6$ monomer purified by nickel affinity chromatography; and (**d**) hTIP60-Ext-Arah2 and hTIP60-Ext-Arah6-Arah2 purified by IAC. Experimental data (black points) and Gaussian fits (red lines) are shown, with melting temperature(s) ($T_m$ or $T_{m1}$/$T_{m2}$) and onset temperature ($T_{onset}$) indicated. (**e**) Negative-stain EM micrographs of hTIP60-Arah2/hTIP60-Arah6-Arah2 (left) and hTIP60-Ext-Arah2/hTIP60-Ext-Arah6-Arah2 (right) from an independent production run (sample 2). A representative nsEM micrograph of the first SEC peak of hTIP60-Ext-Arah6-Arah2 (far right) is shown. Scale bar, 50 nm.

**Fig. S2**

a

**>cf1TIP60-Arah2**
MGILPSPGMPALLSLVSLLSVLLMGCVAESQDPKNIKIMRLVTGEDIIGNISESQGLITIKKAFVIIPMQATPGKPVQLVLSPWQPYTDDKEIVIDDSKV
ITITSPKDDIIKSYESHTRVLETRQMEEILRLEREIERLQRQRDEGVALMNESTQEELRQLHDAEIYRLEKEASGGGGSARQQWELQGDRRCQSQLERAN
LRPCEQHLMQKIQRDEDSYERDPYSPSQDPYSPSPYDRRGAGSSQHQERCCNELNEFENNQRCMCEALQQIMENQSDRLQGRQQEQQFKRELRNLPQQCG
LRAPQRCDLDVESGG

**>cf1TIP60-Arah6-Arah2**
MGILPSPGMPALLSLVSLLSVLLMGCVAESQDPKNIKIMRLVTGEDIIGNISESQGLITIKKAFVIIPMQATPGKPVQLVLSPWQPYTDDKEIVIDDSKV
ITITSPKDDIIKSYESHTRVLETRQMEEILRLEREIERLQRQRDEGVALMNESTQEELRQLHDAEIYRLEKEASGGGGSAMRRERGRQGDSSSCERQVDG
VNLKPCEQHIMQRIMGEQEQYDSYNFGSTRSSDQQQRCCDELNEMENTQRCMCEALQQIMENQCDGLQDRQMVQHFKRELMNLPQQCNFGAPQRCDLDVS
GGRCGGGGSARQQWELQGDRRCQSQLERANLRPCEQHLMQKIQRDEDSYERDPYSPSQDPYSPSPYDRRGAGSSQHQERCCNELNEFENNQRCMCEALQQ
IMENQSDRLQGRQQEQQFKRELRNLPQQCGLRAPQRCDLDVESGG

**>cf2TIP60-Arah2**
MGILPSPGMPALLSLVSLLSVLLMGCVAESQDPKNIKIMRLVTGEDIIGNISESQGLITIKKAFVIIPMQATPGKPVQLVLSPWQPYTDDKEIVIDDSKV
ITITSPKDDIIKSYESHTRVLETRQMEKILALEREIERLQRQHDEGLALMNEGSREELRQRRDAEIYRLEKEASGGGGSARQQWELQGDRRCQSQLERAN
LRPCEQHLMQKIQRDEDSYERDPYSPSQDPYSPSPYDRRGAGSSQHQERCCNELNEFENNQRCMCEALQQIMENQSDRLQGRQQEQQFKRELRNLPQQCG
LRAPQRCDLDVESGG

**>cf2TIP60-Arah6-Arah2**
MGILPSPGMPALLSLVSLLSVLLMGCVAESQDPKNIKIMRLVTGEDIIGNISESQGLITIKKAFVIIPMQATPGKPVQLVLSPWQPYTDDKEIVIDDSKV
ITITSPKDDIIKSYESHTRVLETRQMEKILALEREIERLQRQHDEGLALMNEGSREELRQRRDAEIYRLEKEASGGGGSAMRRERGRQGDSSSCERQVDG
VNLKPCEQHIMQRIMGEQEQYDSYNFGSTRSSDQQQRCCDELNEMENTQRCMCEALQQIMENQCDGLQDRQMVQHFKRELMNLPQQCNFGAPQRCDLDVS
GGRCGGGGSARQQWELQGDRRCQSQLERANLRPCEQHLMQKIQRDEDSYERDPYSPSQDPYSPSPYDRRGAGSSQHQERCCNELNEFENNQRCMCEALQQ
IMENQSDRLQGRQQEQQFKRELRNLPQQCGLRAPQRCDLDVESGG

**>cf2TIP60-Q-Arah2**
MGILPSPGMPALLSLVSLLSVLLMGCVAESQDPKNIKIMRLVTGEDIIGNISESQGLITIKKAFVIIPMQATPGKPVQLVLSPWQPYTDDKEIVIDDSKV
ITITSPKDDIIKSYESHTRVLETRQMEKILALEREIERLQRQHDEGLAQMNEGSREELRQRRDAEIYRLEKEASGGGGSARQQWELQGDRRCQSQLERAN
LRPCEQHLMQKIQRDEDSYERDPYSPSQDPYSPSPYDRRGAGSSQHQERCCNELNEFENNQRCMCEALQQIMENQSDRLQGRQQEQQFKRELRNLPQQCG
LRAPQRCDLDVESGG

**>cf2TIP60-Q-Arah6-Arah2**
MGILPSPGMPALLSLVSLLSVLLMGCVAESQDPKNIKIMRLVTGEDIIGNISESQGLITIKKAFVIIPMQATPGKPVQLVLSPWQPYTDDKEIVIDDSKV
ITITSPKDDIIKSYESHTRVLETRQMEKILALEREIERLQRQHDEGLAQMNEGSREELRQRRDAEIYRLEKEASGGGGSAMRRERGRQGDSSSCERQVDG
VNLKPCEQHIMQRIMGEQEQYDSYNFGSTRSSDQQQRCCDELNEMENTQRCMCEALQQIMENQCDGLQDRQMVQHFKRELMNLPQQCNFGAPQRCDLDVS
GGRCGGGGSARQQWELQGDRRCQSQLERANLRPCEQHLMQKIQRDEDSYERDPYSPSQDPYSPSPYDRRGAGSSQHQERCCNELNEFENNQRCMCEALQQ
IMENQSDRLQGRQQEQQFKRELRNLPQQCGLRAPQRCDLDVESGG

**>cf2TIP60-Q-Ext-Arah2**
MGILPSPGMPALLSLVSLLSVLLMGCVAESQDPKNIKIMRLVTGEDIIGNISESQGLITIKKAFVIIPMQATPGKPVQLVLSPWQPYTDDKEIVIDDSKV
ITITSPKDDIIKSYESHTRVLEAAKAAAETRQMEKILALEREIERLQRQHDEGLAQMNEGSREELRQRRDAEIYRLEKEASGGGGSARQQWELQGDRRCQ
SQLERANLRPCEQHLMQKIQRDEDSYERDPYSPSQDPYSPSPYDRRGAGSSQHQERCCNELNEFENNQRCMCEALQQIMENQSDRLQGRQQEQQFKRELR
NLPQQCGLRAPQRCDLDVESGG

**>cf2TIP60-Q-Ext-Arah6-Arah2**
MGILPSPGMPALLSLVSLLSVLLMGCVAESQDPKNIKIMRLVTGEDIIGNISESQGLITIKKAFVIIPMQATPGKPVQLVLSPWQPYTDDKEIVIDDSKV
ITITSPKDDIIKSYESHTRVLEAAKAAAETRQMEKILALEREIERLQRQHDEGLAQMNEGSREELRQRRDAEIYRLEKEASGGGGSAMRRERGRQGDSSS
CERQVDGVNLKPCEQHIMQRIMGEQEQYDSYNFGSTRSSDQQQRCCDELNEMENTQRCMCEALQQIMENQCDGLQDRQMVQHFKRELMNLPQQCNFGAPQ
RCDLDVSGGRCGGGGSARQQWELQGDRRCQSQLERANLRPCEQHLMQKIQRDEDSYERDPYSPSQDPYSPSPYDRRGAGSSQHQERCCNELNEFENNQRC
MCEALQQIMENQSDRLQGRQQEQQFKRELRNLPQQCGLRAPQRCDLDVESGG

: Leader sequence
: Arah2; : Arah6
Sm cfMyoX: cf1/2TIP60; Sm Ext cfMyoX: cfTIP60-Ext
AS: Enzymatic site; : Linker
**L/A**: cysteine mutation; Q: L117Q mutation
**HHHHHH**: $His_6$ tag

b

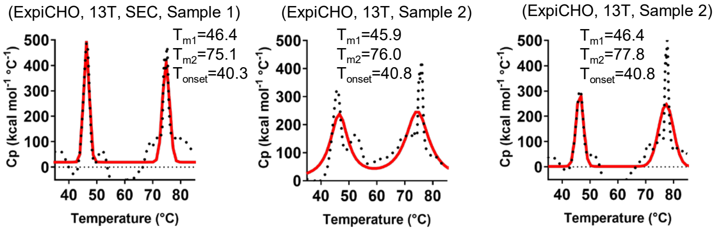


c

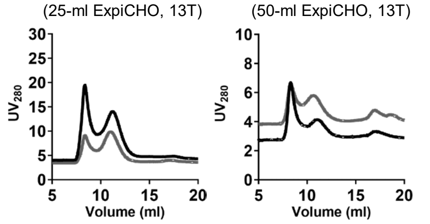

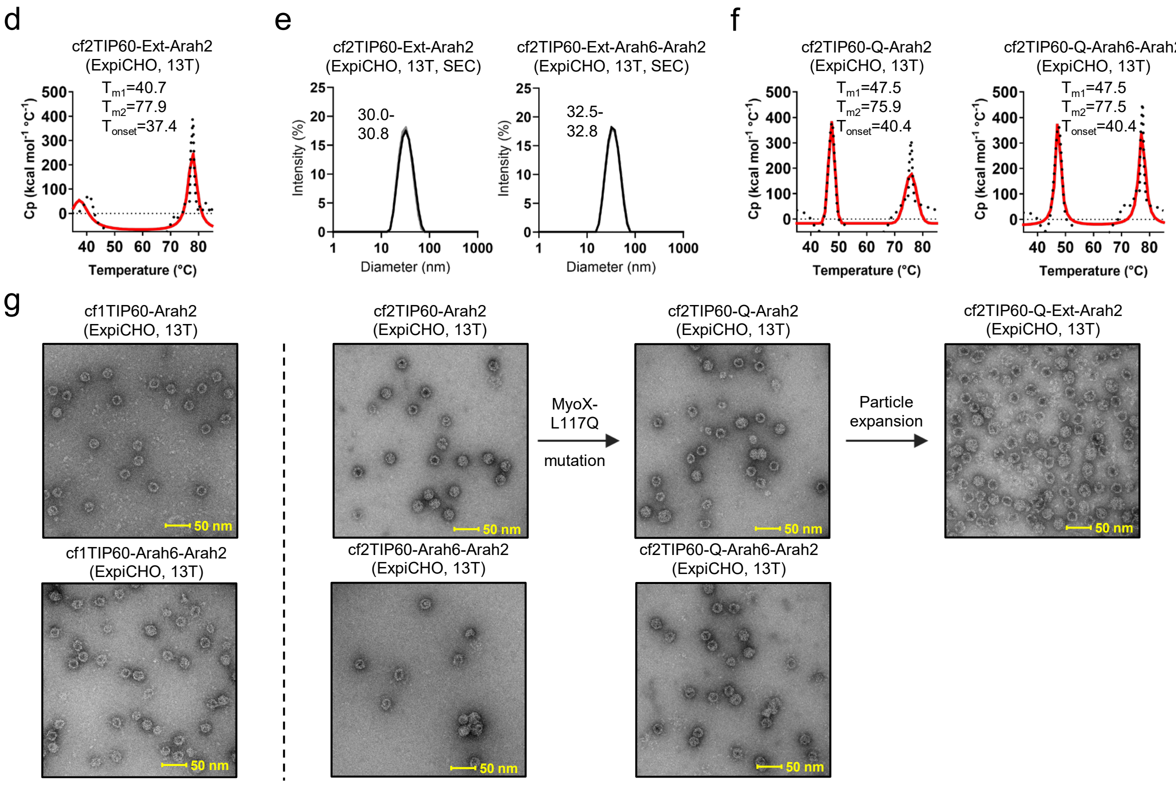


**Fig. S2. In vitro characterization of catfish TIP60 (cf1/2TIP60) constructs displaying peanut allergens**. (**a**) Amino acid sequences of cf1TIP60 and cf2TIP60 nanoparticles displaying Arah2 and tandem Arah6-Arah2, including cf2TIP60 variants with the L117Q mutation (Q) and L117Q plus expansion (Ext). Signal peptide, Arah2, Arah6, Sm-like domain (within the TIP60 subunit), catfish-derived MyoX domain (within the TIP60 subunit), 7-residue expansion segment (Ext), enzymatic site, and linkers are highlighted in yellow, brown, green, dark moss green, olive green, cyan, magenta, and gray, respectively. Cysteine residues in cf1-MyoX and cf2-MyoX are mutated to L and A, respectively (red), and the L117Q mutation in cf2-MyoX is shown in hot pink. (**b**) DSC thermograms of ExpiCHO-expressed cf2TIP60 constructs: cf2TIP60-Arah2 purified by immunoaffinity chromatography (IAC; 13T antibody column) followed by SEC (left), cf2TIP60-Arah2 from an independent production run after IAC (middle), and cf2TIP60-Arah6-Arah2 from an independent production run after IAC (right). (**c**) SEC profiles of ExpiCHO-expressed, IAC-purified cf2TIP60-Ext-Arah2 and cf2TIP60-Ext-Arah6-Arah2. (**d**) DSC thermogram of ExpiCHO-expressed, IAC-purified cf2TIP60-Ext-Arah2. (**e**) DLS profiles of ExpiCHO-expressed, IAC/SEC-purified cf2TIP60-Ext-Arah2 and cf2TIP60-Ext-Arah6-Arah2. (**f**) DSC thermograms of ExpiCHO-expressed, IAC-purified cf2TIP60-Q-Arah2 and cf2TIP60-Q-Arah6-Arah2 from an independent production run. In (**b**,**d**,**f**), experimental data (black points) and Gaussian fits (red lines) are shown, with melting temperatures ($T_{m1}$/$T_{m2}$) and onset temperature ($T_{onset}$) indicated. (**g**) Negative-stain EM micrographs of cf1TIP60 and cf2TIP60 constructs, separated by a vertical dotted line. Left, cf1TIP60-Arah2 and cf1TIP60-Arah6-Arah2. Right, cf2TIP60-Arah2/cf2TIP60-Arah6-Arah2 (column 1), cf2TIP60-Q-Arah2/cf2TIP60-Q-Arah6-Arah2 (column 2), and cf2TIP60-Q-Ext-Arah2 (column 3) from an independent production run. cf2TIP60-Q-Ext-Arah6-Arah2 was not analyzed due to low expression. Scale bar, 50 nm.

**Fig. S3**

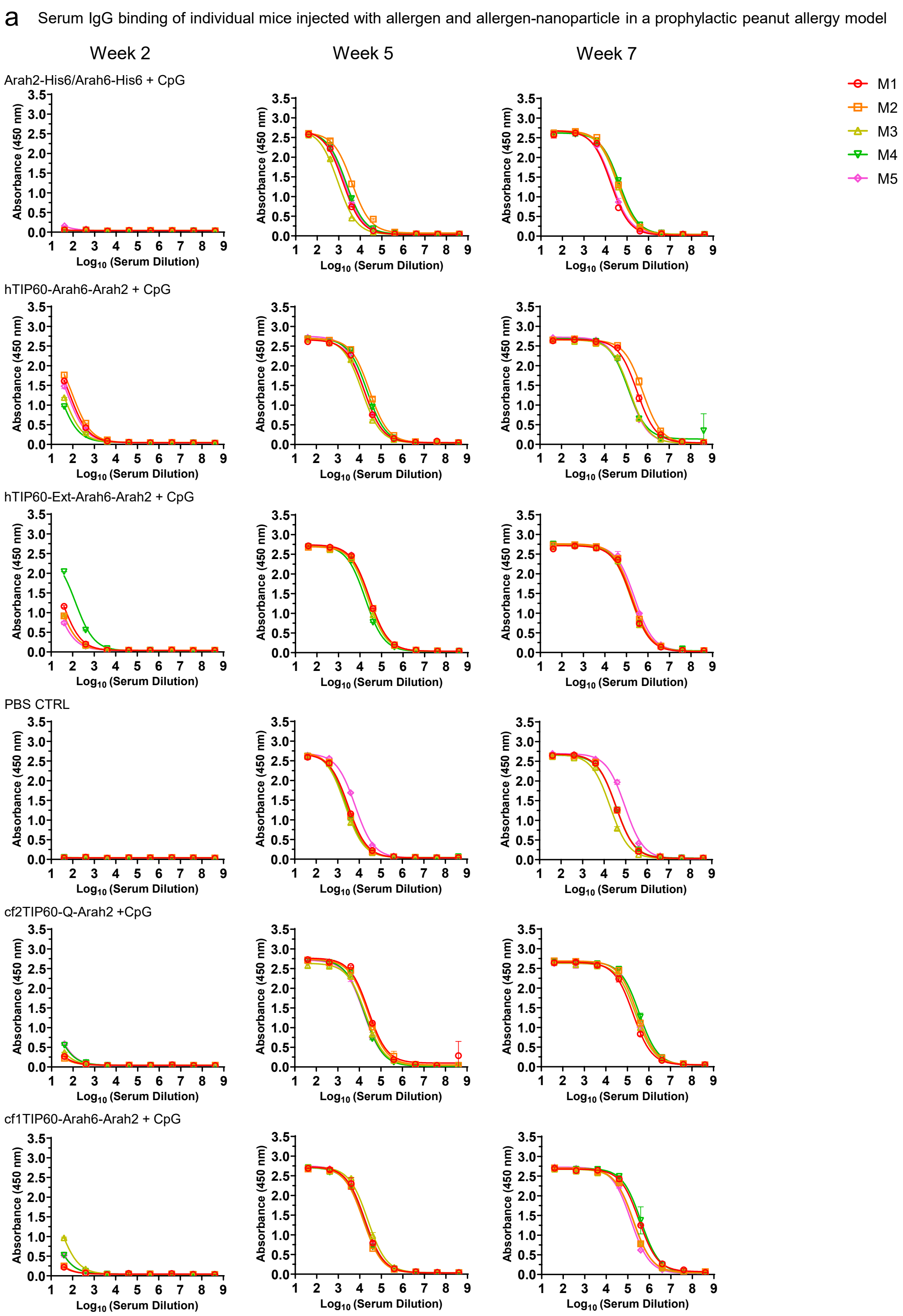
a
Serum IgG binding of individual mice injected with allergen and allergen-nanoparticle in a prophylactic peanut allergy model
Week 2
Week 5
Week 7
Arah2-His6/Arah6-His6 + CpG
hTIP60-Arah6-Arah2 + CpG
hTIP60-Ext-Arah6-Arah2 + CpG
PBS CTRL
cf2TIP60-Q-Arah2 +CpG
cf1TIP60-Arah6-Arah2 + CpG
Absorbance (450 nm)
Log10 (Serum Dilution)
M1
M2
M3
M4
M5

### b Mouse serum IgG ELISA $EC_{50}$ titers

| Antigen | $EC_{50}$ titers (week 2) | | | | | Geometric Mean |
|---|---|---|---|---|---|---|
| | M1 | M2 | M3 | M4 | M5 | |
| Arah2-$His_6$/Arah6-$His_6$ + CpG | 0.33 | 0.17 | 0.14 | 0.12 | 2.05 | 0.3 |
| hTIP60-Arah6-Arah2 + CpG | 71.49 | 95.58 | 35.77 | 24.22 | 57.43 | 50.8 |
| hTIP60-Ext-Arah6-Arah2 + CpG | 33.10 | 22.16 | 22.34 | 134.80 | 15.86 | 32.3 |
| PBS CTRL | 0.00 | 0.00 | 0.00 | 0.21 | 0.02 | 0.1 |

| Antigen | $EC_{50}$ titers (week 5) | | | | | Geometric Mean |
|---|---|---|---|---|---|---|
| | M1 | M2 | M3 | M4 | M5 | |
| Arah2-$His_6$/Arah6-$His_6$ + CpG | 1579 | 4068 | 921 | 2202 | 1855 | 1891 |
| hTIP60-Arah6-Arah2 + CpG | 17091 | 29388 | 13152 | 22425 | 16410 | 18930 |
| hTIP60-Ext-Arah6-Arah2 + CpG | 27877 | 28090 | 22946 | 17578 | 27170 | 24362 |
| PBS CTRL | 3057 | 2748 | 2174 | 2440 | 6442 | 3102 |

| Antigen | $EC_{50}$ titers (week 7) | | | | | Geometric Mean |
|---|---|---|---|---|---|---|
| | M1 | M2 | M3 | M4 | M5 | |
| Arah2-$His_6$/Arah6-$His_6$ + CpG | 17678 | 37813 | 34483 | 44977 | 19737 | 28986 |
| hTIP60-Arah6-Arah2 + CpG | 320312 | 589860 | 152029 | 124939 | 132835 | 216597 |
| hTIP60-Ext-Arah6-Arah2 + CpG | 174276 | 190926 | 161499 | 163303 | 240259 | 183988 |
| PBS CTRL | 34896 | 35489 | 18937 | 35766 | 92478 | 37839 |

| Antigen | $EC_{50}$ titers (week 2) | | | | | Geometric Mean |
|---|---|---|---|---|---|---|
| | M1 | M2 | M3 | M4 | M5 | |
| cf2TIP60-Q-Arah2 + CpG | 4.00 | 3.14 | 6.19 | 10.53 | 11.52 | 6.2 |
| cf1TIP60-Arah6-Arah2 + CpG | 3.10 | 3.71 | 24.13 | 9.70 | 9.84 | 7.7 |

| Antigen | $EC_{50}$ titers (week 5) | | | | | Geometric Mean |
|---|---|---|---|---|---|---|
| | M1 | M2 | M3 | M4 | M5 | |
| cf2TIP60-Q-Arah2 + CpG | 26267 | 24600 | 19815 | 16967 | 16160 | 20374 |
| cf1TIP60-Arah6-Arah2 + CpG | 17274 | 14352 | 23317 | 14801 | 15616 | 16795 |

| Antigen | $EC_{50}$ titers (week 7) | | | | | Geometric Mean |
|---|---|---|---|---|---|---|
| | M1 | M2 | M3 | M4 | M5 | |
| cf2TIP60-Q-Arah2 + CpG | 182291 | 284584 | 245997 | 358697 | 244530 | 256917 |
| cf1TIP60-Arah6-Arah2 + CpG | 333957 | 175982 | 187633 | 405563 | 140466 | 228886 |

### Statistical analysis

| One-way ANOVA with Tukey's multiple comparisons test (w2) | Summary | Adjusted P Value |
|---|---|---|
| Arah2-His6/Arah6-His6 + CpG vs. hTIP60-Arah6-Arah2 + CpG | * | 0.0322 |
| Arah2-His6/Arah6-His6 + CpG vs. hTIP60-Ext-Arah6-Arah2 + CpG | ns | 0.1032 |
| Arah2-His6/Arah6-His6 + CpG vs. PBS CTRL | ns | >0.9999 |
| hTIP60-Arah6-Arah2 + CpG vs. hTIP60-Ext-Arah6-Arah2 + CpG | ns | 0.9253 |
| hTIP60-Arah6-Arah2 + CpG vs. PBS CTRL | * | 0.0307 |
| hTIP60-Ext-Arah6-Arah2 + CpG vs. PBS CTRL | ns | 0.0988 |

| One-way ANOVA with Tukey's multiple comparisons test (w5) | Summary | Adjusted P Value |
|---|---|---|
| Arah2-His6/Arah6-His6 + CpG vs. hTIP60-Arah6-Arah2 + CpG | **** | <0.0001 |
| Arah2-His6/Arah6-His6 + CpG vs. hTIP60-Ext-Arah6-Arah2 + CpG | **** | <0.0001 |
| Arah2-His6/Arah6-His6 + CpG vs. PBS CTRL | ns | 0.9606 |
| hTIP60-Arah6-Arah2 + CpG vs. hTIP60-Ext-Arah6-Arah2 + CpG | ns | 0.2389 |
| hTIP60-Arah6-Arah2 + CpG vs. PBS CTRL | **** | <0.0001 |
| hTIP60-Ext-Arah6-Arah2 + CpG vs. PBS CTRL | **** | <0.0001 |

| One-way ANOVA with Tukey's multiple comparisons test (w7) | Summary | Adjusted P Value |
|---|---|---|
| Arah2-His6/Arah6-His6 + CpG vs. hTIP60-Arah6-Arah2 + CpG | * | 0.0112 |
| Arah2-His6/Arah6-His6 + CpG vs. hTIP60-Ext-Arah6-Arah2 + CpG | ns | 0.1162 |
| Arah2-His6/Arah6-His6 + CpG vs. PBS CTRL | ns | 0.9973 |
| hTIP60-Arah6-Arah2 + CpG vs. hTIP60-Ext-Arah6-Arah2 + CpG | ns | 0.6306 |
| hTIP60-Arah6-Arah2 + CpG vs. PBS CTRL | * | 0.0167 |
| hTIP60-Ext-Arah6-Arah2 + CpG vs. PBS CTRL | ns | 0.1627 |

| Welch's t test (w2) | Statistics | P Value |
|---|---|---|
| cf2TIP60-Q-Arah2 +CpG vs. cf1TIP60-Arah6-Arah2 + CpG | ns | 0.4961 |

| Welch's t test (w5) | Statistics | P Value |
|---|---|---|
| cf2TIP60-Q-Arah2 +CpG vs. cf1TIP60-Arah6-Arah2 + CpG | ns | 0.1952 |

| Welch's t test (w7) | Statistics | P Value |
|---|---|---|
| cf2TIP60-Q-Arah2 +CpG vs. cf1TIP60-Arah6-Arah2 + CpG | ns | 0.8133 |

Fig. S3

C Serum IgG binding of individual mice injected with allergen and allergen-nanoparticle in a therapeutic peanut allergy model

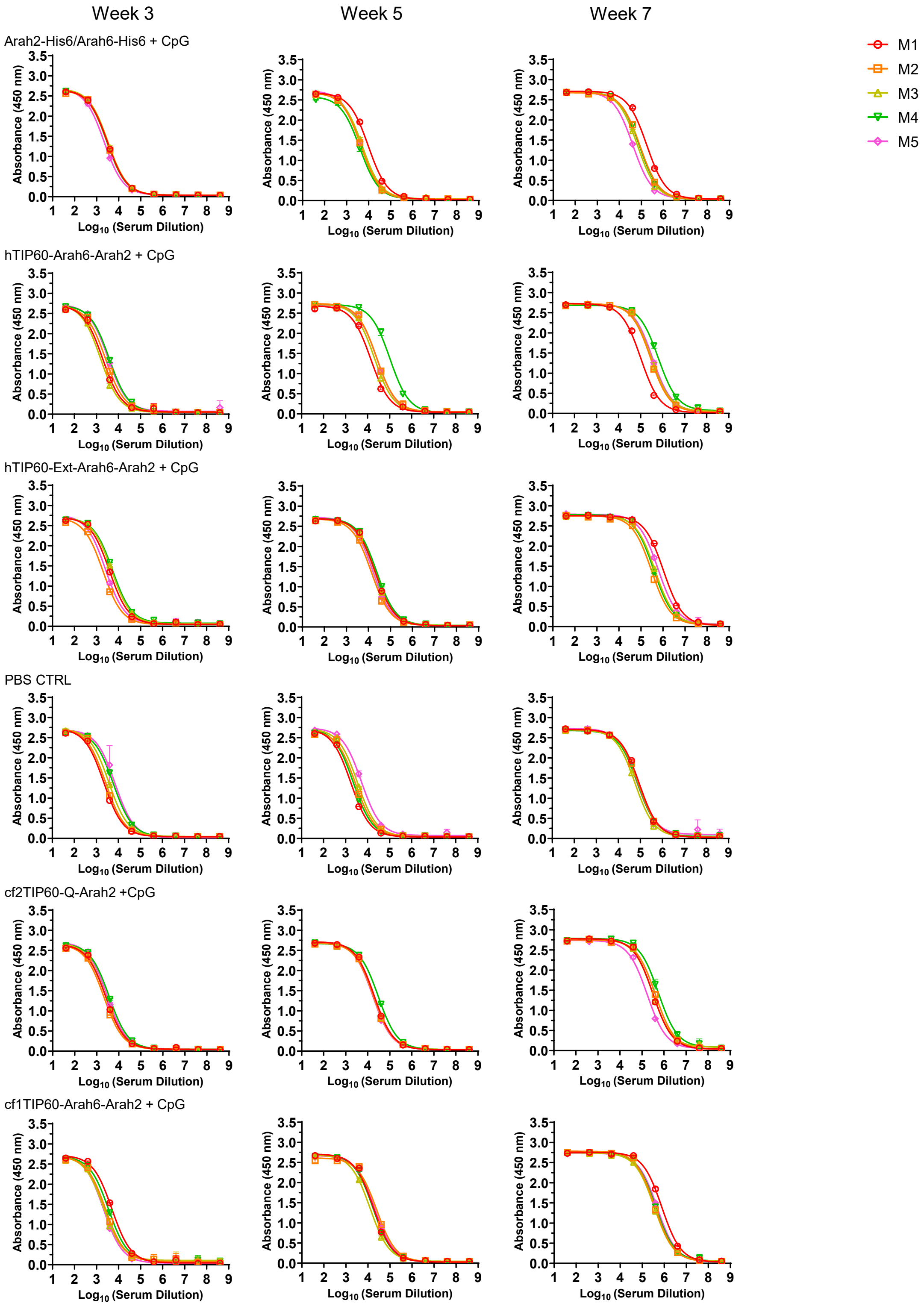

**Fig. S3**

d Mouse serum IgG ELISA $EC_{50}$ titers

| Antigen | $EC_{50}$ titers (week 3) | | | | | Geometric Mean |
|---|---|---|---|---|---|---|
| | M1 | M2 | M3 | M4 | M5 | |
| Arah2-$His_6$/Arah6-$His_6$ + CpG | 3130 | 3095 | 3265 | 2944 | 2181 | 2894 |
| hTIP60-Arah6-Arah2 + CpG | 1909 | 2576 | 1519 | 3825 | 3355 | 2491 |
| hTIP60-Ext-Arah6-Arah2 + CpG | 3860 | 1963 | 4857 | 5368 | 2547 | 3470 |
| PBS CTRL | 2192 | 2631 | 3719 | 5909 | 7115 | 3900 |

| Antigen | $EC_{50}$ titers (week 5) | | | | | Geometric Mean |
|---|---|---|---|---|---|---|
| | M1 | M2 | M3 | M4 | M5 | |
| Arah2-$His_6$/Arah6-$His_6$ + CpG | 9707 | 4435 | 5107 | 3955 | 4525 | 5236 |
| hTIP60-Arah6-Arah2 + CpG | 13872 | 25736 | 20412 | 103368 | 26634 | 28872 |
| hTIP60-Ext-Arah6-Arah2 + CpG | 20656 | 13641 | 19600 | 23894 | 16183 | 18446 |
| PBS CTRL | 1738 | 2825 | 3486 | 2246 | 5256 | 2891 |

| Antigen | $EC_{50}$ titers (week 7) | | | | | Geometric Mean |
|---|---|---|---|---|---|---|
| | M1 | M2 | M3 | M4 | M5 | |
| Arah2-$His_6$/Arah6-$His_6$ + CpG | 177665 | 87894 | 67330 | 82452 | 41520 | 81517 |
| hTIP60-Arah6-Arah2 + CpG | 101572 | 282317 | 306159 | 634550 | 345254 | 286294 |
| hTIP60-Ext-Arah6-Arah2 + CpG | 1050624 | 298827 | 434035 | 378204 | 604802 | 499742 |
| PBS CTRL | 88707 | 81734 | 57656 | 75822 | 67491 | 73460 |

| Antigen | $EC_{50}$ titers (week 3) | | | | | Geometric Mean |
|---|---|---|---|---|---|---|
| | M1 | M2 | M3 | M4 | M5 | |
| cf2TIP60-Q-Arah2 + CpG | 2553 | 2095 | 2556 | 3633 | 2982 | 2717 |
| cf1TIP60-Arah6-Arah2 + CpG | 4989 | 2527 | 2127 | 3491 | 2049 | 2861 |

| Antigen | $EC_{50}$ titers (week 5) | | | | | Geometric Mean |
|---|---|---|---|---|---|---|
| | M1 | M2 | M3 | M4 | M5 | |
| cf2TIP60-Q-Arah2 + CpG | 19618 | 18065 | 18102 | 30067 | 16836 | 20059 |
| cf1TIP60-Arah6-Arah2 + CpG | 17993 | 24937 | 12351 | 16908 | 20296 | 18023 |

| Antigen | $EC_{50}$ titers (week 7) | | | | | Geometric Mean |
|---|---|---|---|---|---|---|
| | M1 | M2 | M3 | M4 | M5 | |
| cf2TIP60-Q-Arah2 + CpG | 317186 | 406012 | 304137 | 555971 | 172785 | 327413 |
| cf1TIP60-Arah6-Arah2 + CpG | 776851 | 370944 | 350161 | 404338 | 475249 | 454481 |

Statistical analysis

| One-way ANOVA with Tukey's multiple comparisons test (w3) | Summary | Adjusted P Value |
|---|---|---|
| Arah2-His6/Arah6-His6 + CpG vs. hTIP60-Arah6-Arah2 + CpG | ns | 0.9877 |
| Arah2-His6/Arah6-His6 + CpG vs. hTIP60-Ext-Arah6-Arah2 + CpG | ns | 0.803 |
| Arah2-His6/Arah6-His6 + CpG vs. PBS CTRL | ns | 0.4175 |
| hTIP60-Arah6-Arah2 + CpG vs. hTIP60-Ext-Arah6-Arah2 + CpG | ns | 0.6184 |
| hTIP60-Arah6-Arah2 + CpG vs. PBS CTRL | ns | 0.2654 |
| hTIP60-Ext-Arah6-Arah2 + CpG vs. PBS CTRL | ns | 0.9051 |

| One-way ANOVA with Tukey's multiple comparisons test (w5) | Summary | Adjusted P Value |
|---|---|---|
| Arah2-His6/Arah6-His6 + CpG vs. hTIP60-Arah6-Arah2 + CpG | ns | 0.0608 |
| Arah2-His6/Arah6-His6 + CpG vs. hTIP60-Ext-Arah6-Arah2 + CpG | ns | 0.6794 |
| Arah2-His6/Arah6-His6 + CpG vs. PBS CTRL | ns | 0.9967 |
| hTIP60-Arah6-Arah2 + CpG vs. hTIP60-Ext-Arah6-Arah2 + CpG | ns | 0.3893 |
| hTIP60-Arah6-Arah2 + CpG vs. PBS CTRL | * | 0.0409 |
| hTIP60-Ext-Arah6-Arah2 + CpG vs. PBS CTRL | ns | 0.5564 |

| One-way ANOVA with Tukey's multiple comparisons test (w7) | Summary | Adjusted P Value |
|---|---|---|
| Arah2-His6/Arah6-His6 + CpG vs. hTIP60-Arah6-Arah2 + CpG | ns | 0.1857 |
| Arah2-His6/Arah6-His6 + CpG vs. hTIP60-Ext-Arah6-Arah2 + CpG | ** | 0.0046 |
| Arah2-His6/Arah6-His6 + CpG vs. PBS CTRL | ns | 0.9987 |
| hTIP60-Arah6-Arah2 + CpG vs. hTIP60-Ext-Arah6-Arah2 + CpG | ns | 0.2566 |
| hTIP60-Arah6-Arah2 + CpG vs. PBS CTRL | ns | 0.1444 |
| hTIP60-Ext-Arah6-Arah2 + CpG vs. PBS CTRL | ** | 0.0034 |

| Welch's t test (w3) | Statistics | P Value |
|---|---|---|
| cf2TIP60-Q-Arah2 +CpG vs. cf1TIP60-Arah6-Arah2 + CpG | ns | 0.6708 |

| Welch's t test (w5) | Statistics | P Value |
|---|---|---|
| cf2TIP60-Q-Arah2 +CpG vs. cf1TIP60-Arah6-Arah2 + CpG | ns | 0.5399 |

| Welch's t test (w7) | Statistics | P Value |
|---|---|---|
| cf2TIP60-Q-Arah2 +CpG vs. cf1TIP60-Arah6-Arah2 + CpG | ns | 0.2534 |

Fig. S3

e Mouse serum IgE ELISA $OD_{450}$ values in a prophylactic peanut allergy model

| Antigen | $OD_{450}$ values (week 2) | | | | | Geometric Mean |
|---|---|---|---|---|---|---|
| | M1 | M2 | M3 | M4 | M5 | |
| Arah2-$His_6$/Arah6-$His_6$ + CpG | 0.0579 | 0.05755 | 0.0607 | 0.05255 | 0.0573 | 0.0571 |
| hTIP60-Arah6-Arah2 + CpG | 0.0581 | 0.05815 | 0.0576 | 0.05825 | 0.0586 | 0.0581 |
| hTIP60-Ext-Arah6-Arah2 + CpG | 0.04525 | 0.0458 | 0.04695 | 0.04565 | 0.0385 | 0.0443 |
| PBS CTRL | 0.04765 | 0.0477 | 0.04635 | 0.041 | 0.04675 | 0.0458 |

| Antigen | $OD_{450}$ values (week 5) | | | | | Geometric Mean |
|---|---|---|---|---|---|---|
| | M1 | M2 | M3 | M4 | M5 | |
| Arah2-$His_6$/Arah6-$His_6$ + CpG | 0.09795 | 0.14875 | 0.0984 | 0.1516 | 0.20835 | 0.1353 |
| hTIP60-Arah6-Arah2 + CpG | 0.0967 | 0.0731 | 0.0896 | 0.0595 | 0.0665 | 0.0758 |
| hTIP60-Ext-Arah6-Arah2 + CpG | 0.086 | 0.06665 | 0.07165 | 0.07295 | 0.11245 | 0.0804 |
| PBS CTRL | 0.0695 | 0.13745 | 0.07065 | 0.08755 | 0.07665 | 0.0853 |

| Antigen | $OD_{450}$ values (week 7) | | | | | Geometric Mean |
|---|---|---|---|---|---|---|
| | M1 | M2 | M3 | M4 | M5 | |
| Arah2-$His_6$/Arah6-$His_6$ + CpG | 0.05615 | 0.07685 | 0.06105 | 0.08455 | 0.15375 | 0.0807 |
| hTIP60-Arah6-Arah2 + CpG | 0.07 | 0.0564 | 0.07805 | 0.06715 | 0.0491 | 0.0633 |
| hTIP60-Ext-Arah6-Arah2 + CpG | 0.0619 | 0.0628 | 0.0641 | 0.0776 | 0.07275 | 0.0676 |
| PBS CTRL | 0.0513 | 0.051 | 0.06335 | 0.0582 | 0.04195 | 0.0527 |

| Antigen | $OD_{450}$ values (week 2) | | | | | Geometric Mean |
|---|---|---|---|---|---|---|
| | M1 | M2 | M3 | M4 | M5 | |
| cf2TIP60-Q-Arah2 + CpG | 0.0565 | 0.0553 | 0.0553 | 0.0558 | 0.05205 | 0.0550 |
| cf1TIP60-Arah6-Arah2 + CpG | 0.0523 | 0.0538 | 0.0595 | 0.05555 | 0.0573 | 0.0556 |

| Antigen | $OD_{450}$ values (week 5) | | | | | Geometric Mean |
|---|---|---|---|---|---|---|
| | M1 | M2 | M3 | M4 | M5 | |
| cf2TIP60-Q-Arah2 + CpG | 0.04845 | 0.0545 | 0.0751 | 0.11505 | 0.1268 | 0.0780 |
| cf1TIP60-Arah6-Arah2 + CpG | 0.0692 | 0.0792 | 0.07705 | 0.08125 | 0.0931 | 0.0796 |

| Antigen | $OD_{450}$ values (week 7) | | | | | Geometric Mean |
|---|---|---|---|---|---|---|
| | M1 | M2 | M3 | M4 | M5 | |
| cf2TIP60-Q-Arah2 + CpG | 0.0822 | 0.0677 | 0.0754 | 0.07275 | 0.08695 | 0.0767 |
| cf1TIP60-Arah6-Arah2 + CpG | 0.0558 | 0.0677 | 0.0598 | 0.06885 | 0.061 | 0.0624 |

Statistical analysis

| One-way ANOVA with Tukey's multiple comparisons test (w2) | Summary | Adjusted P Value |
|---|---|---|
| Arah2-His6/Arah6-His6 + CpG vs. hTIP60-Arah6-Arah2 + CpG | ns | 0.9418 |
| Arah2-His6/Arah6-His6 + CpG vs. hTIP60-Ext-Arah6-Arah2 + CpG | **** | <0.0001 |
| Arah2-His6/Arah6-His6 + CpG vs. PBS CTRL | **** | <0.0001 |
| hTIP60-Arah6-Arah2 + CpG vs. hTIP60-Ext-Arah6-Arah2 + CpG | **** | <0.0001 |
| hTIP60-Arah6-Arah2 + CpG vs. PBS CTRL | **** | <0.0001 |
| hTIP60-Ext-Arah6-Arah2 + CpG vs. PBS CTRL | ns | 0.8185 |

| One-way ANOVA with Tukey's multiple comparisons test (w5) | Summary | Adjusted P Value |
|---|---|---|
| Arah2-His6/Arah6-His6 + CpG vs. hTIP60-Arah6-Arah2 + CpG | * | 0.0165 |
| Arah2-His6/Arah6-His6 + CpG vs. hTIP60-Ext-Arah6-Arah2 + CpG | * | 0.0277 |
| Arah2-His6/Arah6-His6 + CpG vs. PBS CTRL | ns | 0.054 |
| hTIP60-Arah6-Arah2 + CpG vs. hTIP60-Ext-Arah6-Arah2 + CpG | ns | 0.9936 |
| hTIP60-Arah6-Arah2 + CpG vs. PBS CTRL | ns | 0.9292 |
| hTIP60-Ext-Arah6-Arah2 + CpG vs. PBS CTRL | ns | 0.9855 |

| One-way ANOVA with Tukey's multiple comparisons test (w7) | Summary | Adjusted P Value |
|---|---|---|
| Arah2-His6/Arah6-His6 + CpG vs. hTIP60-Arah6-Arah2 + CpG | ns | 0.3714 |
| Arah2-His6/Arah6-His6 + CpG vs. hTIP60-Ext-Arah6-Arah2 + CpG | ns | 0.5216 |
| Arah2-His6/Arah6-His6 + CpG vs. PBS CTRL | ns | 0.1001 |
| hTIP60-Arah6-Arah2 + CpG vs. hTIP60-Ext-Arah6-Arah2 + CpG | ns | 0.9924 |
| hTIP60-Arah6-Arah2 + CpG vs. PBS CTRL | ns | 0.8441 |
| hTIP60-Ext-Arah6-Arah2 + CpG vs. PBS CTRL | ns | 0.697 |

| Welch's t test (w2) | Statistics | P Value |
|---|---|---|
| cf2TIP60-Q-Arah2 +CpG vs. cf1TIP60-Arah6-Arah2 + CpG | ns | 0.6521 |

| Welch's t test (w5) | Statistics | P Value |
|---|---|---|
| cf2TIP60-Q-Arah2 +CpG vs. cf1TIP60-Arah6-Arah2 + CpG | ns | 0.816 |

| Welch's t test (w7) | Statistics | P Value |
|---|---|---|
| cf2TIP60-Q-Arah2 +CpG vs. cf1TIP60-Arah6-Arah2 + CpG | * | 0.0106 |

**f** Mouse serum IgE ELISA $OD_{450}$ values in a therapeutic peanut allergy model

| Antigen | $OD_{450}$ values (week 3) | | | | | Geometric Mean |
|---|---|---|---|---|---|---|
| | M1 | M2 | M3 | M4 | M5 | |
| Arah2-$His_6$/Arah6-$His_6$ + CpG | 0.12365 | 0.13965 | 0.1329 | 0.1451 | 0.1228 | 0.1325 |
| hTIP60-Arah6-Arah2 + CpG | 0.10445 | 0.1115 | 0.11555 | 0.11865 | 0.11255 | 0.1124 |
| hTIP60-Ext-Arah6-Arah2 + CpG | 0.0959 | 0.12295 | 0.11845 | 0.1336 | 0.1452 | 0.1221 |
| PBS CTRL | 0.1361 | 0.1234 | 0.1399 | 0.17545 | 0.1354 | 0.1410 |

| Antigen | $OD_{450}$ values (week 5) | | | | | Geometric Mean |
|---|---|---|---|---|---|---|
| | M1 | M2 | M3 | M4 | M5 | |
| Arah2-$His_6$/Arah6-$His_6$ + CpG | 0.30835 | 0.1268 | 0.33245 | 0.2009 | 0.34075 | 0.2454 |
| hTIP60-Arah6-Arah2 + CpG | 0.0875 | 0.13975 | 0.11035 | 0.0787 | 0.2286 | 0.1194 |
| hTIP60-Ext-Arah6-Arah2 + CpG | 0.082 | 0.1293 | 0.2494 | 0.20665 | 0.05595 | 0.1250 |
| PBS CTRL | 0.1756 | 0.27995 | 0.21975 | 0.32045 | 0.22855 | 0.2397 |

| Antigen | $OD_{450}$ values (week 7) | | | | | Geometric Mean |
|---|---|---|---|---|---|---|
| | M1 | M2 | M3 | M4 | M5 | |
| Arah2-$His_6$/Arah6-$His_6$ + CpG | 0.0836 | 0.07125 | 0.1018 | 0.0689 | 0.09875 | 0.0838 |
| hTIP60-Arah6-Arah2 + CpG | 0.08105 | 0.08335 | 0.0843 | 0.06115 | 0.0925 | 0.0797 |
| hTIP60-Ext-Arah6-Arah2 + CpG | 0.0579 | 0.08 | 0.1068 | 0.0846 | 0.09345 | 0.0829 |
| PBS CTRL | 0.0869 | 0.0789 | 0.0739 | 0.08735 | 0.0777 | 0.0808 |

| Antigen | $OD_{450}$ values (week 3) | | | | | Geometric Mean |
|---|---|---|---|---|---|---|
| | M1 | M2 | M3 | M4 | M5 | |
| cf2TIP60-Q-Arah2 + CpG | 0.14315 | 0.21475 | 0.112 | 0.11215 | 0.11715 | 0.1352 |
| cf1TIP60-Arah6-Arah2 + CpG | 0.1307 | 0.1309 | 0.1197 | 0.1473 | 0.1456 | 0.1344 |

| Antigen | $OD_{450}$ values (week 5) | | | | | Geometric Mean |
|---|---|---|---|---|---|---|
| | M1 | M2 | M3 | M4 | M5 | |
| cf2TIP60-Q-Arah2 + CpG | 0.09045 | 0.204 | 0.1522 | 0.1396 | 0.18175 | 0.1481 |
| cf1TIP60-Arah6-Arah2 + CpG | 0.1169 | 0.156 | 0.1658 | 0.19865 | 0.26725 | 0.1742 |

| Antigen | $OD_{450}$ values (week 7) | | | | | Geometric Mean |
|---|---|---|---|---|---|---|
| | M1 | M2 | M3 | M4 | M5 | |
| cf2TIP60-Q-Arah2 + CpG | 0.0729 | 0.078 | 0.0908 | 0.07545 | 0.1288 | 0.0871 |
| cf1TIP60-Arah6-Arah2 + CpG | 0.0746 | 0.09445 | 0.08285 | 0.1074 | 0.10075 | 0.0912 |

Statistical analysis

| One-way ANOVA with Tukey's multiple comparisons test (w3) | Summary | Adjusted P Value |
|---|---|---|
| Arah2-His6/Arah6-His6 + CpG vs. hTIP60-Arah6-Arah2 + CpG | ns | 0.1658 |
| Arah2-His6/Arah6-His6 + CpG vs. hTIP60-Ext-Arah6-Arah2 + CpG | ns | 0.7286 |
| Arah2-His6/Arah6-His6 + CpG vs. PBS CTRL | ns | 0.7511 |
| hTIP60-Arah6-Arah2 + CpG vs. hTIP60-Ext-Arah6-Arah2 + CpG | ns | 0.6605 |
| hTIP60-Arah6-Arah2 + CpG vs. PBS CTRL | * | 0.0258 |
| hTIP60-Ext-Arah6-Arah2 + CpG vs. PBS CTRL | ns | 0.2143 |

| One-way ANOVA with Tukey's multiple comparisons test (w5) | Summary | Adjusted P Value |
|---|---|---|
| Arah2-His6/Arah6-His6 + CpG vs. hTIP60-Arah6-Arah2 + CpG | ns | 0.0551 |
| Arah2-His6/Arah6-His6 + CpG vs. hTIP60-Ext-Arah6-Arah2 + CpG | ns | 0.102 |
| Arah2-His6/Arah6-His6 + CpG vs. PBS CTRL | ns | 0.9835 |
| hTIP60-Arah6-Arah2 + CpG vs. hTIP60-Ext-Arah6-Arah2 + CpG | ns | 0.987 |
| hTIP60-Arah6-Arah2 + CpG vs. PBS CTRL | ns | 0.1072 |
| hTIP60-Ext-Arah6-Arah2 + CpG vs. PBS CTRL | ns | 0.1892 |

| One-way ANOVA with Tukey's multiple comparisons test (w7) | Summary | Adjusted P Value |
|---|---|---|
| Arah2-His6/Arah6-His6 + CpG vs. hTIP60-Arah6-Arah2 + CpG | ns | 0.9543 |
| Arah2-His6/Arah6-His6 + CpG vs. hTIP60-Ext-Arah6-Arah2 + CpG | ns | >0.9999 |
| Arah2-His6/Arah6-His6 + CpG vs. PBS CTRL | ns | 0.967 |
| hTIP60-Arah6-Arah2 + CpG vs. hTIP60-Ext-Arah6-Arah2 + CpG | ns | 0.9628 |
| hTIP60-Arah6-Arah2 + CpG vs. PBS CTRL | ns | >0.9999 |
| hTIP60-Ext-Arah6-Arah2 + CpG vs. PBS CTRL | ns | 0.9739 |

| Welch's t test (w3) | Statistics | P Value |
|---|---|---|
| cf2TIP60-Q-Arah2 +CpG vs. cf1TIP60-Arah6-Arah2 + CpG | ns | 0.8158 |

| Welch's t test (w5) | Statistics | P Value |
|---|---|---|
| cf2TIP60-Q-Arah2 +CpG vs. cf1TIP60-Arah6-Arah2 + CpG | ns | 0.417 |

| Welch's t test (w7) | Statistics | P Value |
|---|---|---|
| cf2TIP60-Q-Arah2 +CpG vs. cf1TIP60-Arah6-Arah2 + CpG | ns | 0.8209 |

**Fig. S3. Antibody responses of adjuvanted allergen and allergen-nanoparticle in mice.** (**a**) ELISA IgG binding curves of mouse sera from allergen and allergen-nanoparticle groups (n = 5 mice/group) in a prophylactic peanut allergy model binding to the coating antigen crude peanut extract (CPE). (**b**) (Top) Summary of geometric mean $EC_{50}$ titers measured for allergen and allergen-nanoparticle groups against CPE. Color coding indicates $EC_{50}$ levels (green to red: low to high binding). (Bottom) Summary of statistical analysis performed for each timepoint. Note: $EC_{50}$ values at week 2 were derived by setting the minimum/maximum $OD_{450}$ values to 0.0/2.5. (**c**) ELISA IgG binding curves of mouse sera from allergen and allergen-nanoparticle groups (n = 5 mice/group) in a therapeutic peanut allergy model binding to the coating antigen CPE. (**d**) (Top) Summary of geometric mean $EC_{50}$ titers measured for allergen and allergen-nanoparticle groups against CPE. Color coding indicates $EC_{50}$ levels (green to red: low to high binding). (Bottom) Summary of statistical analysis performed for each timepoint. (**e**) Allergen-induced IgE responses of mouse sera from allergen and allergen-nanoparticle groups in a prophylactic peanut allergy model binding to CPE. Summary of absorbance at 450 nm ($A_{450}$) values from serum IgE induced by allergen and allergen-nanoparticle groups against CPE. Color coding indicates $A_{450}$ values (green to red: low to high binding). Summary of statistical analysis performed for each timepoint. (**f**) Summary of absorbance at $A_{450}$ values from serum IgE induced by allergen and allergen-nanoparticle groups in a therapeutic peanut allergy model against CPE. Summary of statistical analysis performed for each timepoint. Error bars represent the difference between duplicate measurements at each concentration for each sample. $EC_{50}$ values were calculated using GraphPad Prism version 10.6.1. Data were analyzed using one-way ANOVA, followed by Tukey's multiple comparison post hoc test for each timepoint. Two-tailed unpaired t-tests with Welch's correction were used when comparing two independent groups. For significance, ns (not significant), $*p < 0.05$, $**p < 0.01$, and $****p < 0.0001$.

Fig. S4

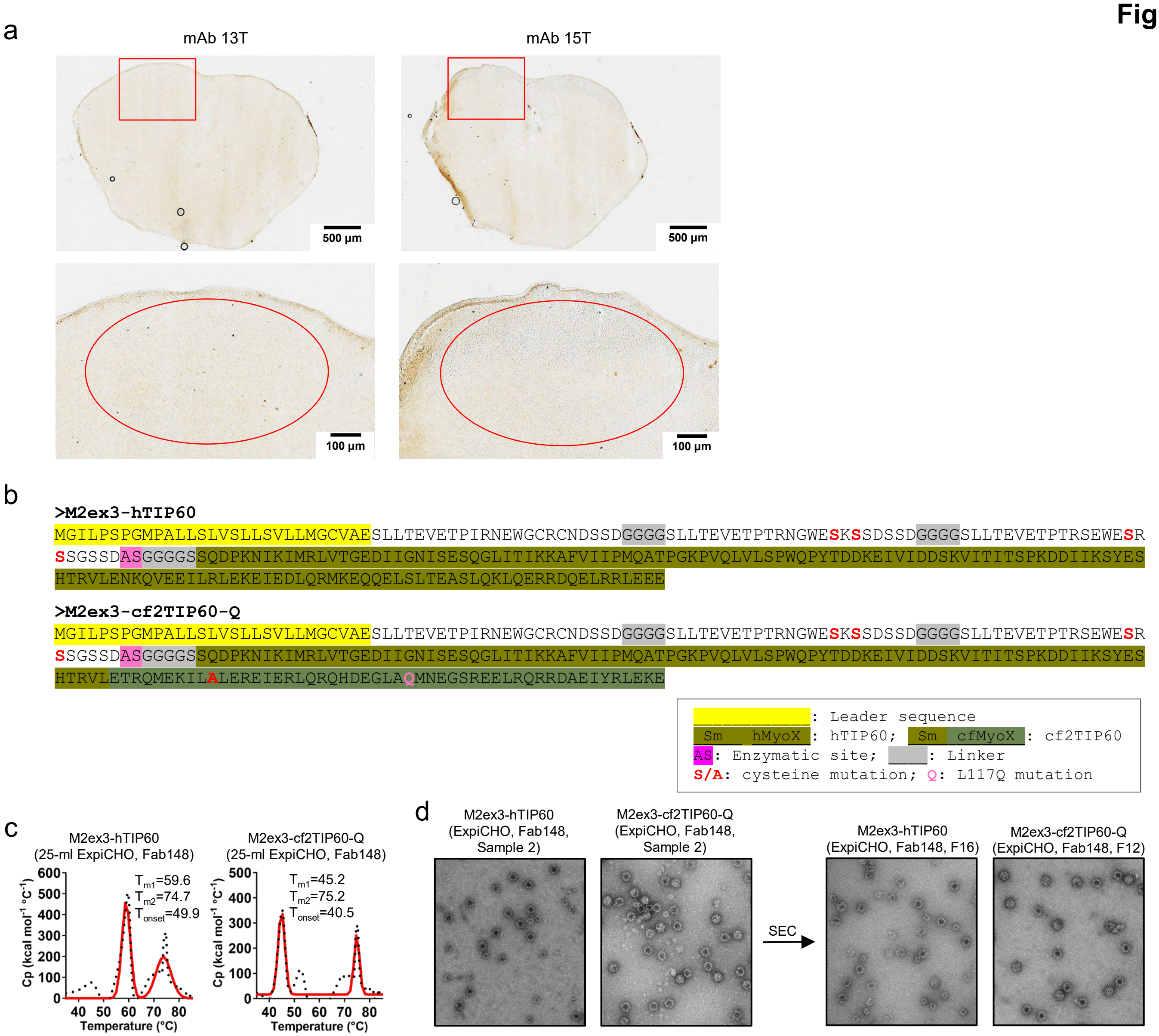


**Fig. S4. Immunohistological analysis of human and catfish TIP60 nanoparticles presenting a surrogate influenza M2ex3 antigen.** (**a**) Immunostaining images of lymph node tissues 48 hours after injection with cf2TIP60-Q-Arah6-Arah2, stained using antibodies 13T or 15T. No signals were observed in lymph node tissues. (**b**) Amino acid sequences of two M2ex3-displaying TIP60 constructs, M2ex3-hTIP60 and M2ex3-cf2TIP60-Q. A 5GS linker is inserted between the tandem M2ex3 antigen and the TIP60 nanoparticle backbone. Signal peptide, Sm-like domain (within the TIP60 subunit), catfish-derived MyoX domain (within the TIP60 subunit), enzymatic site, and linkers are highlighted in yellow, brown, green, dark moss green, olive green, magenta, and gray, respectively. Cysteine residues in the second and third M2e peptides and in cf2-MyoX are mutated to S and A, respectively (red), and the L117Q mutation in cf2-MyoX is shown in hot pink. (**c**) DSC thermograms of ExpiCHO-expressed M2ex3-hTIP60 and M2ex3-cf2TIP60-Q following immunoaffinity chromatography (IAC) purification on a Fab148 antibody column. Experimental data (black points) and Gaussian fits (red lines) are shown, with melting temperatures (Tm1/Tm2) and onset temperatures (Tonset) indicated. (**d**) Negative-stain EM micrographs of M2ex3-hTIP60 and M2ex3-cf2TIP60-Q before (left) and after (right) SEC purification. The fraction number is indicated for each SEC-purified sample. Scale bar, 50 nm.

**Fig. S5**

a cf2TIP60-Arah6-Arah2 (yellow arrow) aligned on FDC dendrites
at 12 hours

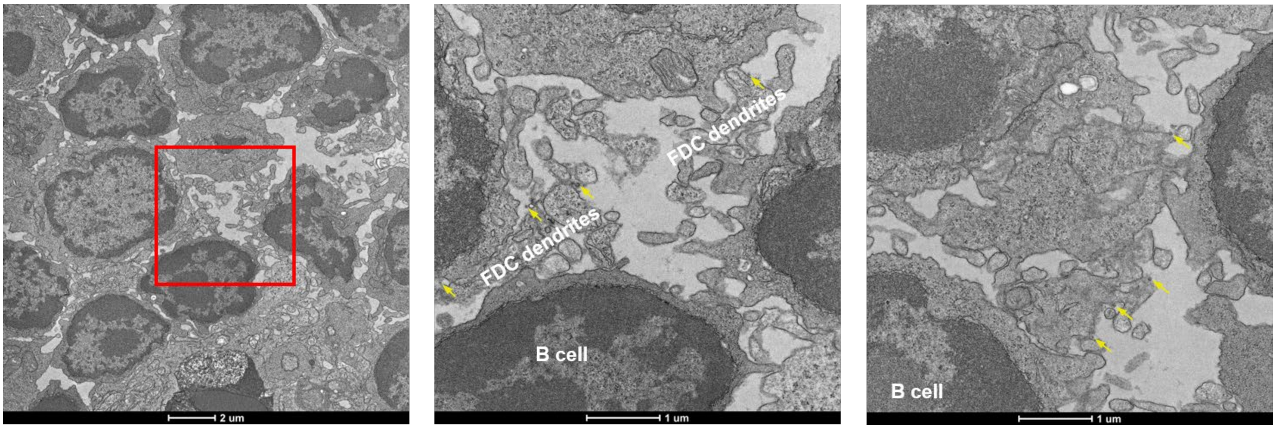


b cf2TIP60-Arah6-Arah2 (yellow arrow) aligned on FDC dendrites
at 48 hours

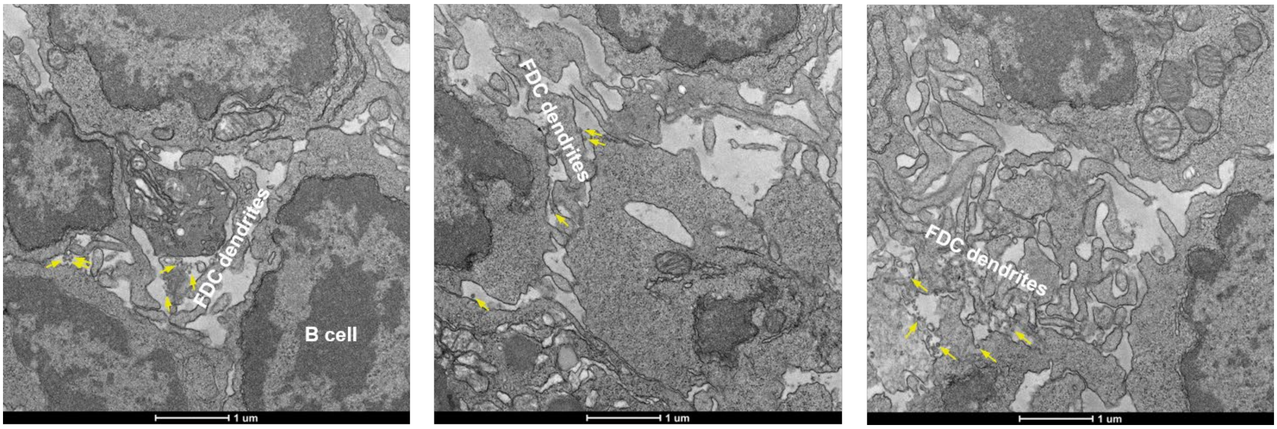


c cf2TIP60-Arah2 (yellow arrow) internalized by B cells
at 48 hours

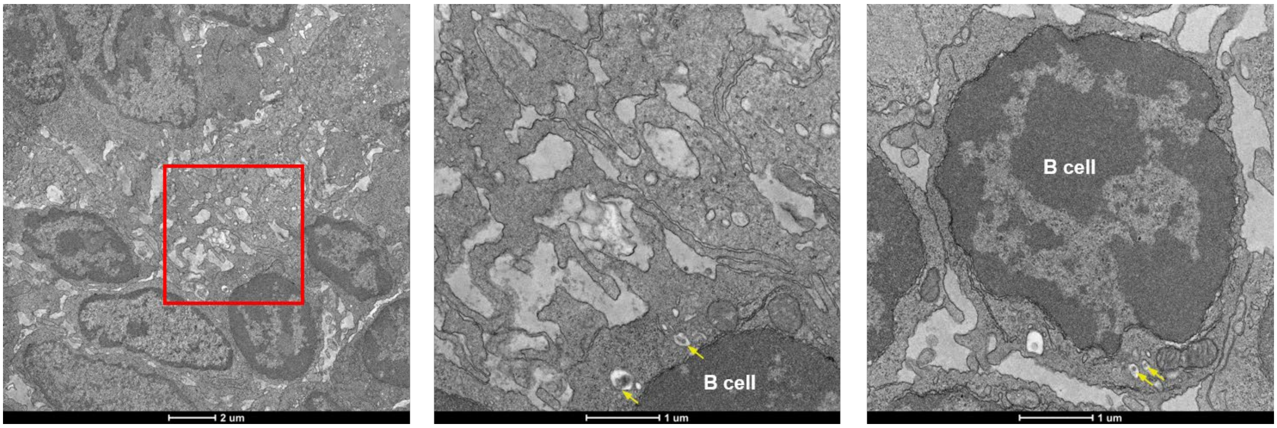

d cf2TIP60-Arah2 (yellow arrow) interact with macrophages at 12 hours

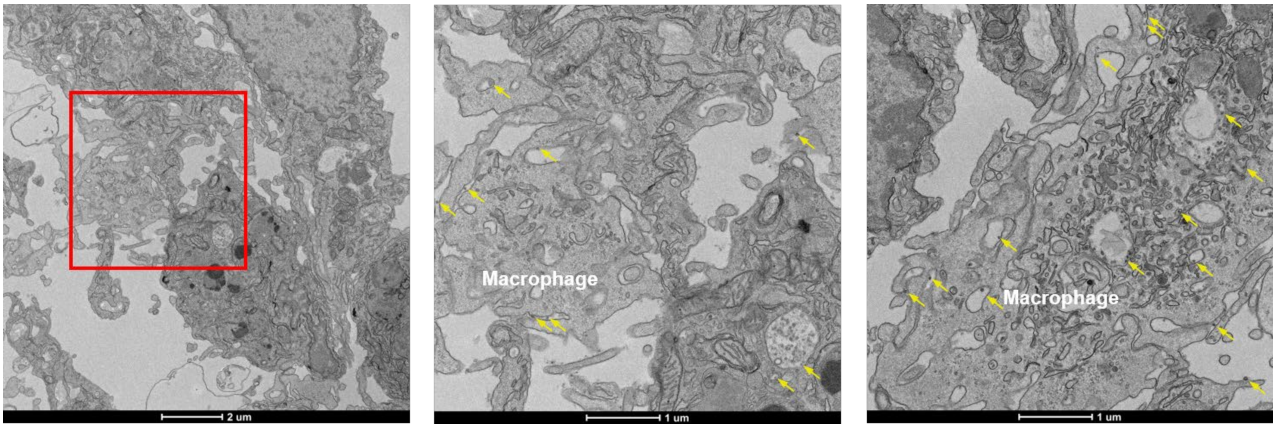


**Fig. S5. TEM images of cf2TIP60-Arah2 and cf2TIP60-Arah6-Arah2 interacting with FDCs, B cells and macrophages in a lymph nodes.** cf2TIP60-Arah6-Arah2 aligned on FDC dendrites at (**a**) 12 hours and (**b**) 48 hours after a single-dose injection (2 footpads, 50 µg/footpad). (**c**) cf2TIP60-Arah2 internalized by B cells at 48 hours post-injection. (**d**) cf2TIP60-Arah2 interact with macrophages at 12 hours post-injection. TEM images were performed on two popliteal sentinel lymph nodes for each construct. cf2TIP60-Arah2 and cf2TIP60-Arah6-Arah2 are indicated by yellow arrows.

**Fig. S6**

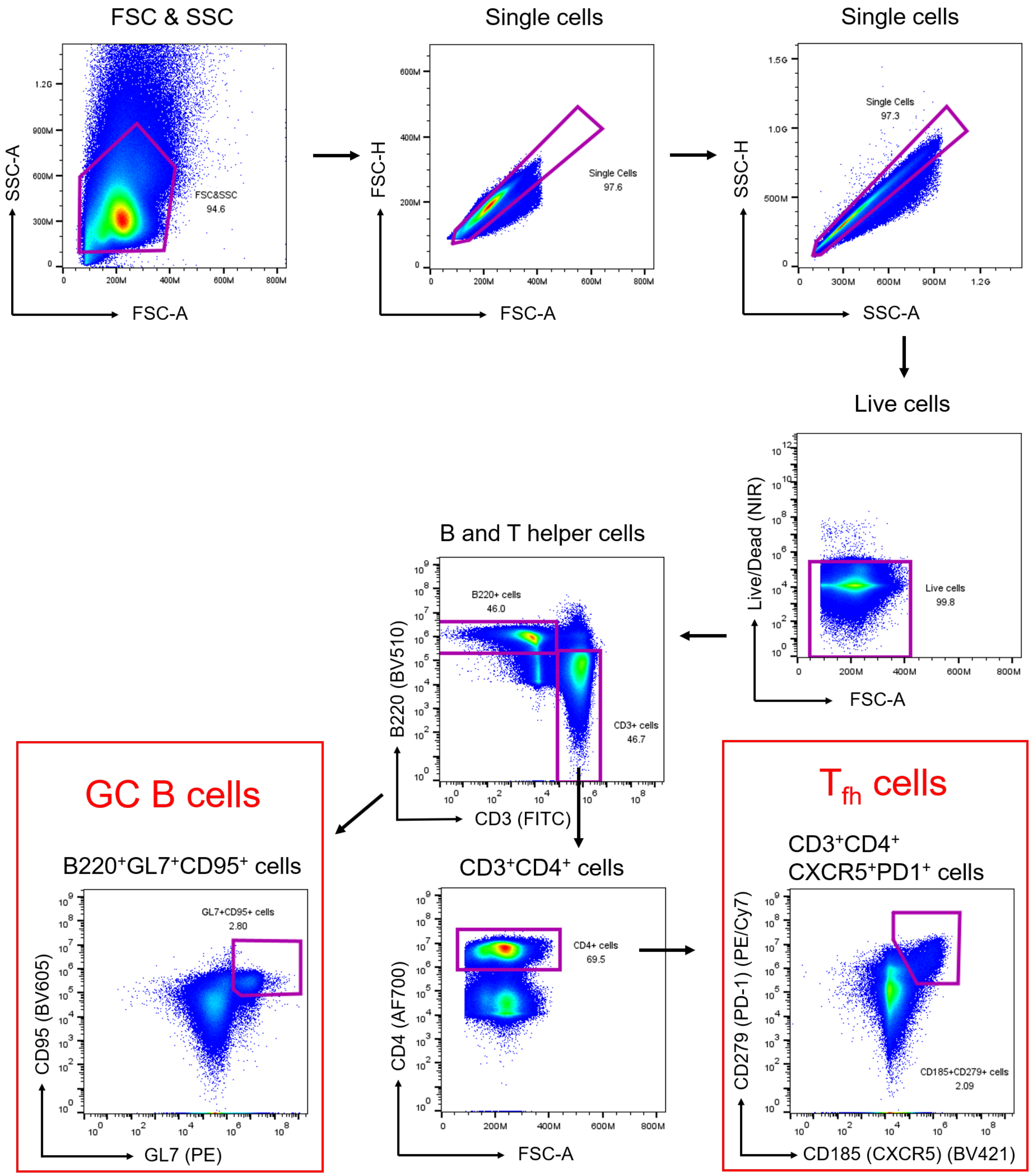


**Fig. S6. Flow cytometry analysis of germinal centers (GCs) induced by adjuvanted allergen and allergen-nanoparticles.** Gating strategy for identifying GC B cells and T follicular helper ($T_{fh}$) cells by flow cytometry (n = 5 mice per group).

**Fig. S7**

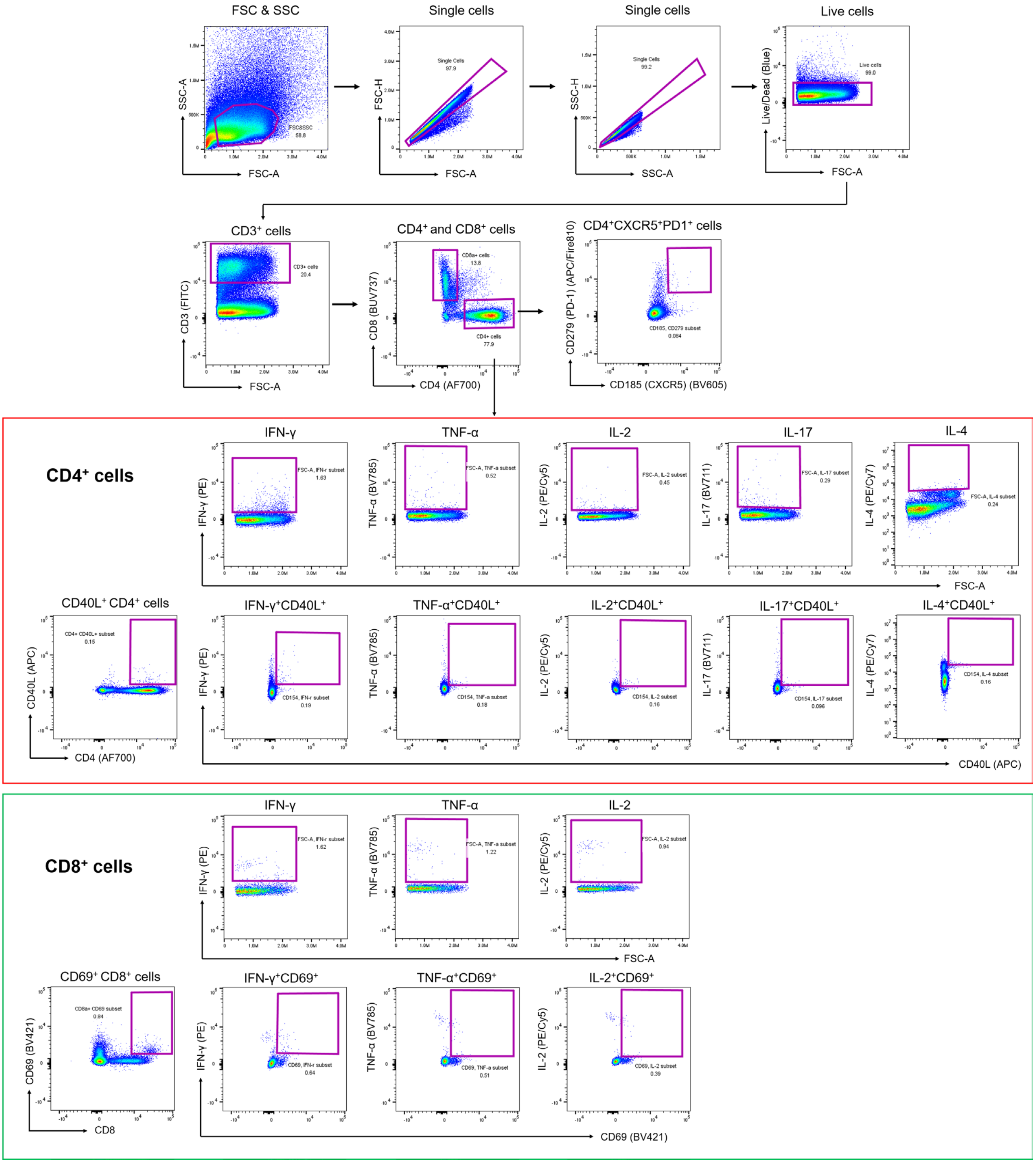


**Fig. S7. Flow cytometry analysis of Arah2-stimulated T-cell responses in adjuvanted allergen and allergen-nanoparticle groups.** Gating strategy used to quantify functional CD4+ and CD8+ T-cell responses following Arah2 allergen stimulation, assessed across adjuvanted allergen and allergen-displaying SApNP groups (n = 5 mice per group).

a

**>cf2TIP60-Q-NPE-Arah2**

MGILPSPGMPALLSLVSLLSVLLMGCVAESQDPKNIKIMRLVTGEDIIGNISESQGLITIKKAFVIIPMQATPGKPVQLVLSPWQPYTDDKEIVIDDSKV
ITITSPKDDIIKSYESHTRVLETRQMEKILALEREIERLQRQHDENLAQMNPESREELRQRRDAEIYRLEKEASGGGGSARQQWELQGDRRCQSQLERAN
LRPCEQHLMQKIQRDEDSYERDPYSPSQDPYSPSPYDRRGAGSSQHQERCCNELNEFENNQRCMCEALQQIMENQSDRLQGRQQEQQFKRELRNLPQQCG
LRAPQRCDLDVESGG

**>cf2TIP60-Q-Arah6-NPE-Arah2**

MGILPSPGMPALLSLVSLLSVLLMGCVAESQDPKNIKIMRLVTGEDIIGNISESQGLITIKKAFVIIPMQATPGKPVQLVLSPWQPYTDDKEIVIDDSKV
ITITSPKDDIIKSYESHTRVLETRQMEKILALEREIERLQRQHDENLAQMNPESREELRQRRDAEIYRLEKEASGGGGSAMRRERGRQGDSSSCERQVDG
VNLKPCEQHIMQRIMGEQEQYDSYNFGSTRSSDQQQRCCDELNEMENTQRCMCEALQQIMENQCDGLQDRQMVQHFKRELMNLPQQCNFGAPQRCDLDVS
GGRCGGGGSARQQWELQGDRRCQSQLERANLRPCEQHLMQKIQRDEDSYERDPYSPSQDPYSPSPYDRRGAGSSQHQERCCNELNEFENNQRCMCEALQQ
IMENQSDRLQGRQQEQQFKRELRNLPQQCGLRAPQRCDLDVESGG

____________: Leader sequence
____________: Arah2; __________: Arah6
Sm cfMyoX: cf2TIP60; AS: Enzymatic site; ____: Linker
**L/A**: cysteine mutation; **Q**: L117Q mutation
**N/PE**: G114N and E120P/G121E mutation
**HHHHHH**: $His_6$ tag

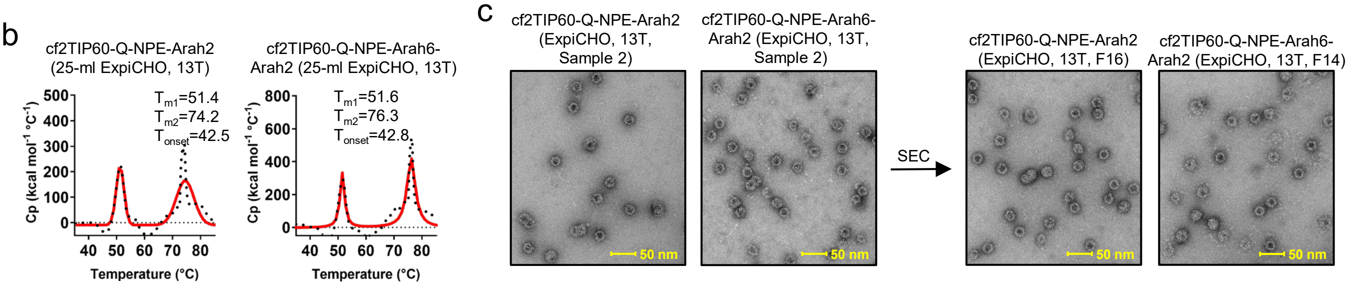


**Fig. S8. Design and evaluation of a thermostable cf2TIP60-Q-NPE SApNP scaffold for allergen display.** (**a**) Amino acid sequences of cf2TIP60-Q-NPE nanoparticles displaying Arah2 and tandem Arah6-Arah2. Signal peptide, Arah2, Arah6, Sm-like domain (within the TIP60 subunit), catfish-derived MyoX domain (within the TIP60 subunit), enzymatic site, and linkers are highlighted in yellow, brown, green, dark moss green, olive green, magenta, and gray, respectively. The cysteine in cf2-MyoX is mutated to A (red), the L117Q, G114N, and E120P/G121E mutations in cf2-MyoX are shown in hot pink. (**b**) DSC thermograms of ExpiCHO-expressed cf2TIP60-Q-NPE-Arah2 and f2TIP60-Q-NPE-Arah6-Arah2 purified by immunoaffinity chromatography (IAC) using a 13T antibody column. (**c**) Negative-stain EM micrographs of cf2TIP60-Q-NPE-Arah2 and cf2TIP60-Q-NPE-Arah6-Arah2 from an independent production run (sample 2) before (left) and after (right) SEC purification. Scale bar, 50 nm.

**Fig. S9**

a Serum IgG binding of individual female and male mice injected with allergen and allergen-nanoparticle in a prophylactic peanut allergy model

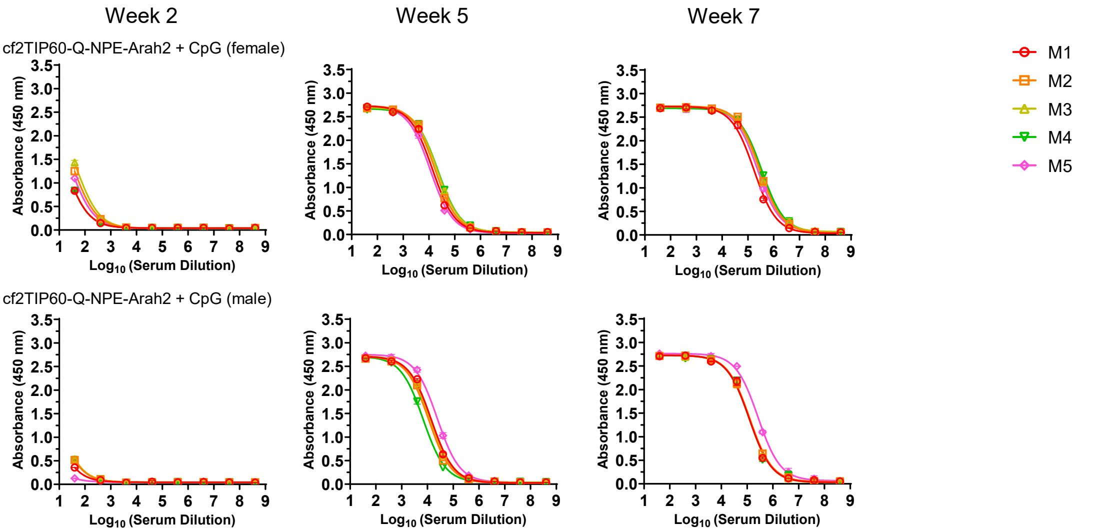


b Mouse serum IgG ELISA $EC_{50}$ titers

| Antigen | $EC_{50}$ titers (week 2) | | | | | Geometric Mean |
|---|---|---|---|---|---|---|
| | M1 | M2 | M3 | M4 | M5 | |
| cf2TIP60-Q-NPE-Arah2 + CpG (female) | 18.99 | 38.29 | 50.16 | 18.97 | 30.00 | 29.1 |
| cf2TIP60-Q-NPE-Arah2 + CpG (male) | 5.88 | 9.51 | 10.14 | 9.56 | 1.38 | 5.9 |

| Antigen | $EC_{50}$ titers (week 5) | | | | | Geometric Mean |
|---|---|---|---|---|---|---|
| | M1 | M2 | M3 | M4 | M5 | |
| cf2TIP60-Q-NPE-Arah2 + CpG (female) | 13652 | 17237 | 20919 | 22337 | 11170 | 16514 |
| cf2TIP60-Q-NPE-Arah2 + CpG (male) | 14110 | 11208 | 12682 | 6852 | 24062 | 12702 |

| Antigen | $EC_{50}$ titers (week 7) | | | | | Geometric Mean |
|---|---|---|---|---|---|---|
| | M1 | M2 | M3 | M4 | M5 | |
| cf2TIP60-Q-NPE-Arah2 + CpG (female) | 171123 | 293856 | 278777 | 345153 | 236314 | 258012 |
| cf2TIP60-Q-NPE-Arah2 + CpG (male) | 121920 | 126637 | 124084 | 121156 | 264711 | 143778 |

Statistical analysis

| Welch's t test (w2) | Statistics | P Value |
|---|---|---|
| female vs. male | * | 0.0136 |

| Welch's t test (w5) | Statistics | P Value |
|---|---|---|
| female vs. male | ns | 0.3835 |

| Welch's t test (w7) | Statistics | P Value |
|---|---|---|
| female vs. male | * | 0.0237 |

c Serum IgG binding of individual female and male mice injected with allergen and allergen-nanoparticle in a therapeutic peanut allergy model

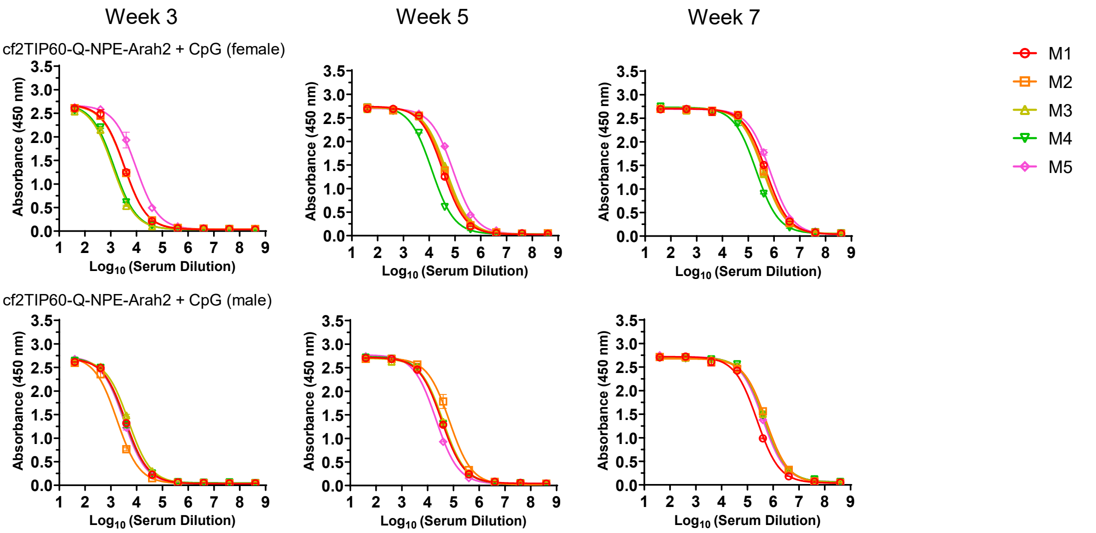

**Fig. S9**

### d Mouse serum IgG ELISA $EC_{50}$ titers

| Antigen | $EC_{50}$ titers (week 3) | | | | | Geometric Mean |
|---|---|---|---|---|---|---|
| | M1 | M2 | M3 | M4 | M5 | |
| cf2TIP60-Q-NPE-Arah2 + CpG (female) | 3450 | 3353 | 1206 | 1348 | 9597 | 2827 |
| cf2TIP60-Q-NPE-Arah2 + CpG (male) | 3769 | 1728 | 4829 | 3528 | 3163 | 3229 |

| Antigen | $EC_{50}$ titers (week 5) | | | | | Geometric Mean |
|---|---|---|---|---|---|---|
| | M1 | M2 | M3 | M4 | M5 | |
| cf2TIP60-Q-NPE-Arah2 + CpG (female) | 34076 | 40341 | 47136 | 12683 | 86163 | 37155 |
| cf2TIP60-Q-NPE-Arah2 + CpG (male) | 34857 | 70797 | 39618 | 35827 | 21158 | 37495 |

| Antigen | $EC_{50}$ titers (week 7) | | | | | Geometric Mean |
|---|---|---|---|---|---|---|
| | M1 | M2 | M3 | M4 | M5 | |
| cf2TIP60-Q-NPE-Arah2 + CpG (female) | 499864 | 410697 | 379928 | 203382 | 717118 | 408503 |
| cf2TIP60-Q-NPE-Arah2 + CpG (male) | 235544 | 533038 | 480643 | 465768 | 397377 | 407010 |

Statistical analysis

| Welch's t test (w3) | Statistics | P Value |
|---|---|---|
| female vs. male | ns | 0.8195 |

| Welch's t test (w5) | Statistics | P Value |
|---|---|---|
| female vs. male | ns | 0.81 |

| Welch's t test (w7) | Statistics | P Value |
|---|---|---|
| female vs. male | ns | 0.8472 |

### e Mouse serum IgE ELISA $OD_{450}$ values in a prophylactic peanut allergy model

| Antigen | $OD_{450}$ values (week 2) | | | | | Geometric Mean |
|---|---|---|---|---|---|---|
| | M1 | M2 | M3 | M4 | M5 | |
| cf2TIP60-Q-NPE-Arah2 + CpG (female) | 0.05815 | 0.058 | 0.0564 | 0.05895 | 0.0552 | 0.0573 |
| cf2TIP60-Q-NPE-Arah2 + CpG (male) | 0.05705 | 0.0573 | 0.0602 | 0.0599 | 0.0579 | 0.0585 |

| Antigen | $OD_{450}$ values (week 5) | | | | | Geometric Mean |
|---|---|---|---|---|---|---|
| | M1 | M2 | M3 | M4 | M5 | |
| cf2TIP60-Q-NPE-Arah2 + CpG (female) | 0.0946 | 0.0838 | 0.08045 | 0.11025 | 0.0626 | 0.0849 |
| cf2TIP60-Q-NPE-Arah2 + CpG (male) | 0.06895 | 0.0547 | 0.0648 | 0.09125 | 0.0486 | 0.0641 |

| Antigen | $OD_{450}$ values (week 7) | | | | | Geometric Mean |
|---|---|---|---|---|---|---|
| | M1 | M2 | M3 | M4 | M5 | |
| cf2TIP60-Q-NPE-Arah2 + CpG (female) | 0.0913 | 0.08935 | 0.0695 | 0.0901 | 0.0751 | 0.0826 |
| cf2TIP60-Q-NPE-Arah2 + CpG (male) | 0.0784 | 0.0655 | 0.08145 | 0.0978 | 0.05795 | 0.0750 |

Statistical analysis

| Welch's t test (w2) | Statistics | P Value |
|---|---|---|
| female vs. male | ns | 0.2665 |

| Welch's t test (w5) | Statistics | P Value |
|---|---|---|
| female vs. male | ns | 0.0914 |

| Welch's t test (w7) | Statistics | P Value |
|---|---|---|
| female vs. male | ns | 0.4325 |

### f Mouse serum IgE ELISA $OD_{450}$ values in a therapeutic peanut allergy model

| Antigen | $OD_{450}$ values (week 3) | | | | | Geometric Mean |
|---|---|---|---|---|---|---|
| | M1 | M2 | M3 | M4 | M5 | |
| cf2TIP60-Q-NPE-Arah2 + CpG (female) | 0.14365 | 0.12815 | 0.12765 | 0.10195 | 0.1342 | 0.1263 |
| cf2TIP60-Q-NPE-Arah2 + CpG (male) | 0.07645 | 0.0989 | 0.0914 | 0.12195 | 0.1067 | 0.0979 |

| Antigen | $OD_{450}$ values (week 5) | | | | | Geometric Mean |
|---|---|---|---|---|---|---|
| | M1 | M2 | M3 | M4 | M5 | |
| cf2TIP60-Q-NPE-Arah2 + CpG (female) | 0.15885 | 0.18635 | 0.20705 | 0.1066 | 0.18245 | 0.1642 |
| cf2TIP60-Q-NPE-Arah2 + CpG (male) | 0.0521 | 0.0791 | 0.07395 | 0.152 | 0.14025 | 0.0917 |

| Antigen | $OD_{450}$ values (week 7) | | | | | Geometric Mean |
|---|---|---|---|---|---|---|
| | M1 | M2 | M3 | M4 | M5 | |
| cf2TIP60-Q-NPE-Arah2 + CpG (female) | 0.1233 | 0.08455 | 0.12035 | 0.08355 | 0.0727 | 0.0947 |
| cf2TIP60-Q-NPE-Arah2 + CpG (male) | 0.05755 | 0.06375 | 0.06405 | 0.0812 | 0.08225 | 0.0690 |

Statistical analysis

| Welch's t test (w3) | Statistics | P Value |
|---|---|---|
| female vs. male | * | 0.0261 |

| Welch's t test (w5) | Statistics | P Value |
|---|---|---|
| female vs. male | * | 0.0305 |

| Welch's t test (w7) | Statistics | P Value |
|---|---|---|
| female vs. male | ns | 0.0588 |

**Fig. S9. Antibody responses of adjuvanted allergen-displaying SApNPs in female and male mice.** (**a**) ELISA IgG binding curves of female and male mouse sera from allergen-nanoparticle groups (n = 5 mice/group) in a prophylactic peanut allergy model binding to the coating antigen crude peanut extract (CPE). (**b**) (Top) Summary of geometric mean $EC_{50}$ titers measured for allergen-nanoparticle groups against CPE. Color coding indicates $EC_{50}$ levels (green to red: low to high binding). (Bottom) Summary of statistical analysis performed for each timepoint. Note: $EC_{50}$ values at week 2 were derived by setting the minimum/maximum $OD_{450}$ values to 0.0/2.5. (**c**) ELISA IgG binding curves of female and male mouse sera from allergen-nanoparticle groups (n = 5 mice/group) in a therapeutic peanut allergy model binding to the coating antigen CPE. (**d**) (Top) Summary of geometric mean $EC_{50}$ titers measured for allergen-nanoparticle groups against CPE. Color coding indicates $EC_{50}$ levels (green to red: low to high binding). (Bottom) Summary of statistical analysis performed for each timepoint. (**e**) Allergen-induced IgE responses of female and male mouse sera from allergen-nanoparticle groups in a prophylactic peanut allergy model binding to CPE. Summary of absorbance at 450 nm ($A_{450}$) values from serum IgE induced by allergen-nanoparticle groups against CPE. Color coding indicates $A_{450}$ values (green to red: low to high binding). Summary of statistical analysis performed for each timepoint. (**f**) Summary of absorbance at $A_{450}$ values from serum IgE induced by allergen-nanoparticle groups in a therapeutic peanut allergy model against CPE. Summary of statistical analysis performed for each timepoint. Error bars represent the difference between duplicate measurements at each concentration for each sample. $EC_{50}$ values were calculated using GraphPad Prism version 10.6.1. Data were analyzed using one-way ANOVA, followed by Tukey's multiple comparison post hoc test for each timepoint. Two-tailed unpaired t-tests with Welch's correction were used when comparing two independent groups. For significance, ns (not significant), $*p < 0.05$.
